# Mathematical Methodology for Dynamic Models of Insecticide Selection Assuming a Polygenic Basis of Resistance

**DOI:** 10.1101/2024.04.30.591816

**Authors:** Neil Philip Hobbs, Ian Michael Hastings

## Abstract

Mathematical models for evaluating insecticide resistance management (IRM) have primarily assumed insecticide resistance (IR) is monogenic. Modelling using a quantitative genetics framework to model polygenic IR has less frequently been used. We introduce a complex mathematical model for polygenic IR with a focus on public health insecticide deployments for vector control. Conventional polygenic models assume selection differentials are constant over the course of selection. We instead propose calculating the selection differentials dynamically depending on the level of IR and the amount of insecticide encountered. Dynamically calculating the selection differentials increases biological and operational realism, allowing for the evaluation of strategies of policy relevance, including reduced dose mixtures or the deployment of long-lasting insecticide-treated nets and indoor residual spraying in combination. The dynamic calculations of insecticide selection allow for two methods: 1) Truncation (“polytruncate”) – where only the most resistant individuals in the population survive, and 2) Probabilistic (“polysmooth”) – where an individual’s survival probability is dependent on their own level of IR. We describe in detail the calculation and calibration of these models. The models (“polytruncate” and “polysmooth”) are compared against a previous polygenic model (“polyres”) and the monogenic literature for the IRM strategies of rotations, sequences and full-dose mixtures. We demonstrate consistency in results of full-dose mixtures remaining the best IRM strategy, with sequences and rotations being similar in their efficacy between the two selection processes, and consistency in “global conclusions” with previous models. Consistency between the “polysmooth”, “polytruncate” and previous models helps provide confidence in their predictions, as operational interpretations are not overly impacted by model assumptions. This increases confidence in the application of these dynamic models to investigate more complex IRM strategies and scenarios, and their future applications will investigate more scenario specific evaluations of IRM strategies.

## 1. Introduction

Insecticides are the main tool used to control vectors of human disease including malaria, dengue and zika. Widespread deployment of insecticides as long-lasting insecticide-treated nets (LLINs) and indoor residual spray (IRS) are strong contributory to insecticide resistance (IR) (e.g. Liu, 2015). Insecticide resistance management (IRM) strategies (see Table 1 for definitions and Supplement 1 for examples) have been identified which may slow the spread of IR and mitigate any impact of IR on transmission (WHO, 2012). The imminent arrival of new insecticides for vector control, alongside next-generation mixture LLINs, means complex IRM strategies become possible.

**Table 1:** Definition and Description of Terms and Acronyms. Here we outline and define our preferred terminologies used throughout.

|  |
| --- |
| **Table 1: Definition and Description of Terms and Acronyms.** Here we outline and define our preferred terminologies used throughout. |
| **Insecticide Arsenal:** The total insecticides in the simulation; and for mixtures how those insecticides are available as mixtures. |
| **Withdrawal Threshold:** The bioassay survival which determines an individual insecticide has reached a defined “failure” point and must no longer be used. |
| **Return Threshold:** The bioassay survival which a previously “failed” insecticide must reach before the insecticide can be re-considered for deployment. |
| **Deployment Interval:** The frequency with which insecticide deployments are conducted. Note, for combinations; the LLIN and IRS can have different deployment intervals. For example, LLINs are re-distributed every 3 years, whereas IRS is re-applied every year. |
| **Insecticide Switching Strategy:** Defines the rules regarding how insecticide deployment decisions are made. |
| **Insecticide Deployment Strategy:** Defines the way in which insecticides are deployed and is one of: monotherapy, mixtures, micro-mosaics and combinations:<br>• **Monotherapy:** the deployment of a single insecticide at any time<br>• **Mixtures:** the deployment of two insecticides in a single formulation. Any mosquito encountering the mixture formulation <u>inevitably</u> contacts both insecticides.<br>• **Micro-mosaics:** Are spatial deployments at the household deployment level and involve distributing different LLINs or IRS formulations to different households in the same village.<br>• **Combinations:** Insecticides are deployed as LLINs and IRS. A single house can receive both, just the LLIN or just the IRS. Mosquitoes entering a house with both may contact both, just the LLIN or just the IRS. |
| Examples of the Deployment and Switching Strategies are provided in Supplement 1. |
| Acronyms:<br>IR = Insecticide Resistance<br>IRM = Insecticide Resistance Management<br>PRS = Polygenic Resistance Score<br>LLIN = Long-Lasting Insecticide Treated Net<br>ITN = Insecticide-treated Net<br>IRS = Indoor Residual Spray |

IRM strategies are frequently evaluated using mathematical modelling (Rex Consortium, 2007). The value of modelling is greatly improved if models: (i) cover a range of assumptions regarding the underlying biology of IR; (ii) accurately reflect the circumstances under which IR selection occurs; (iii) show consistency of results between different models or show how inconsistencies arise. Several key features which may influence the choice and effectiveness of strategies are highlighted as lacking from models, notably: insecticide dosing; insecticide decay (South et al., 2020); single insecticide exposures; cross resistance; and polygenic resistance (Rex Consortium, 2013).

The ability to include insecticide dosing in models is required for sufficient evaluation of mixtures. Reduced dose mixtures have long been highlighted as a likely compromise given the costs associated with mixtures (Curtis, 1985), and this issue is a practical reality with next-generation mixture LLINs. Insecticide decay is an important consideration for the deployment of insecticide in public health. Unlike agriculture where insecticides are replaced weekly, insecticides used in public health have long residual lifespans, replaced on a timescale of months (IRS) to years (LLINs). There are large periods where insecticide decay can occur occurs due to chemical degradation and physical degradation such as fabric damage to LLINs (e.g., Mechan et al., 2022). Insecticide decay is widely recognised but its operational impact in driving IR is unclear, and these concerns especially extend to mixtures (Curtis, 1985). A recent study by South et al (2020) attempted to quantify the effect of insecticide decay and argued, using a single-gene argument, that it could potentially rapidly increase selection for IR. Despite this to our knowledge insecticide decay is absent from models especially those evaluating IRM in a public health context.

Alternative small-scale spatial insecticide deployments (e.g., combinations or micro-mosaics (Table 1)) may allow a mosquito to encounter different insecticides throughout their lifespan. This requires extending standard modelling techniques to allow for multiple gonotrophic cycles (S. Jones et al., 2023). This work describes only the second model allowing differential exposure across gonotrophic cycles which is vital to properly evaluate the impact of combinations and micro-mosaics.

A major assumption of models evaluating IRM concerns the genetic basis of resistance. Most models assume resistance is a monogenic trait (Rex Consortium, 2010) encoded by mutations at single genes. The previous multi-gonotrophic cycle model (Jones et al, 2023) was a monogenic model so there is a need for models to check for consistency between difference genetic bases of IR. Polygenic models of IR selection (e.g., Hobbs et al., 2023) are based on the “Breeder’s”/Lush equation (Walsh & Lynch, 2018; page 482, Eq 13.1):

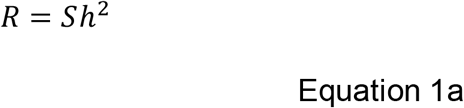

The response (*R*) is the between generation change in the mean trait value. The selection differential (*S*) is the within generation change in the mean trait value (i.e., the difference in IR level in individuals at birth and in those surviving to breed). *h*^2^ is the heritability of the trait. This equation is modified with a scaling factor (*β*) to account for uncertainty in *S* and *h*^2^ and calibrates the simulations to a defined timeframe (Hobbs et al., 2023):

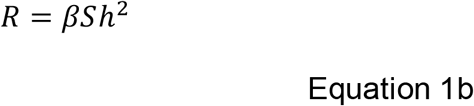

In the “polyres” model the selection differentials (for each sex) are calculated based in the level of exposure only (Equations 3a and 3b in Hobbs et al., 2023), with the parameter *β* being used to scale the simulations to a defined timeframe. This means the response to selection in “polyres” was constant throughout a simulation, regardless of the level of resistance in the population. This limits the ability of the “polyres” model to evaluate important policy and operational scenarios as the model is limited to evaluating only novel insecticides. By not being explicit in the mechanistic calculation of the selection differentials, this limited the ability to extend the model to include insecticide dosing and decay and the implication of selection at higher resistance levels. There the development of mechanistic quantitative genetics models are needed to add in the additional biological and operational realism required.

This is required as, for example, at low levels of resistance, selection is likely to be slower as most of the population is killed and the parental population is mostly individuals who avoided the insecticide, but as the insecticide decays more individuals (e.g., the more resistant individuals in the population) will survive the exposure becoming a larger portion of the parental population. Clearly, how selection occurs in the field is also unclear, and there is a need to model this as both probabilistic and truncation selection processes (Figure 1).

**Figure 1.**
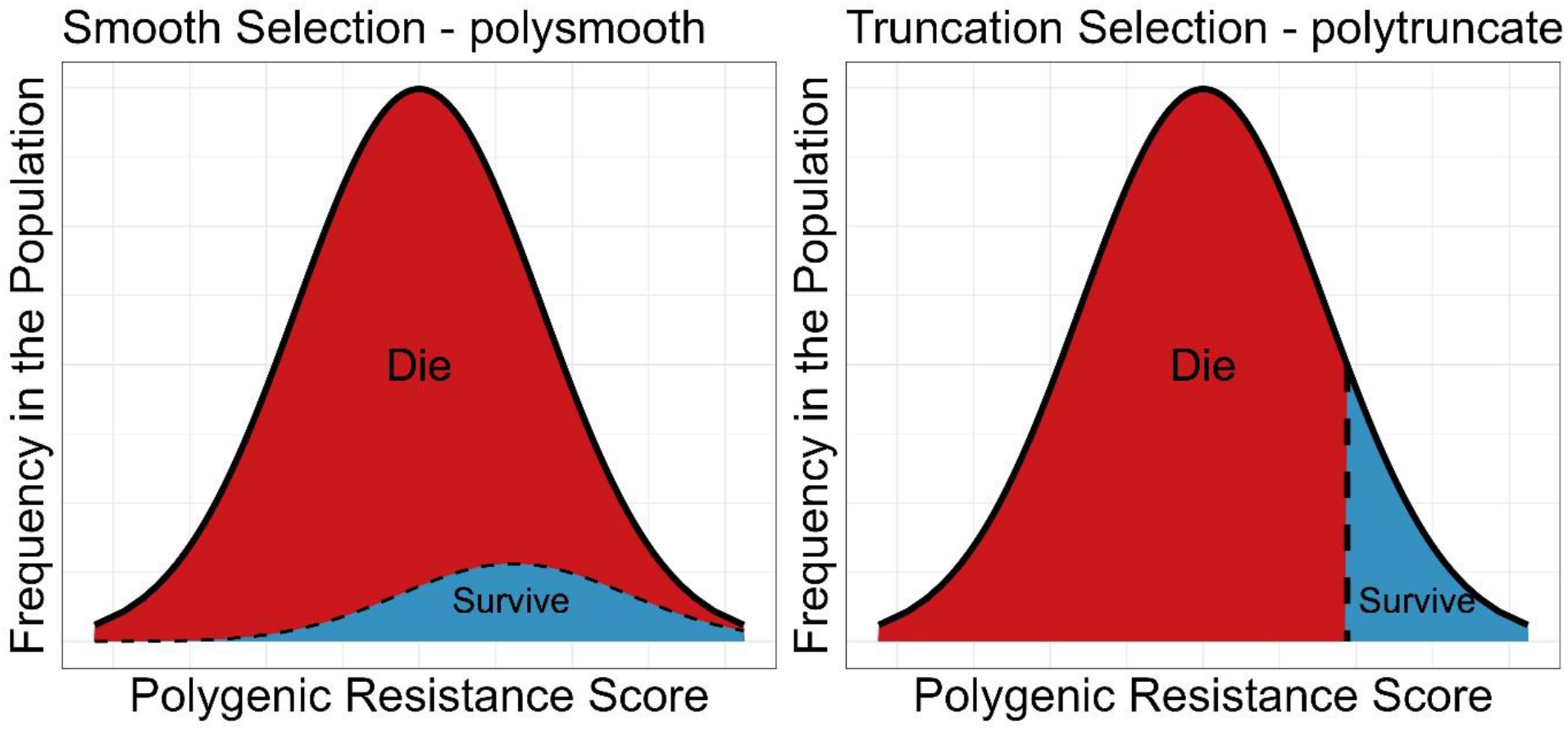
Diagrammatic Representation of Smooth and Truncation Selection. Left panel: Smooth Selection (“polysmooth”). Smooth Selection (using similar terminology to Gardner et al. 1998) defines the survival probability of an individual mosquito to be a function of its own polygenic resistance score. **Right panel: Truncation Selection (“polytruncate”)**. Truncation selection occurs when only mosquitoes with a resistance level above a certain threshold survive insecticide contact. In our model this is such that if the mean survival probability of the population was calculated as 10% to the insecticide efficacy, those individuals would be coming from the top 10% of the population. This threshold is dependent on the amount of insecticide and the degree of resistance in the population. In truncation selection, only the most resistant individuals in the population will survive the exposure. The black dashed line indicates the threshold for selection 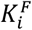 and is therefore the proportion of individuals expected to survive based on the mean PRS of the population. The individuals in red section of the distribution therefore all die, and the individuals in the blue section of the distribution all survive. The threshold PRS value at which selection occurs varies depends on the mean resistance level of the population and the current efficacy of the insecticide (unlike artificial selection where truncation selection typically always selects the top x% of the population to survive irrespective of mean population level).

Cross resistance is frequently highlighted as an important issue for the deployment of insecticides yet is lacking from most models evaluating IRM strategies (Rex Consortium, 2013). Using a previously detailed methodology (Hobbs et al., 2023) cross resistance is incorporated into the model, which to our knowledge is the first time cross resistance is implemented with insecticide dosing, insecticide decay and multiple insecticide exposures.

We present methodological extensions for a highly flexible model for the evaluation of IRM strategies which allows for questions with immediate operational implications and current policy relevance to be answered. The methodology described here is extensive but necessary to incorporate these five key features (insecticide dosing, insecticide decay, multiple insecticide exposures, polygenic resistance, and cross resistance) into a single modelling framework. Implementing these five features in a single model allows for the interactions between them to be explored. This will provide new and important insight into the conditions IRM strategies require for effective IRM.

## 2. Methods

Insecticide deployments often involve two or more insecticides, so simulations must track resistance to several insecticides simultaneously. We use lowercase letters *i, j, k* etc to refer to insecticides and uppercase letters *I, J, K* etc to refer to the corresponding resistance trait. We describe the model introducing concepts in a logical development over four methods sections:

- Methods Section 1: Quantification of Insecticide Resistance and the Inclusion of Insecticide Efficacy.
- Methods Section 2: Biology of the Response to Insecticide Selection.
- Methods Section 3: Biology of the Response to Insecticide Selection Over Multiple Gonotrophic Cycles.
- Methods Section 4: Model Calibration, and The Application to Investigating Different IRM Strategies.

The methods section requires considerable technical detail to make the methodology as flexible as possible. To aid readability Table 2 provides the key points and underlying assumptions in developing the methodology. Table 3 provides a summary of the capability our three polygenic models i.e., “polyres”, as described earlier in Hobbs et al (2023) and which assumes a constant selection differential, and the two new model branches described here i.e., “polytruncate” and “polysmooth”. Descriptions of symbols used are included in the tables in Supplement 3 in the order they appear.

**Table 2.**
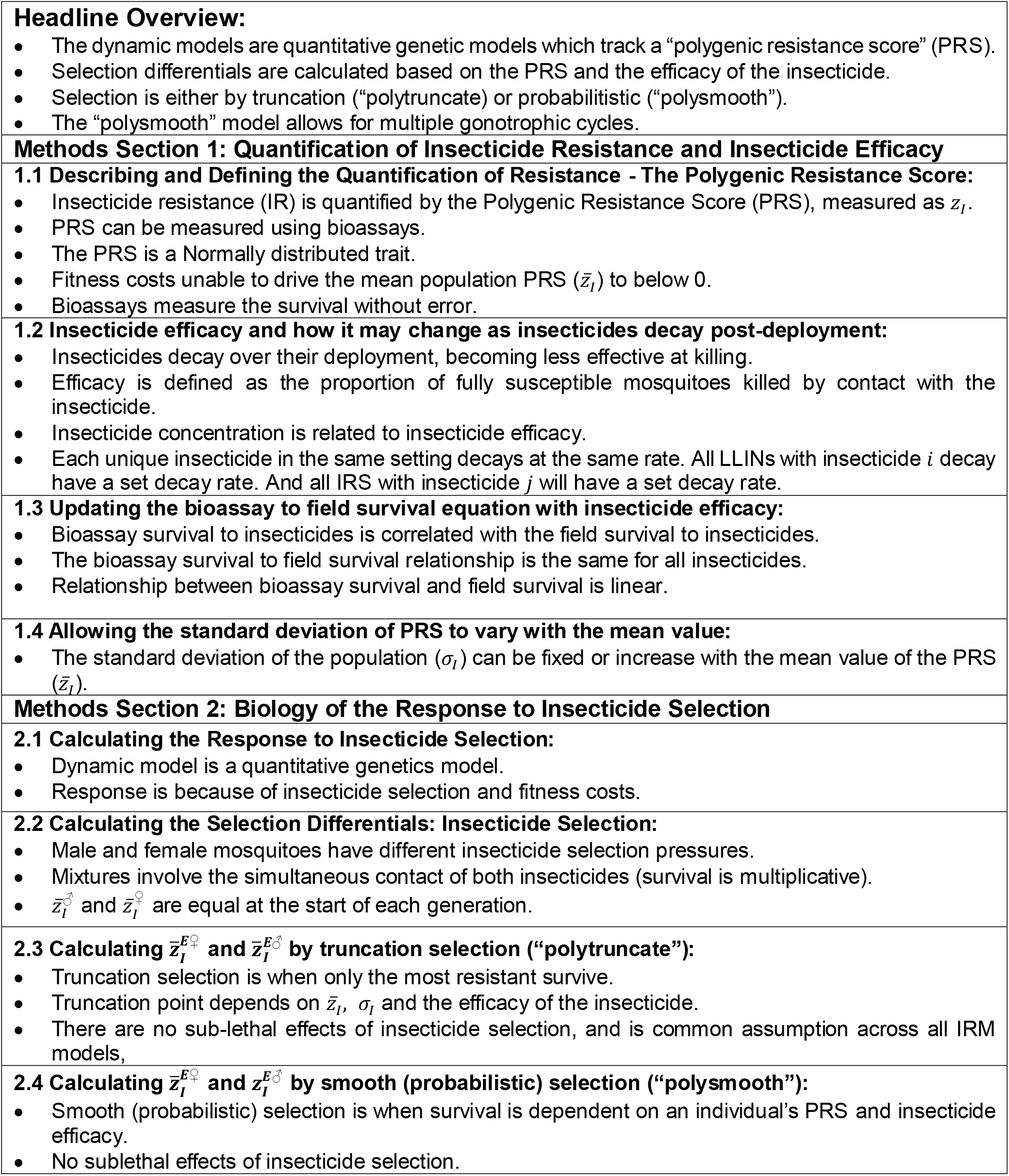
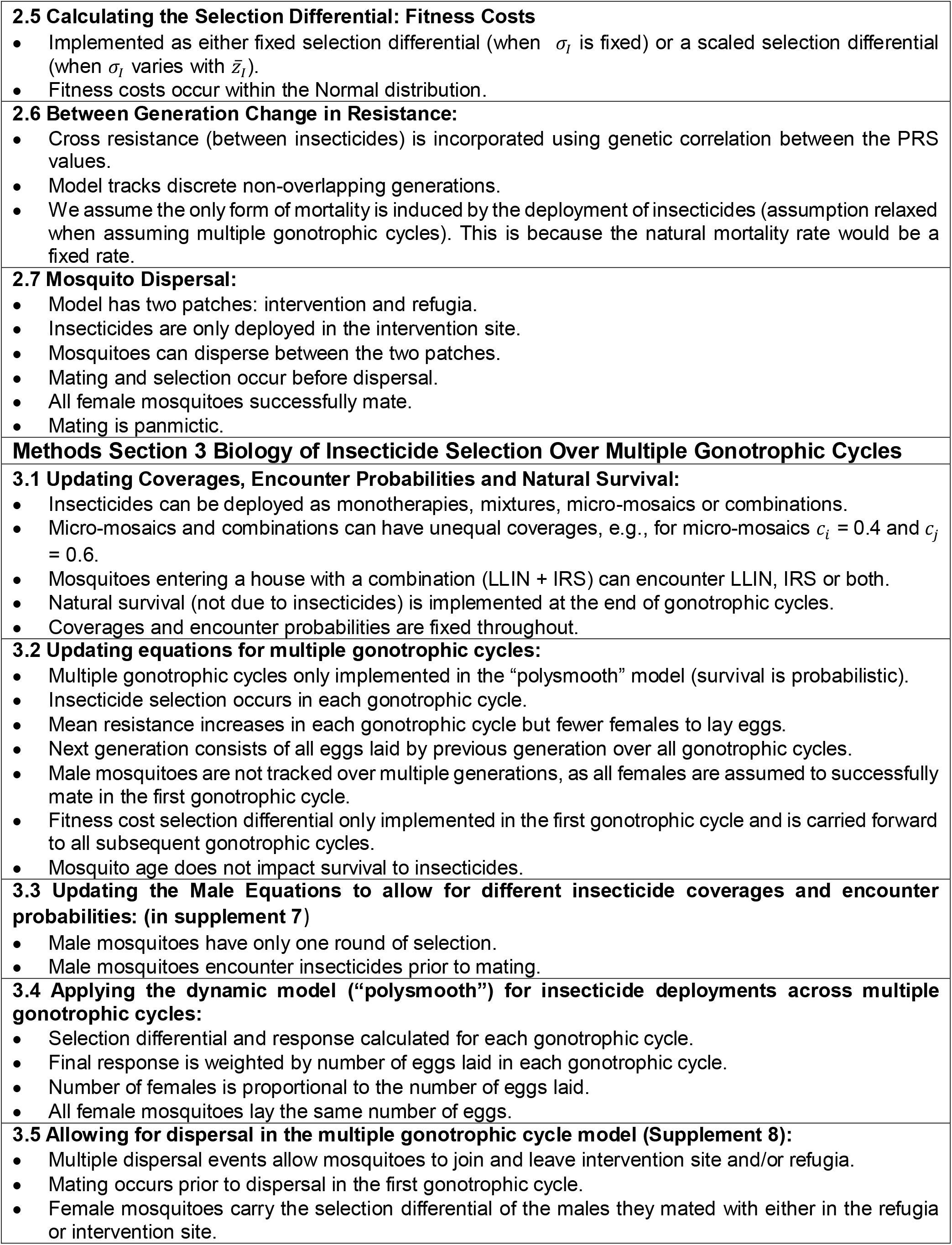
Summary of Model Methodology and key Assumptions. This table summarises and highlights the key points and assumptions for each model methodology section (except Methods Section 4).

**Table 3.** Summary of the capabilities of the dynamic model “polysmooth” and “polytruncate” compared to the previously published “polyres” model.

|  | **Polytruncate** | **Polysmooth** | **Polyres** |
| --- | --- | --- | --- |
| **Insecticide Deployment Options** |  |  |  |
| Monotherapies | Yes | Yes | Yes |
| Mixtures | Yes | Yes | Yes |
| Combinations | No | Yes | No |
| Micro-Mosaics | No | Yes | No |
| Number of Insecticides in Arsenal | Technically Unlimited | Technically Unlimited | Technically unlimited |
| **Parameter Variables** |  |  |  |
| Cross Resistance | Yes, as correlated responses. | Yes, as correlated responses. | Yes, as correlated responses. |
| Insecticide Dosing | Variable (unique per insecticide) | Variable (unique per insecticide) | Fixed, full-concentration only. |
| Insecticide Decay | Yes (unique per insecticide) | Yes (unique per insecticide) | No |
| Mechanism of Selection | Truncation | Probabilistic | Not defined |
| Multiple gonotrophic cycles | No | Yes (with natural mortality between gonotrophic cycles) | No |
| Linkage disequilibrium. | Does not explicitly account for linkage disequilibrium | Does not explicitly account for linkage disequilibrium | Does not explicitly account for linkage disequilibrium |

The mathematical models are written in R. Model code is available from the github repositories: https://github.com/NeilHobbs/polytruncate and https://github.com/NeilHobbs/polysmooth.

## Methods Section 1: Quantification of Insecticide Resistance and Inclusion of Insecticide Efficacy

### Methods Section 1.1: Describing and Defining the Quantification of Resistance - The Polygenic Resistance Score

We previously described a “Polygenic Resistance Score” (PRS), (i.e., in the “polyres” methodology in Hobbs et al., 2023) to quantify the level of insecticide resistance, and use this scale again in this manuscript. The underlying rationale and mathematics are briefly recapped in Equations 2a and 2b in Supplement 4. To summarise, the PRS is a quantitative trait of polygenic IR, with a mean value denoted 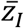 where *I* indicates *z* is measuring PRS to insecticide *i*. The PRS is measurable in bioassays (e.g., W HO tube tests). We calibrate the model such that a value of *z*_*I*_=0 indicates resistance is absent and a value of *z*_*I*_=100 gives 10% bioassay survival to insecticide *i*, a threshold commonly used to a define a “failing” insecticide. Mosquitoes with a negative polygenic resistance score are modelled though they have a score of 0. That is, they will be reliably killed even if they encounter very concentrations of insecticide. There is an issue in having a “more than susceptible” reference population. For example, the bioassay survival to field survival relationship was modelled using paired experimental hut and bioassay survival. Where, if the bioassay survival is zero, then it is not possible to know the “true” PRS value of the population, as all mosquitoes with PRS values of zero or below would not survive. This also prevents any single PRS value giving a negative survival value. For the truncation model, this assumption does not matter, as those individuals would not survive regardless. For the probabilistic selection, this does slow down the initial rate of selection, until less of the distribution is below 0, after which resistance “takes off”.

### Methods Section 1.2: Insecticide efficacy and how it may change as insecticides decay post-deployment

Insecticides and insecticidal products decay over time, a worrying process in control programmes that results in reduced insecticide effectiveness and, in all probability, a change in the intensity of selection for resistance. Insecticide decay is routinely absent from models of IR evolution; see discussion with examples in (South et al., 2020).

Ideally insecticides are deployed at the manufacturers’ specified concentration after which insecticide concentrations are likely to decay exponentially after application:

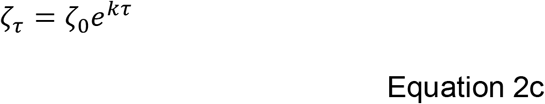

Insecticides are deployed at an initial concentration (*ζ*_0_) at time zero, decaying at rate *k* each mosquito generation (*τ*) to give the insecticide concentration (*ζ*_*τ*_) at *τ* generations post deployment. Insecticide concentration must be converted to efficacy in terms of killing the key insect target species. We define insecticide efficacy as the ability of the insecticide to kill fully susceptible mosquitoes (*z*_*i*_ ≤ 0). Therefore, insecticides deployed at the manufacturer specified concentration such as a brand new single-insecticide LLIN have an initial insecticide efficacy 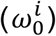 of 1.

Insecticide decay depends on both insecticide chemistry and environmental conditions (Mutagahywa et al., 2015). Decay may be slow initially, becoming rapid after the intended longevity of the product is exceeded. Figure 4 shows an example insecticide decay profile. In stage 1 (blue part of Figure 4) the insecticide is newly deployed, and the decay rate is low. Insecticide efficacy at time *τ* after deployment is calculated as follows:

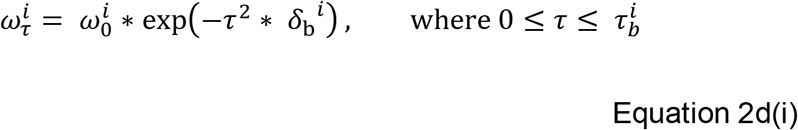

Where 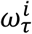 is the efficacy of insecticide *i* at time *τ* since deployment, in mosquito generations. *δ*_b_^*i*^ is the basal decay rate of insecticide *i*. 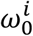 is efficacy of insecticide *i* at *τ*=0.

In stage 2 (red part of Figure 4), the insecticide has been deployed beyond a longevity threshold, 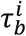. After this threshold there is a change in the decay rate and insecticide efficacy is calculated:

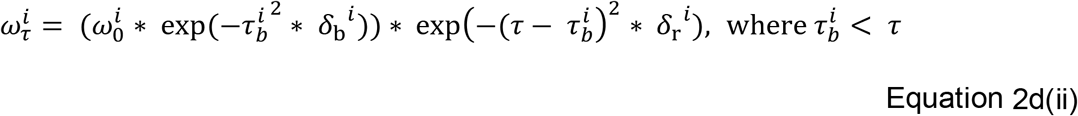

Where 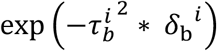 is the insecticide decay occurring before the rapid degradation, and 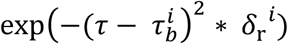 is the rapid decay occurring after the longevity threshold has been exceeded which occurs at time 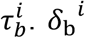 is the efficacy decay rate during the *τ*_b_ generations prior to the longevity threshold and *δ*_r_^*i*^ is rate of efficacy decay after the longevity threshold at time 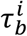.

The initial efficacy of the deployed insecticide can be set by specifying the initial value Of 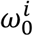. This therefore allows for the inclusion of over 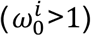 and under 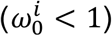 spraying which can occur during IRS implementation. This concept can also be applied to mixtures where each insecticide may not be deployed at the manufacturer specific monotherapy dose.

The use of a two-step decay process aims to allow for the model to be as flexible as possible with regards to its ability to evaluate different insecticide decay profiles. The decay profiles this two-step process produces aims to be a composite of both insecticide concentration and physical damage with nets. That is the efficacy of the LLIN (its ability to kill) is a compositite of both the insecticide concentration and the physical net integrity. The insecticidal capacity of LLINs (for example as measured in Toé et al., 2019) does not additionally account for damage to the net (and therefore the probability of reduced contact times). By including this two-step decay process it is therefore possible to capture the impact of reduced efficacy as a result of a decay in the efficacy of the insecticide, the physical decay of the net and the throwing away of nets which are deemed no longer effective (Martin et al., 2024). The flexibility of the model does of course allow for only including a single linear decay, by setting it such that the threshold generation decay value is the same time frame as deployments. Sudden changes in the insecticide efficacy of an IRS could be due to painting or plastering over sprayed walls.

### Methods Section 1.3. Updating the bioassay to field survival equation with insecticide efficacy

Equation 2b (Supplement 4) describes the relationship between bioassay survival 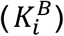 and field survival 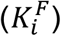 as estimated by a linear model (Hobbs et al., 2023). This relationship is updated to account for insecticide efficacy 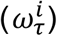.

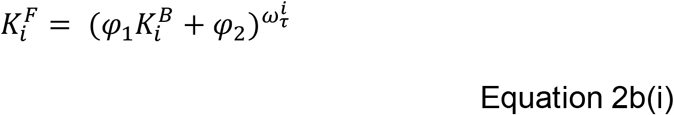

Where *φ*_1_ and *φ*_2_ are the regression coefficient and intercept of the linear model respectively. The value of 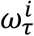 is calculated above in equations 2d(i) and 2d(ii). W hen 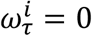, the insecticide has decayed to an ineffective concentration and survival is 100%. The relationships between PRS, efficacy and field survival are demonstrated in Figure 5.

### Methods Section 1.4: Allowing the standard deviation of PRS to vary with the mean PRS value

An important question when considering a Normally distributed trait, such as the PRS, is the value of its phenotypic standard deviation (*σ*_*I*_). There are two distinct scenarios. First, *σ*_*I*_ remains constant over the course of IR selection. Second, *σ*_*I*_ changes with respect to the current 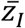 (for example the coefficient of variation SD/mean may remain constant such that the arithmetic value of the standard deviation depends on the mean) this scenario will plausibly become necessary when considering insecticides when deployed against mosquito populations with high levels of IR. These two options are detailed in Supplement 5 (Equation 2e). The choice of which option is used depends on the scenario being evaluated.

## Methods Section 2: Biology of the Response to Insecticide Selection

### Methods Section 2.1: Calculating the Response to Insecticide Selection

The deployment of insecticides leads to selection which drives an increasing 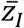 over successive generations. Conversely, fitness costs may reduce 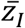 over successive generations. The overall selection differential 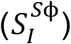 is therefore a result of insecticide selection 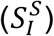 and fitness costs 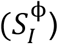. As male and female mosquitoes often have different behaviours and physiologies the selection differentials are calculated separately. Equations are suffixed *♂* (male) and *♀* (female).

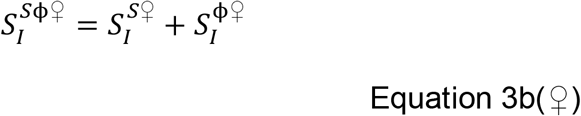

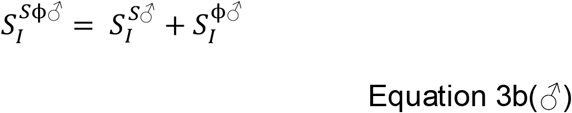

The overall selection differentials are then used in the sex-specific Breeder’s equation (Walsh & Lynch, 2018; page 485, Eq 13.5) to calculate the response:

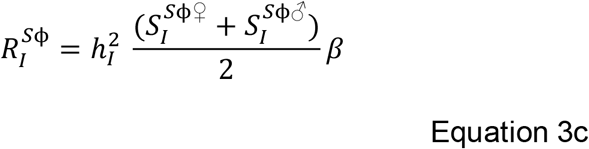

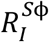 is the response in trait *I* (i.e., resistance to insecticide *i*), 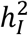 is the heritability of trait *I. β* is the exposure scaling factor and is used to calibrate simulations to defined timescales. When the insecticide is not present, the sex-specific Breeder’s equation includes only fitness cost selection differentials, and the response is therefore 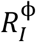. This occurs in the refugia, or in the intervention site when the insecticide is not deployed.

### Methods Section 2.2 Calculating the Selection Differentials: Insecticide Selection

Where this model differs from previous models of polygenic IR is that the insecticide selection differential 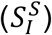 is calculated dynamically based on both the level of IR and insecticide efficacy (“polytruncate”: Figure 2 panel 5&6 and “polysmooth”:Figure 3 panel 5&6), rather than being assumed constant as previously implemented (Hobbs et al., 2023). Insecticide selection differentials are calculated separately for males and females:

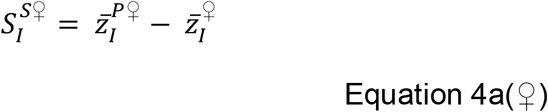

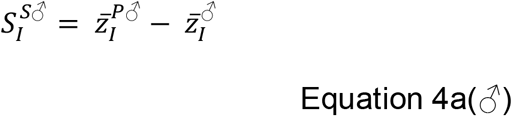

**Figure 2:**
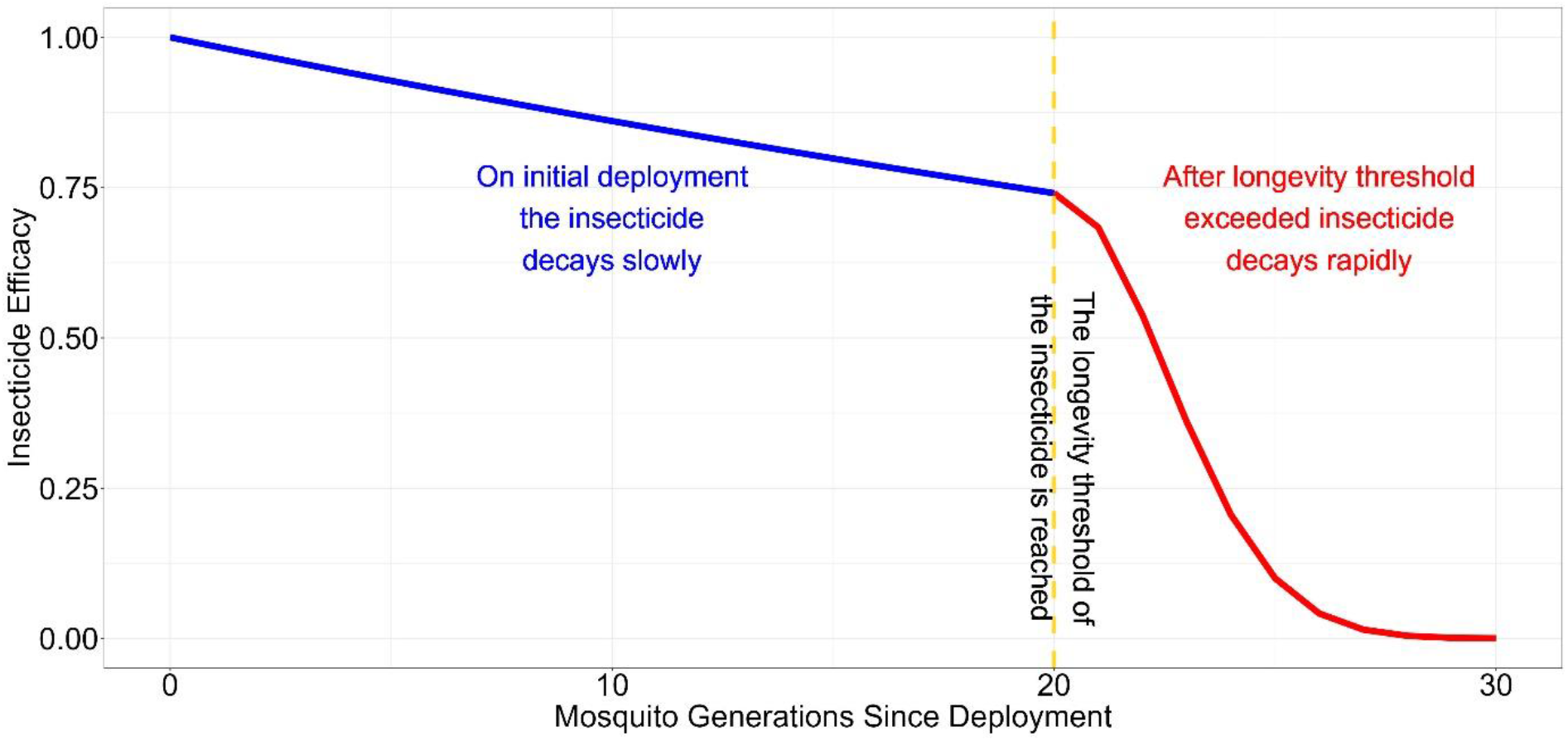
An Example Insecticide Decay Profile For an LLIN. In this example the insecticide is deployed at its manufacturers recommended concentration with an initial deployed efficacy of the insecticide 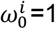. During the first two years (∼20 mosquito generations), the insecticide decays slowly at its base decay rate (blue background). After the two years (∼20 generations) the decay threshold has been reached (yellow dashed line), the insecticide decays rapidly (red background). In this example the base decay rate *δ*_b_ ^*i*^ was 0.015, the rapid decay rate *δ*_r_ ^*i*^ was 0.08 and the decay threshold 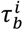 was 20 generations (∼2 years). These values are user-defined inputs and can be varied allowing insecticides to decay faster or slower. Setting *δ*_b_ ^*i*^ and *δ*_r_ ^*i*^ to 0 prevents insecticide decay from occurring (a key assumption in previous models). Insecticides deployed in mixture can have different properties such that the rapid decay rate, longevity threshold and rapid decay rates are not the same.

**Figure 3.**
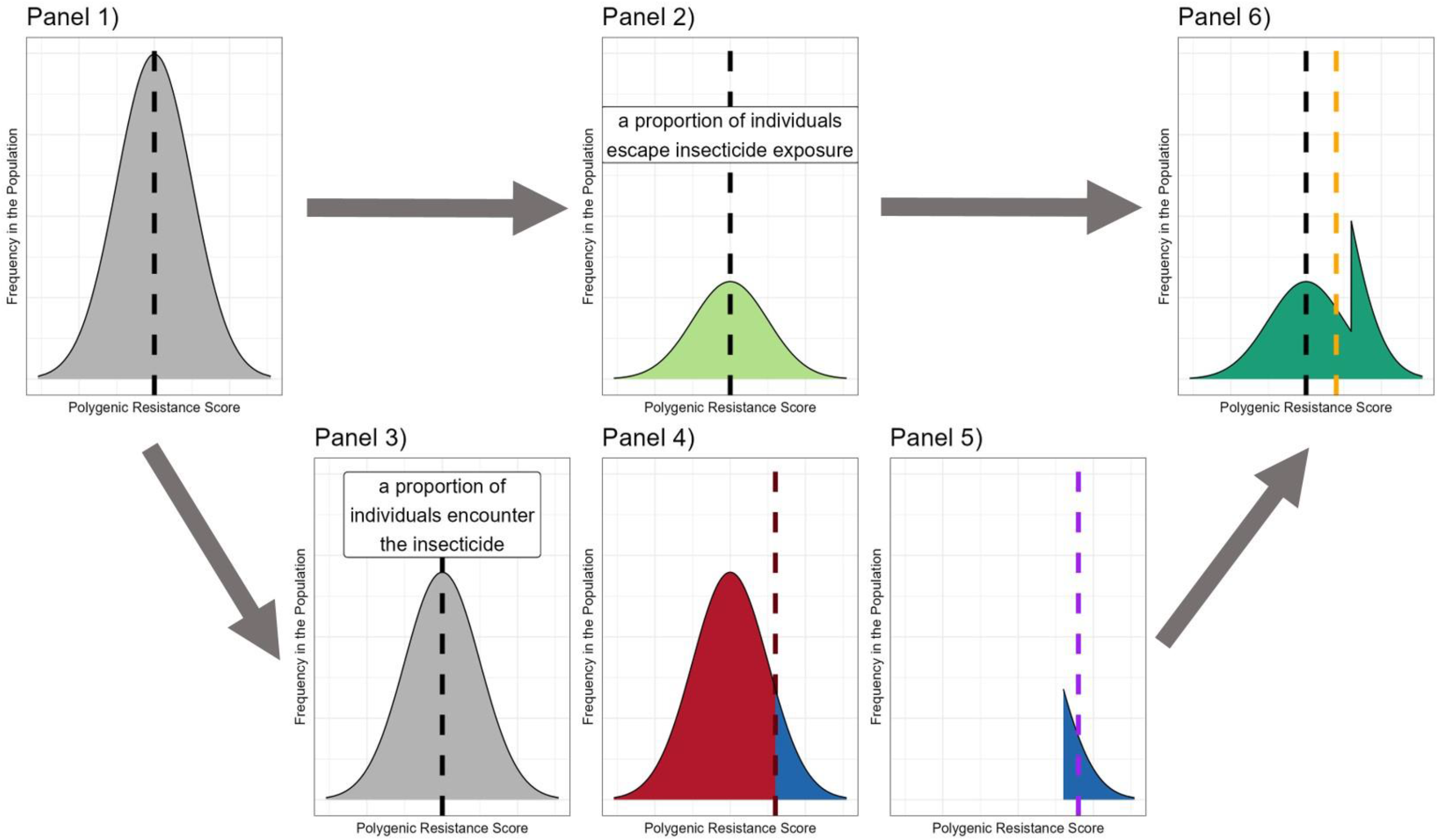
Compartmental Diagram of Truncation Selection Process for a Single Generation. This figure represents a diagrammatic representation of the model when implementing truncation selection (“polytruncate”). It shows the process considering the whole population but note that this process is done separately for males and females. **Panel 1**. The mosquito population emerges, with a PRS that is Normally distributed, and has a mean of 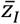 and standard deviation *σ*_*i*_. At emergence there are a total of *N*^*T*^ individuals. **Panel 2**. A proportion (males:1 − *mx* and females:1 − *x*) of the mosquitoes avoid the insecticide and therefore avoids insecticide selection; the mean of this group is therefore unchanged as 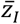, there will be *N*^*u*^ number of these individuals (Equation 4d). ***Panel 3***. A proportion of the mosquitoes do encounter the insecticide (males: *mx* and females: *x*) (Equation 5d). ***Panel 4***. These individuals are selected by truncation selection (Equation 5c) where individuals with a PRS less than the defined threshold (dark red dashed line) die (red), and the individuals with a PRS above the threshold survive (blue). ***Panel 5***. Only the most resistant individuals in the population will have survived the insecticide encounter, these individuals have a new updated mean of 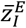 (purple dotted line) (Equation 5c)), and there are 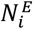 individuals surviving the insecticide selection (Equation 5d). ***Panel 6***. The unexposed group (Panel 2) and the exposed survivors (Panel 5) form the final breeding parental population. They have a mean of 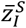 (orange dotted line) (Equation 4b)), which would be expected to be higher than the original population mean (black dotted line) to create the insecticide selection differential (which would be calculated as orange dotted line minus the black dotted line (Equation 3b).

**Figure 4.**
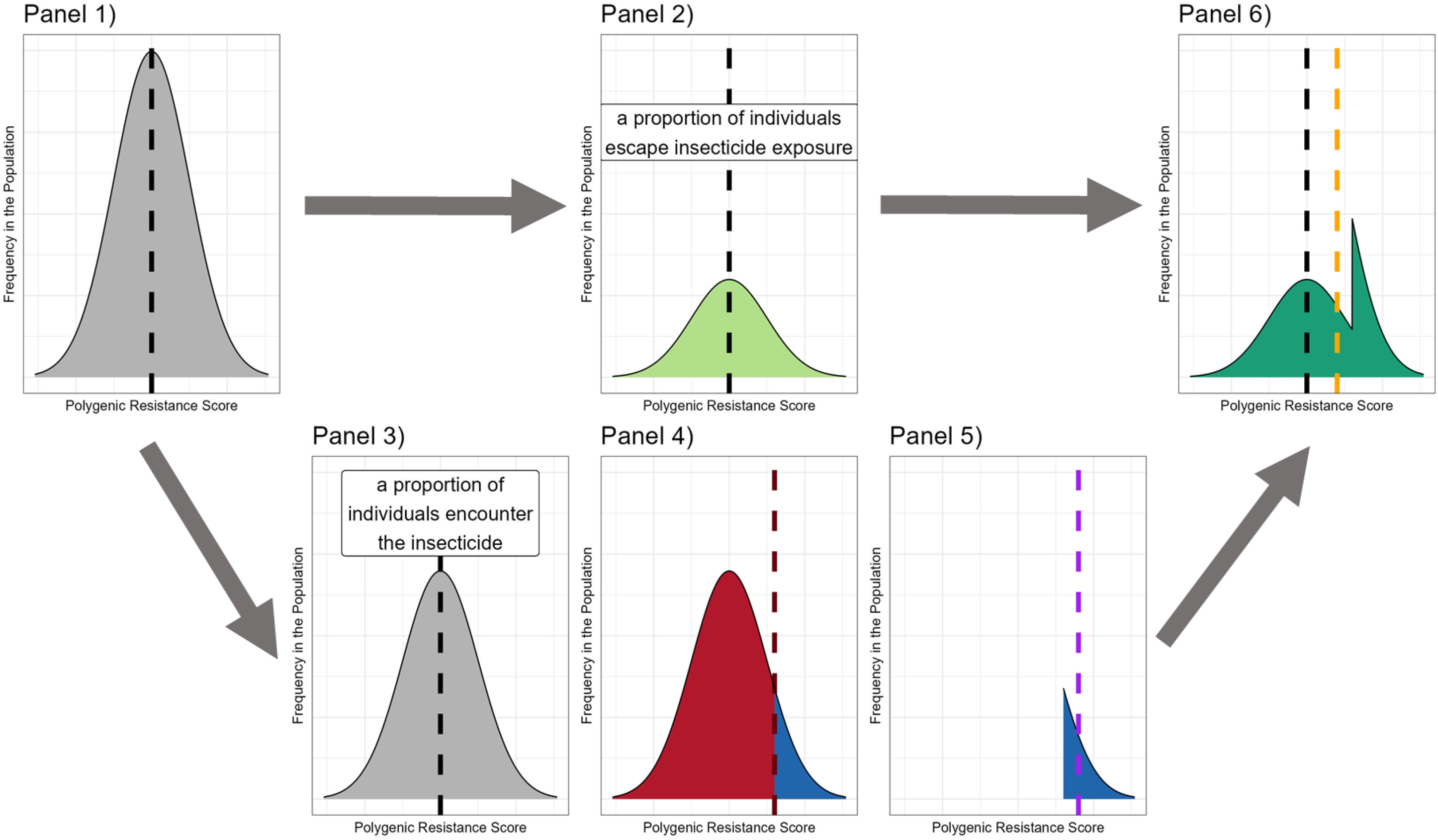
Compartmental Diagram of the Smooth Selection Process for a Single Generation. Panels 1-3 and 6 have same description as Figure 2. Where the two models diverge is the calculation of who survives/dies insecticide exposure. **Panel 4:** The survival of the insecticide encounter will be dependent on an individual’s polygenic resistance score, individuals with a higher polygenic resistance score will be more likely to survive. **Panel 5**. The mosquitoes surviving exposure will have a higher mean polygenic resistance score (purple dotted line (Equation 6c)) and there will be and there are 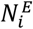 individuals (Equation 6b).

**Figure 5.**
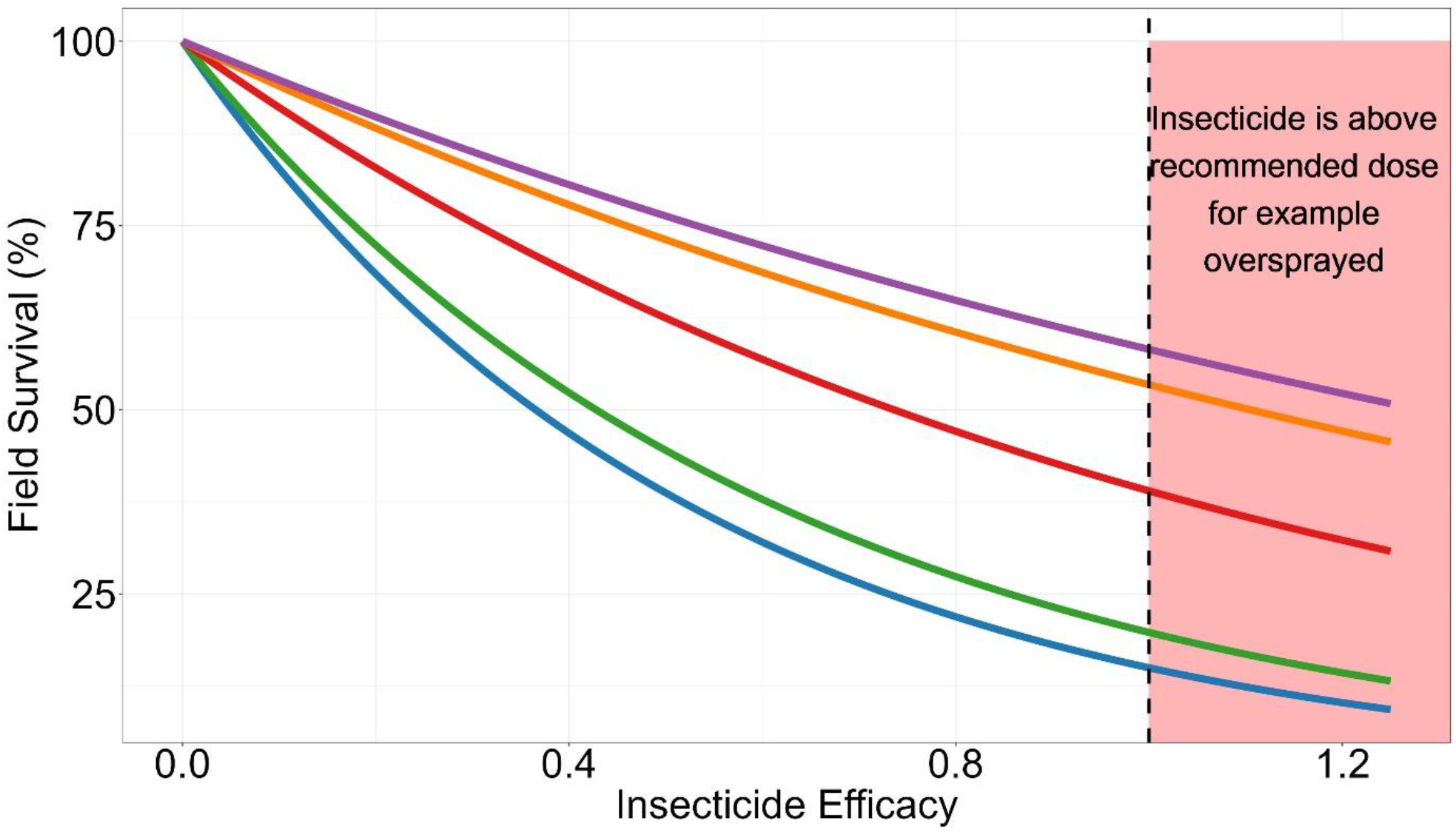
Impact of Insecticide Efficacy on Field Survival at Given Polygenic Resistance Scores. This plot displays the field survival of mosquitoes which encounter an insecticide of a particular efficacy, calculated using Equation 1b(i). The colour of the line corresponds to the polygenic resistance score of the mosquito. Blue = 0, Green = 10, Red = 100, Orange = 900, Purple = 8100; these correspond to 0, 2, 10, 50 and 80% bioassay survivals respectively. The black dashed line indicates the insecticide is at its manufacturers recommended concentration for 100% efficacy, which is 1. The red background indicates the insecticide is above the manufactures recommended dosing/concentration as may occur, for example, with over-spraying when performing indoor residual spraying.

Where 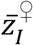 and 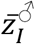 are the mean PRS of female and male mosquitoes when hatching prior to insecticide selection (we assume both sexes have the same level of IR at birth i.e.,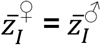). The values 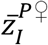 and 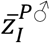 are the mean values of the final breeding parental population (“polytruncate”: Figure 2 Panel 6 and “polysmooth”: Figure 3 Panel 6). The “*P*” superscript indicates the final breeding parental population.

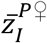 and 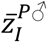 are calculated from those surviving insecticide exposure (“polytruncate”: Figure 2 Panel 5 and “polysmooth”: Figure 3 Panel 5)) and those which avoided the insecticide (“polytruncate”: Figure 2 panel 2 and “polysmooth”: Figure 3 panel 2):

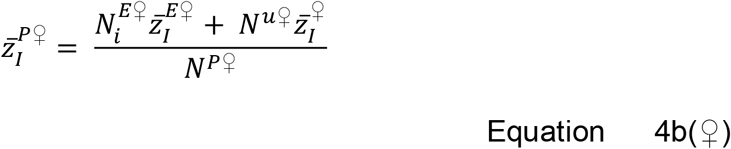

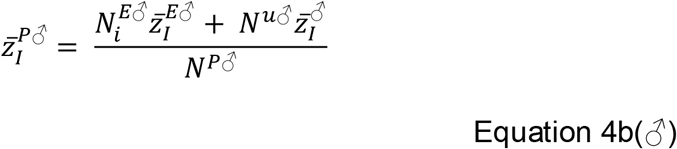

The number of mosquitoes surviving the insecticide exposure is 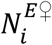 and 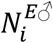, and these individuals are expected to have a higher mean PRS of 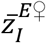 and 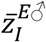 (“polytruncate”: Figure 2 Panel 5 and “polysmooth”: Figure 3 Panel 5). The “*E*” superscript indicates they underwent insecticide exposure.

The number of mosquitoes which avoid the insecticide(s) and survive to become parents is *N*^*u♀*^ and *N*^*u♂*^. The “*u*” superscript indicates they were not exposed to the insecticide(s) (Figure 2 Panel 2 and Figure 3 Panel 2) and by not having insecticide selection these individuals have an unchanged mean PRS of 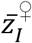 and 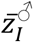.

*N*^*P♀*^ and *N*^*P♂*^ (Figure 2 Panel 6 and Figure 3 Panel 6) are the total number of breeding parents consisting of those surviving exposure (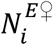 and 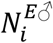) and those avoiding exposure (*N*^*u♀*^ and *N*^*u♂*^):

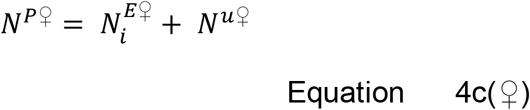

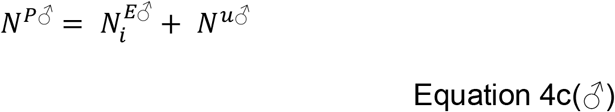

*N*^*u♀*^ and *N*^*u♂*^ are the number of mosquitoes (female and male) avoiding the insecticide encounter (Figure 2 Panel 2 and Figure 3 Panel 2) and surviving to become breeding parents:

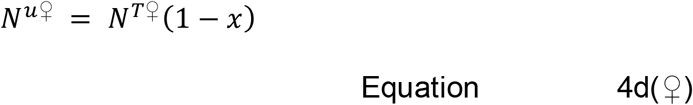

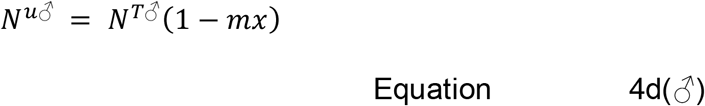

Equation 4d(*♂*) Where *x* is the proportion exposed to insecticide and *m* is the male exposure as proportion of female exposure (generally *m*<1 as males behaviour make them less likely to encounter insecticides).

The important question is how to calculate the mean resistance of the exposed survivors (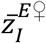 and 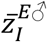) to be used in Equations 4b. This can be calculated directly as is the case for (probabilistic) selection. Or 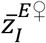 and 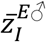 can be calculated by first calculating the selection differentials between the exposed survivors and the mean population (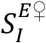 and 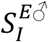) and using this to calculate 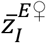 and 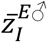:

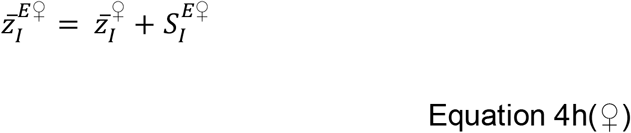

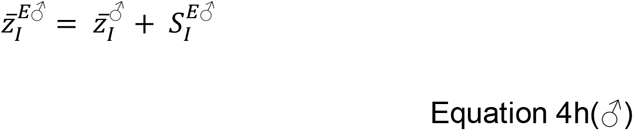

For truncation selection, the values of 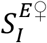 and 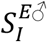 are calculated to subsequently calculate 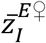 and 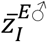.

We consider two selection processes (Figure 1), truncation selection (“polytruncate”) and smooth (probabilistic) selection (“polysmooth”) whose conceptual descriptions of the whole selection process are given in Figures 2 and 3 respectively.

### Methods Section 2.3: Calculating 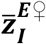 and 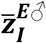 by truncation selection (“polytruncate”)

In truncation selection (**Error! Reference source not found**.1 and Figure 2) only the most resistant individuals survive insecticide exposure. Truncation selection is implemented using Truncation selection is implemented using the equation for the intensity of selection (Walsh & Lynch, 2018, equation 14.3a, page 508):

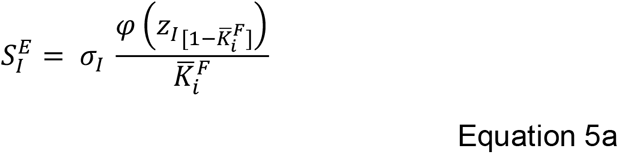

The value 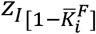 is the value of the PRS which would mean that the average survival rate of the population is 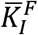. The value *σ*_*I*_ is the standard deviation. *φ*(*z*_*I*_) is the unit Normal density distribution function:

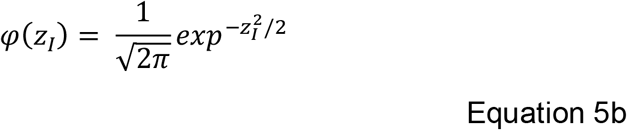

In the model code this is implemented as dnorm(qnorm 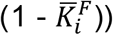.

The mean PRS of the surviving exposed mosquitoes 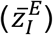 is calculated as sex-specific equations:

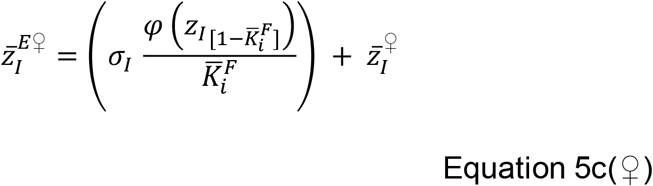

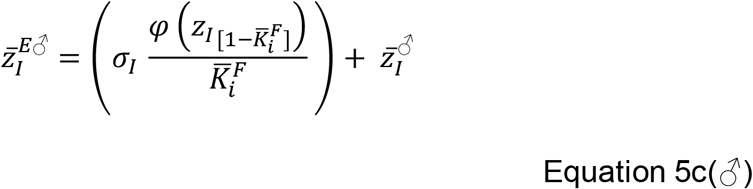

The number of exposed surviving females and males is the number exposed (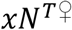 for females, and 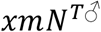 for males) multiplied by proportion surviving 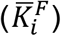:

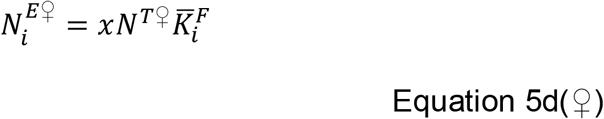

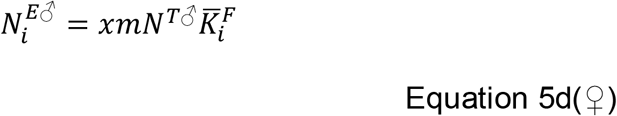

Extending for mixtures, where mosquitoes must simultaneously survive the encounter to both insecticides:

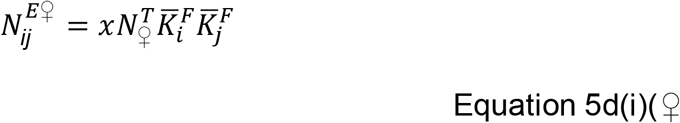

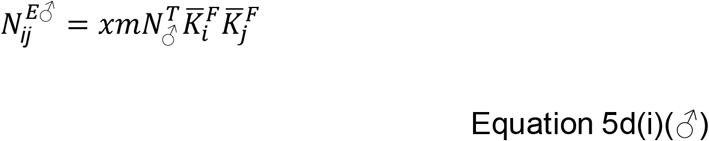

Where 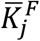 is the mean survival to insecticide *j* and is calculated from Equation 2b(i) and is dependent on 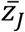 and the efficacy of insecticide *j*.

The values of 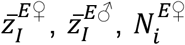 and 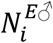 are returned to Equations 4b. If a mixture was deployed 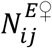 and 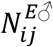 replaces 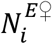 and 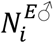 in Equations 4b. The solving of Equations 4b gives 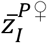 and 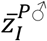, which is required to calculate the insecticide selection differential (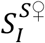 and 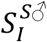) which can be combined with any fitness costs to give the overall selection differentials which are used in the sex-specific Breeder’s Equations (Equation 3c) to calculate the overall response which can be passed through to Equation 8 to give the mean of the next generation of mosquitoes allowing the cycle to continue.

### Methods Section 2.4: Calculating 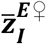 and 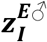 by smooth (probabilistic) selection (“polysmooth”)

In smooth selection, survival is modelled as a probabilistic process (see Figures 1 and 3). The PRS is a continuous, quantitative trait, so the values of *z*_*I*_ are “binned” into discrete classes with a corresponding frequency 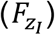 (Supplement 3, Figure S1). For each sex and resistance trait the model code tracks two vectors, one which contain the discrete “binned” values of *z*_*I*_ and a corresponding vector of containing the corresponding frequencies 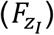 Tracking these changes in the distributions of the “binned” values allows for simple computational implementation.

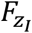 is the frequency of individuals with PRS *z*_*I*_ and is calculated using the unit Normal density distribution function:

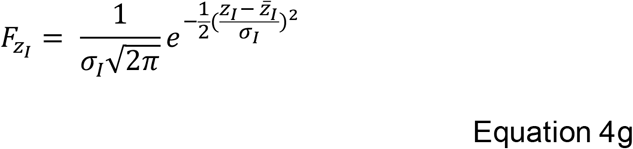

Therefore, for females 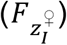 and males 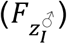 are each half of 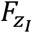 at hatching.

The total population size prior to insecticide selection is therefore the sum all the frequencies of each binned value of *z*_*I*_ in the population. This enable total population size to be calculated as

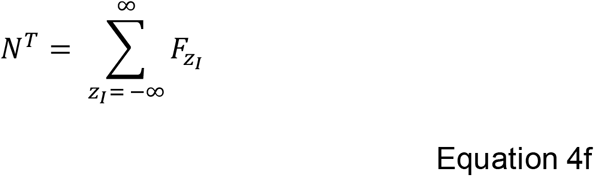

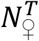 and 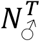 are calculated as half of the total population size before any selection has occurred, assuming the ratio at hatching of males to females is 1:1.

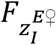 and 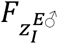 are the frequency of bins of 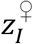 and 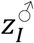 after insecticide exposure and selection, with “*E*” indicating these mosquitoes have survived insecticide exposure. 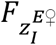 is calculated as the frequency of 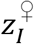 of females at birth 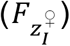 multiplied by the probability of insecticide encounter (*x*) and the probability of surviving the insecticide exposure 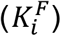. Survival is dependent on 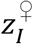 and the current insecticide efficacy of insecticide 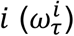 as calculated from Equation 2b(i).

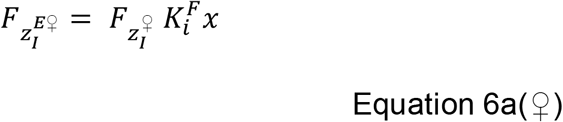

For males, where 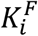 which is instead dependent on 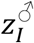, and the probability of insecticide encounter is *xm*:

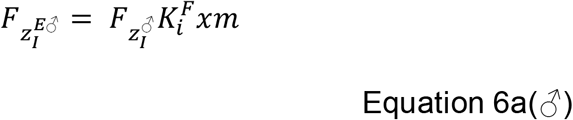

To extend to mixtures where mosquitoes simultaneously encounter both insecticides, and therefore the mosquitoes must also survive the encounter with insecticide *j*:

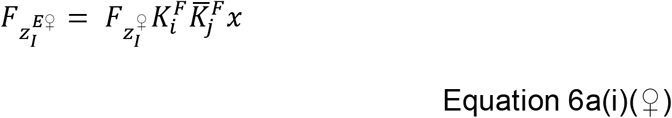

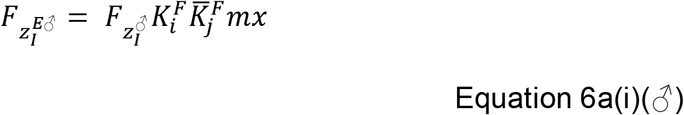

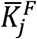 is the mean population survival to insecticide *j* and is calculated from Equation 2b(i) and is dependent on the mean resistance to insecticide 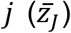 and the current efficacy of insecticide 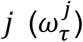. The model does not allow for individual level genetic correlation but includes population level cross resistance (see Equations 8b, 8c and 8d later).

The total number of mosquitoes surviving insecticide exposure (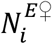 and 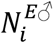), is therefore the sum of the frequencies of the exposed survivors (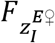 and 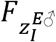).

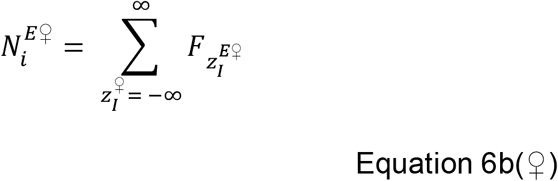

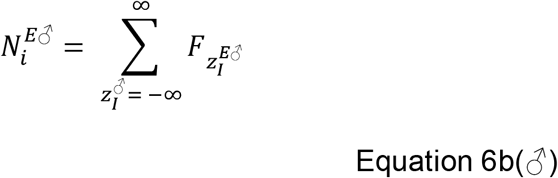

The mean PRS of the exposed survivors is then calculated:

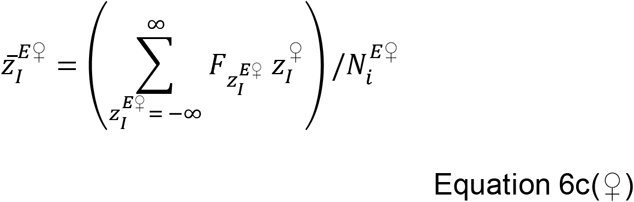

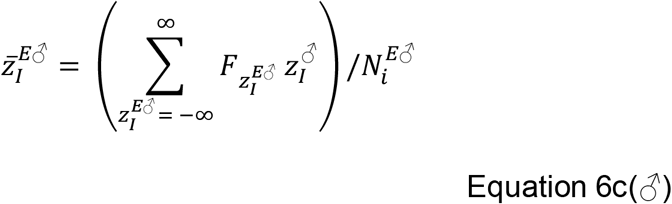

For mixtures the terms 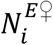 and 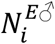 are replaced by 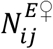 and 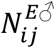 Equations 6b and 6c). The values of 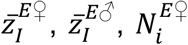 and 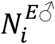 are returned to Equations 4b. If a mixture was deployed 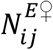 and 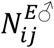 replaces 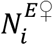 and 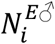 in Equations 4b. The solving of Equations 4b gives 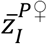 and 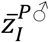, which is required to calculate the insecticide selection differential (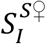 and 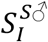) which can be combined with any fitness costs to give the overall selection differentials which are used in the sex-specific Breeder’s Equations (Equation 3c) to calculate the overall response which can be passed through to Equation 8 to give the mean for the next generation.

### Methods Section 2.5: Calculating the Selection Differential: Fitness Costs

Fitness costs can be associated with IR (Freeman et al., 2021). There are perhaps two intuitive ways to calculate the selection differential associated with fitness costs 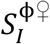 and 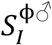. Option 1 assumes the fitness cost selection differential remains a fixed constant over the course of selection. Option 2 instead assumes the fitness selection differential varies with *σ*_*I*_. Option 1 is used when *σ*_*I*_ is a fixed value, and option 2 is used when *σ*_*I*_ changes with 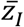. This is detailed in Equation 7 in Supplement 6.

### Methods Section 2.6: Between Generation Change in Resistance

The simulations consist of two operational sites, the intervention and the refugia sites. The refugia is the part of the landscape where insecticides are never deployed. For convenience we assume a single intervention site and a single refugia which can be viewed as amalgamation all intervention sites together and all refugia together with a global migration between the sites (see section 2.7 below). Insecticide deployments and the decisions regarding deployment are made in the intervention site.

For insecticide deployments 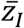 is tracked in the intervention site 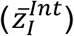 and refugia 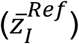 for each insecticide in each generation. When insecticide *i* is deployed in monotherapy the response is a result of both insecticide selection and fitness costs.

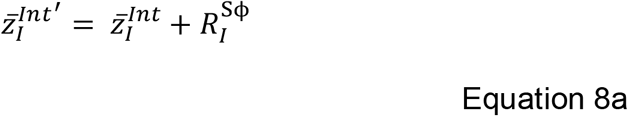

If insecticide *i* is not deployed and insecticide *γ* is deployed, there may be indirect selection on trait *I* from insecticide *γ* from the genetic correlation between trait Γ and trait *I*, termed *α*_ΓI_ (see Hobbs et al., 2023 for details). This effect is referred to as both “cross selection” or “cross resistance” and the terms are often used interchangeably, and we use the term cross resistance.

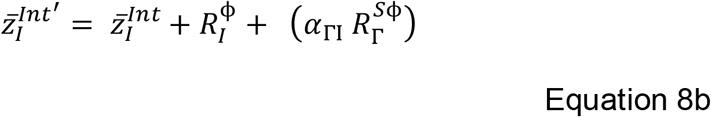

Where *γ* ∈ {*j, k, l* … } is the set of insecticides that may be deployed other than insecticide *i*. Where Γ ∈ {*J, K, L* … } is the set of traits that correspond to the PRS to the corresponding insecticide except for trait *I*.

For mixture insecticide deployments when insecticide *i* is deployed in a mixture formulation consisting of insecticides *i* and *j* and there may be cross resistance between both insecticides:

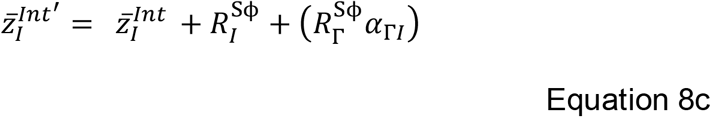

If insecticide *i* is not deployed, but a mixture of *j* and *k* is deployed, there may be indirect selection on trait *I* from both insecticide *j* and insecticide *k*.

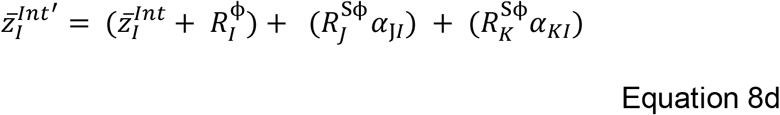

In the refugia, where insecticides are not deployed, the response 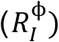 is only from fitness costs regardless of whether insecticides are deployed as monotherapies or mixtures:

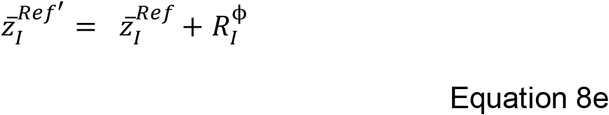

### Methods Section 2.7 Mosquito Dispersal

After insecticide selection and mating, mosquitoes can migrate between the intervention site and refugia (Equations 9a and 9b). Dispersal results in gene flow that will alter the dynamics of how resistance evolves so must be an option in the modelling (see Hastings et al., 2022 for details). Dispersal is described as follows:

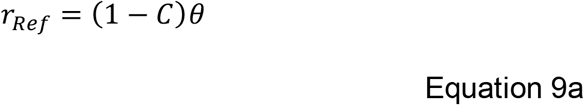

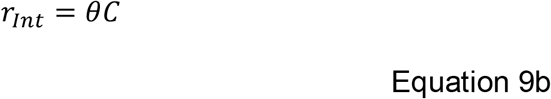

Where *C* is the coverage of the insecticide and corresponds to the proportion of the mosquito population in the intervention site. *θ* is the proportion of female mosquitoes dispersing. *r*_*Int*_ is the number of female mosquitoes migrating from the intervention site to the refugia. *r*_*Ref*_ is the number of female mosquitoes migrating from the refugia to the intervention site.

The mean resistance of the eggs being laid by females in the intervention site is therefore:

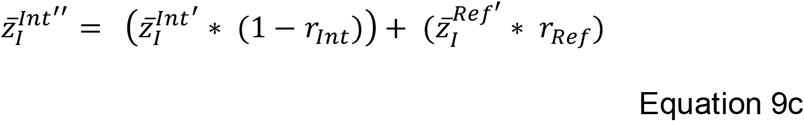

The mean resistance of the eggs being laid by females in the refugia is therefore:

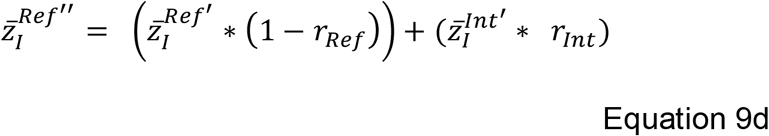

Note the use of primes: single primes for IR levels before dispersal, and double primes after dispersal: it is the double prime 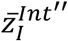 and 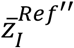 which become the mean PRS for the next generation in each site (intervention and refugia).

### Methods Section 3: Biology of Insecticide Selection Over Multiple Gonotrophic Cycles

A proposed benefit of micro-mosaic and combination deployments is a “temporal mixture” where a mosquito surviving one insecticide is killed by a different insecticide subsequent gonotrophic cycle. Evaluation of micro-mosaics and combinations therefore requires incorporating insecticide selection over multiple gonotrophic cycles. This is technically complex (see below) but is justified by allowing the models to accurately simulate deployment of micro-mosaics and combinations.

Extending “polytruncate” to multiple gonotrophic cycles appears problematic because it is unclear how to implement the truncation process after selection in the first cycle, when the frequency distribution is no longer a Normal distribution. We therefore extend only the “polysmooth” model to include multiple gonotrophic cycles in the absence of dispersal. We subsequently further extend to allow dispersal between the intervention site and refugia between each gonotrophic cycle.

A conceptualisation of the selection process over multiple-gonotrophic cycles is shown in **Error! Reference source not found**.6. It highlights the number of females laying eggs would decrease with each subsequent gonotrophic cycle due to both insecticide induced and “natural” mortality (Figure 6, Panel 1). After each round of insecticide selection in each gonotrophic cycle it is expected the mean level of resistance of the surviving females increases (Figure 6, Panel 2). This increase in the mean level of resistance increases the female insecticide selection differential in each gonotrophic cycle. However, as females mate only once in the first gonotrophic cycle, they carry the same male selection differential for all gonotrophic cycle (Figure 6, Panel 3). This then corresponds to an increase in the response such that in each subsequent gonotrophic cycle eggs will be laid from which more resistant individuals will hatch (Figure 6, Panel 4), however there will be fewer of these eggs laid as the number of females in later gonotrophic cycles is lower (Figure 6, Panel 1). There is evidence that as mosquitoes age they become more suscepible to insecicides (Jones et al., 2012). Adding in senescence would therefore mean the first gonotrophic cycle becomes ever more important, in which case this can be modelled just assuming a single gonotrophic cycle as done previously.

**Figure 6.**
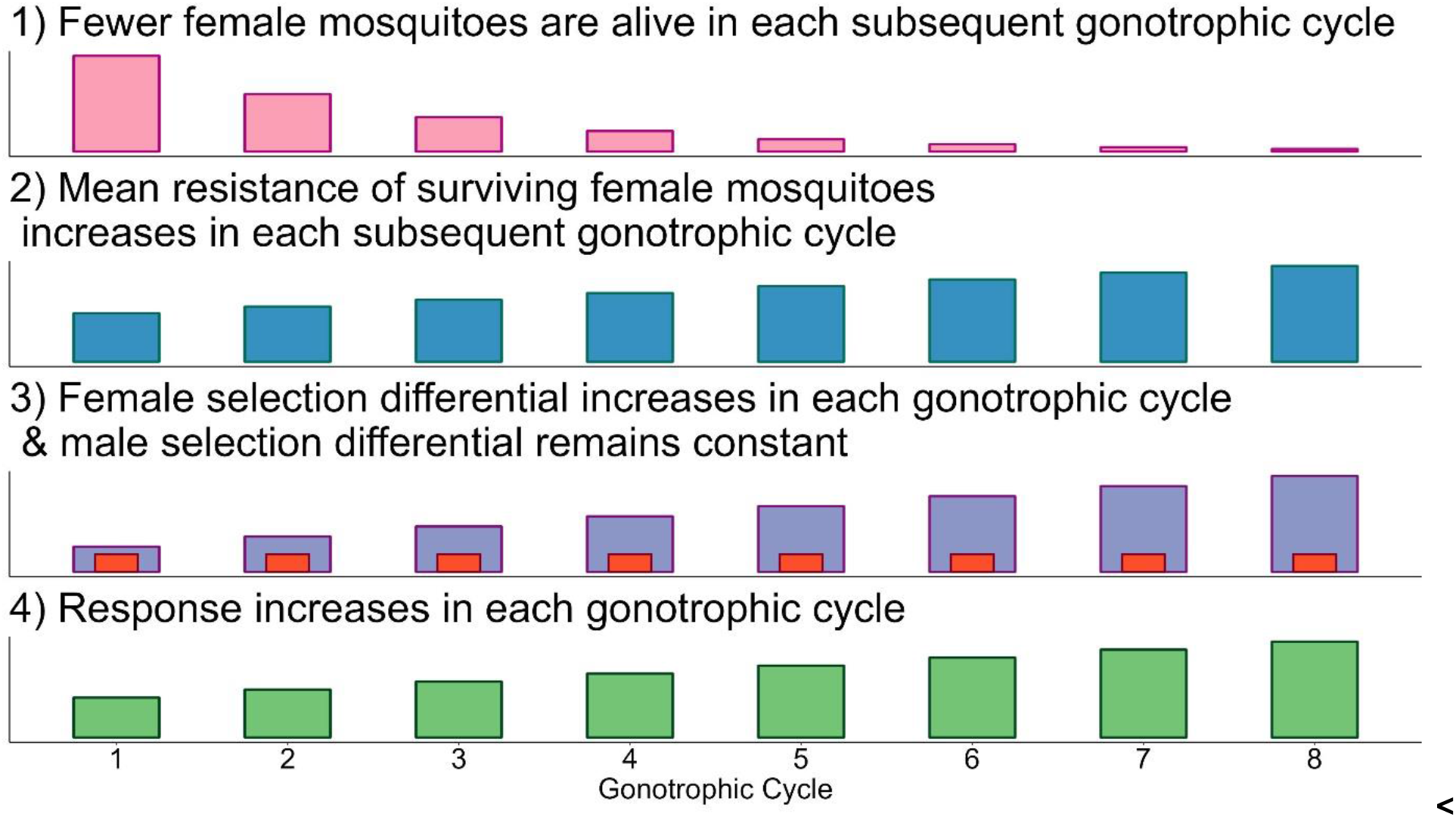
Conceptualisation of Insecticide Selection Over Multiple Gonotrophic Cycles. 1): The number of females in each subsequent gonotrophic cycle decreases due to insecticide selection (e.g.,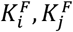) and natural mortality (*ρ*). This results in fewer eggs being laid in total in subsequent gonotrophic cycles. **2):** As a result of insecticide selection, surviving females have a higher mean PRS than the previous gonotrophic cycle as only resistant individuals would be able to survive multiple insecticide encounters. **3):** An increasing mean PRS of females gives an increased female insecticide selection differential (blue bars). The male selection differential (red bars) remains constant as female mosquitoes mate only once. **4):** As the female selection differential increases, the response increases. Eggs laid by females in subsequent gonotrophic cycles give more resistant individuals. The updated mean for the next generation is the mean of the eggs laid over all the gonotrophic cycles.

### Methods Section 3.1. Updating Coverages, Encounter Probabilities and Natural Survival

Combinations and micro-mosaics change how mosquitoes encounter insecticides. Combinations can have houses treated with two (or more) insecticides. For example, a house may receive both an LLIN (containing insecticide *i*) and IRS (containing insecticide *j*). Or the household may receive a LLIN with different insecticides on different panels (for example insecticide *i* on the top and insecticide *j* on the sides).

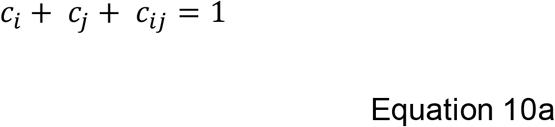

Where *c*_*i*_ is the proportion of treated houses with only insecticide *i, c*_*j*_ is the proportion of treated houses with only insecticide *j, c*_*ij*_ is the proportion of treated houses with both insecticide *i* and *j*. If *c*_*ij*_ is set to 0, the strategy becomes is a micro-mosaic:

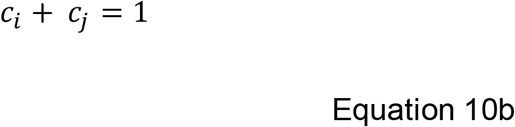

If *c*_*i*_ or *c*_*j*_ is further set to 1 then it is de-facto monotherapy.

Mosquitoes entering a house with two insecticides *i* and *j* which are spatially separated, may encounter *i* only (Λ_*i*|*ij*_), *j* only (Λ_*j*|*ij*_) or both *i* and *j* (Λ_*ij*|*ij*_). These parameters will be the result of mosquito behaviour (likely sex-specific), insecticidal properties and the distance between insecticides within the household. Mosquitoes encountering both insecticides (Λ_*ij*|*ij*_) are treated as though encountering a mixture.

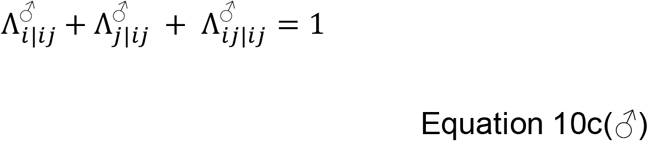

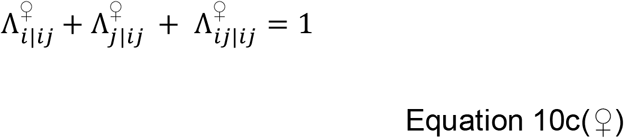

Note, if both 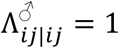 and 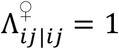, then the insecticides become a de-facto mixture. A compartmental description of the coverage-encounter process is given in Figure 7.

**Figure 7.**
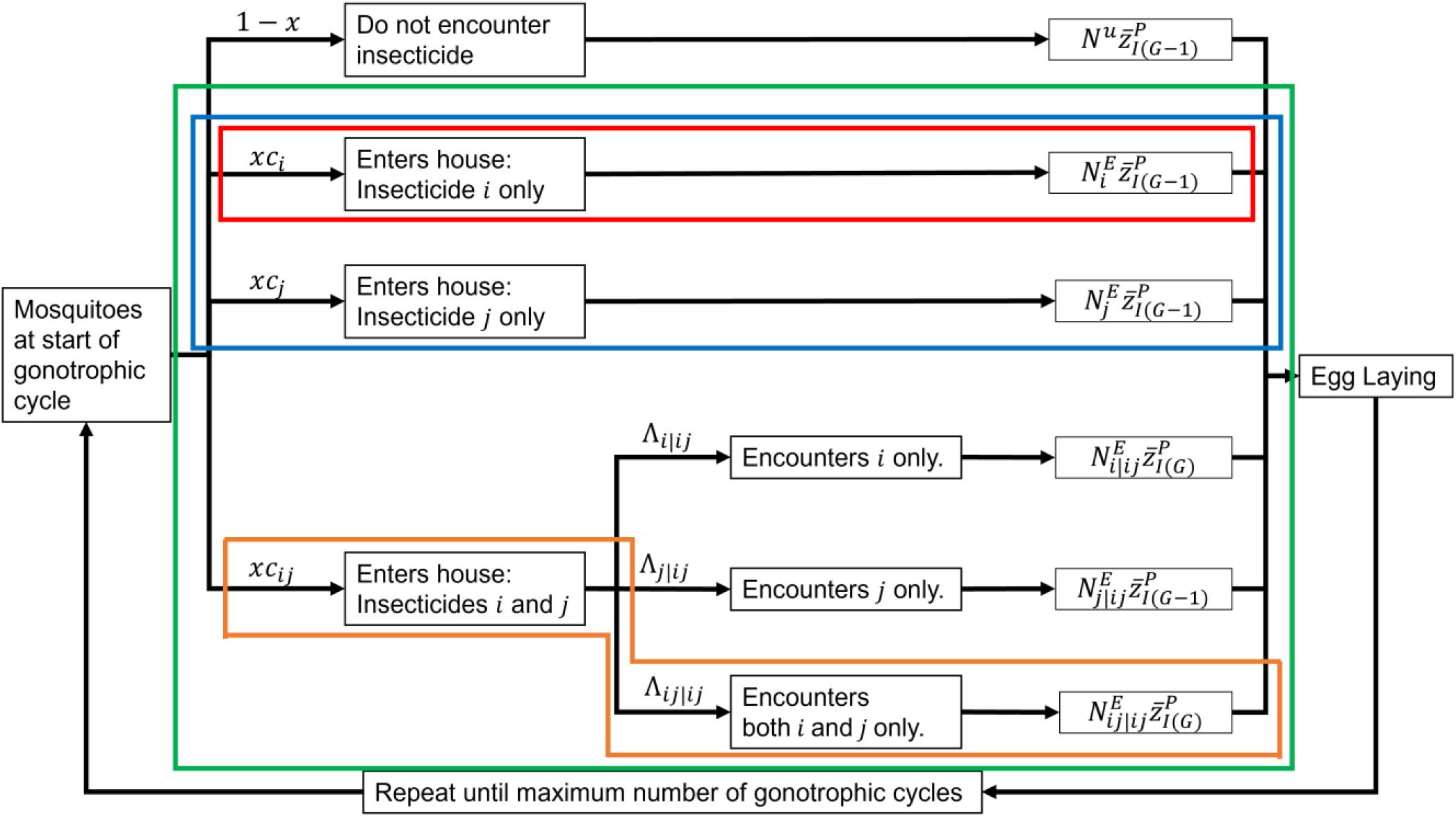
Compartmental diagram of the selection process over multiple gonotrophic cycles for IRM strategies. The unifying framework of the multiple gonotrophic cycle model allows for multiple different strategies to be evaluated: Red indicates only a single insecticide is deployed, orange is a mixture, blue is a micro-mosaic, and green is a combination of LLIN and IRS. These different strategies apply differing levels of selection producing different mean PRS values post insecticide selection and the total number of females surviving each gonotrophic cycle.

The multiple gonotrophic cycle model needs to account for natural survival between gonotrophic cycles (*ρ*). We assume all females taking a bloodmeal lay eggs in that gonotrophic cycle (as assumed in single gonotrophic cycle models) and each individual female lays the same number of eggs in each gonotrophic cycle (although it would be possible to relax this assumption by adding a decay term i.e., Equation 11c and 11d). The “natural” survival between gonotrophic cycles is:

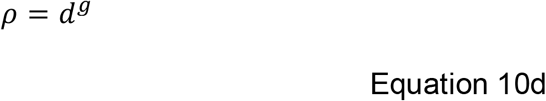

*ρ* is the proportion of female mosquitoes at the end of the previous gonotrophic cycle starting the next gonotrophic cycle and is dependent on the “natural” daily survival probability (*d*) (i.e., mortality not caused by insecticides) and gonotrophic cycle length in days (*g*). “Natural” daily survival is assumed to be constant for all PRS values and for each gonotrophic cycle.

### Methods Section 3.2. Updating equations for multiple gonotrophic cycles

The female insecticide selection differential for a gonotrophic cycle, (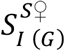, where *G*=1,2,3 to *G*_*max*_) is the change in the mean PRS between parents in the current gonotrophic cycle 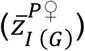 and at hatching 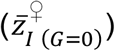:

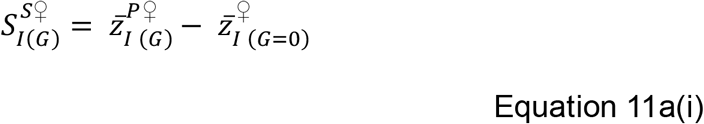

The overall selection differential 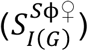 is from insecticide selection (subscript “*S*”) and fitness costs (subscript “ϕ”). We may expect most of the fitness costs to occur during larval development (e.g. Osoro et al., 2021) and/or mating (e.g. Platt et al., 2015), therefore the fitness cost selection differential remains constant between gonotrophic cycles:

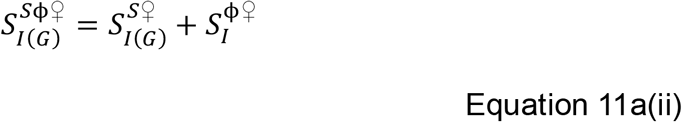

There is evidence that as mosquitoes age they not only become less resistant (Jones et al., 2012), but also produce fewer eggs for example. The fitness costs selection differential accounts for all aspects of life-history, for example larval growth rates, pupation rates and mating success, most of which will occur prior to or shortly after insecticide selection. However, as the model is not explicit in how the fitness costs are implemented (doing so would require an additional population dynamics structure).

The response 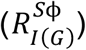 is calculated by updating the sex-specific Breeder’s equation (Equation 3c) for the current gonotrophic cycle:

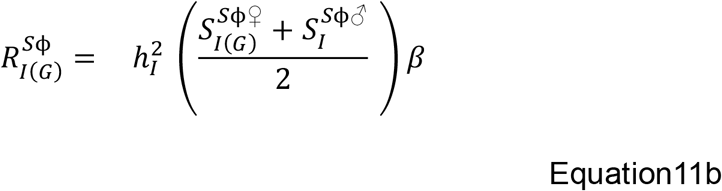

Surviving female mosquitoes progress through gonotrophic cycles until *G*_*max*_ is reached. *G*_*max*_ is a user-defined maximum number of gonotrophic cycles (we use a *G*_*max*_= 5 unless otherwise stated). When *G*_*max*_ is reached the total number of oviposition events (*N*_*o*_) is calculated:

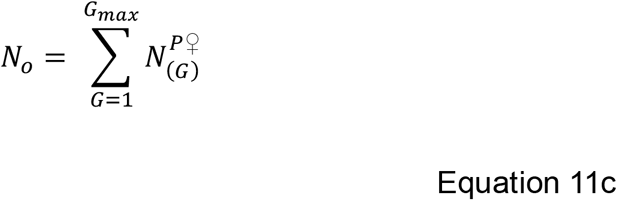

*N*_*o*_ is used below to weight the responses for each gonotrophic cycle 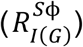 to calculate the overall response 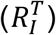. Cross resistance can be incorporated using correlated responses (see Equations 8c and 8d) such that each correlated response each gonotrophic cycle is also weighted by the number of oviposition events:

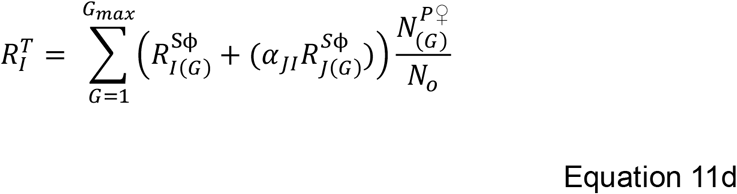

The mean PRS of the next generation is then calculated as the:

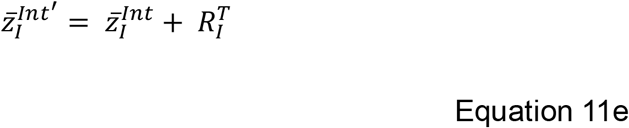

In the absence of refugia and dispersal, 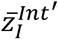 becomes the mean PRS of the next generation.

### Methods Section 3.3: Applying the dynamic model (“polysmooth”) for insecticide deployments across multiple gonotrophic cycles

We must first account for mosquitoes being able to contact complex insecticide deployments which will impact the level of selection on mosquitoes and therefore impact the insecticide selection differential.

This involves updating Equation 4b(*♂*), to allow for male mosquitoes to encounter just insecticide *i*, just insecticide *j* or both insecticide *i* and *j* in their single round of selection (Supplement 7, Equations 12a(*♂*) to Equation 12h(*♂*)). And also involves updating Equation 4b(*♀*), to allow for female mosquitoes to encounter just insecticide *i*, just insecticide *j* or both insecticide *i* and *j* in each gonotrophic cycle.

The mean PRS of the female mosquitoes laying eggs as the parental population for each gonotrophic cycle 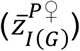 is calculated from all insecticide encounters, where the superscript “*P*” indicates the parental population. The superscript “*E*” indicates the mosquitoes survived their encounter with the insecticide(s) and the superscript “*u*” indicates those females did not encounter the insecticide in the current gonotrophic cycle. The mean PRS for females in a gonotrophic cycle 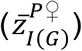 is therefore:

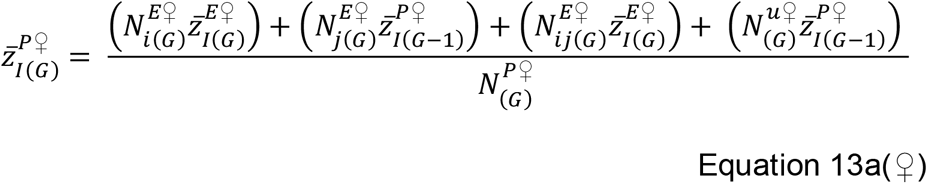

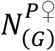 is the total number of female mosquitoes in gonotrophic cycle *G* laying eggs and is the sum of all female mosquitoes laying eggs in the current gonotrophic cycle:

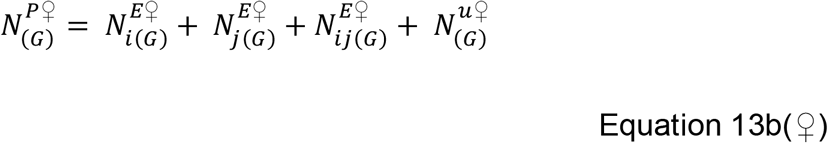

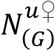 is the number of females not encountering any insecticides in gonotrophic cycle *G*. For the first gonotrophic cycle 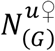 is calculated as initial number of females (*N*^*T♀*^, calculated from Equation 5f(*♀*)) multiplied by the probability of not encountering insecticides:

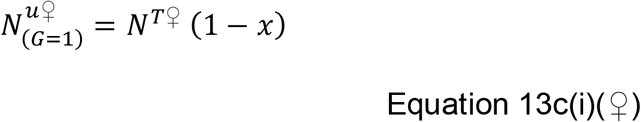

For all subsequent gonotrophic cycles, 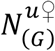 is calculated:

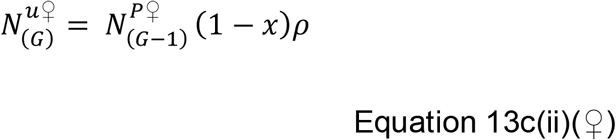

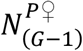 is the number of female mosquitoes at the end of the previous gonotrophic cycle and *ρ* is the between gonotrophic cycle survival probability (calculated from Equation 10d). These individuals 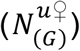 will have the same mean PRS as the individuals at the end of the previous gonotrophic cycle being 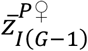.

The number surviving the encounter with insecticide *i* only is, for the first gonotrophic cycle:

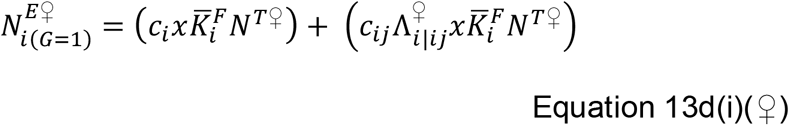

And for subsequent gonotrophic cycles:

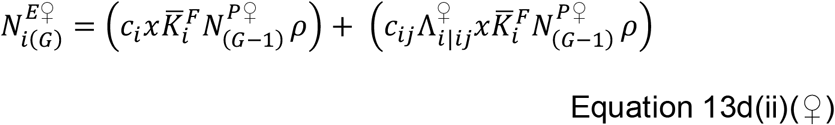

The number surviving the encounter with insecticide *j* only is, for the first gonotrophic cycle:

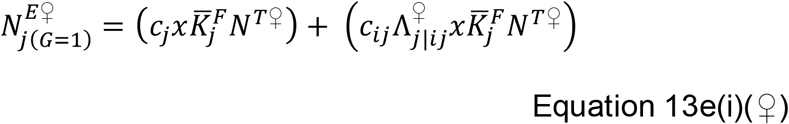

And for subsequent gonotrophic cycles:

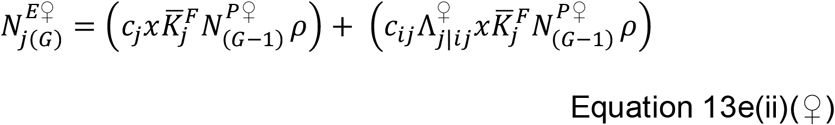

The number surviving the encounter with insecticide *i* and *j* is, for the first gonotrophic cycle:

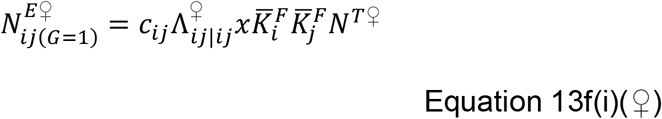

And for subsequent gonotrophic cycles:

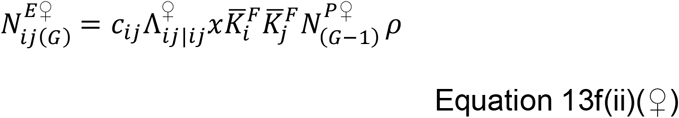

The frequency of the binned values of 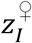 after insecticide selection for the first gonotrophic cycle is:

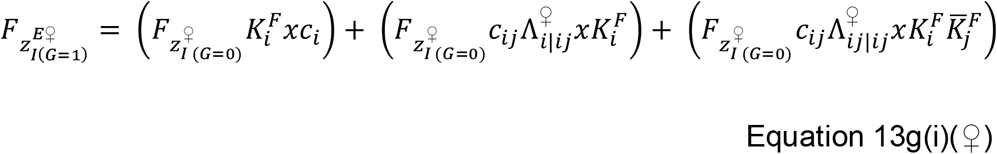

For all subsequent gonotrophic cycles:

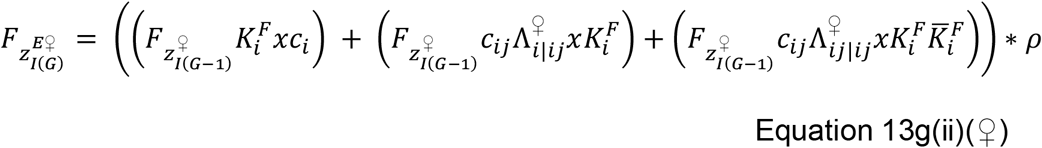

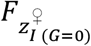 is the initial frequency of 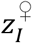 (see Equation 4g(*♀*)) and 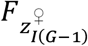 is the frequency at the end of the previous gonotrophic cycle. 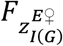 is transferred to equation 13i(*♀*). The frequency at the end of the first gonotrophic cycle is:

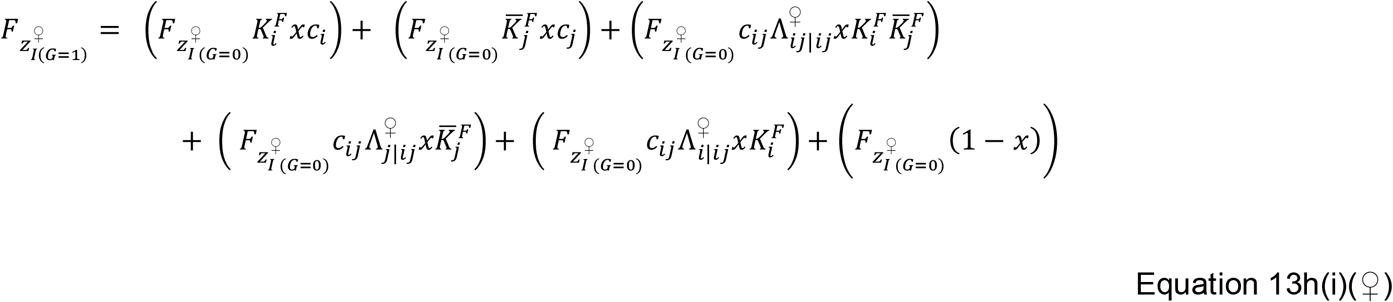

And for all subsequent gonotrophic cycles:

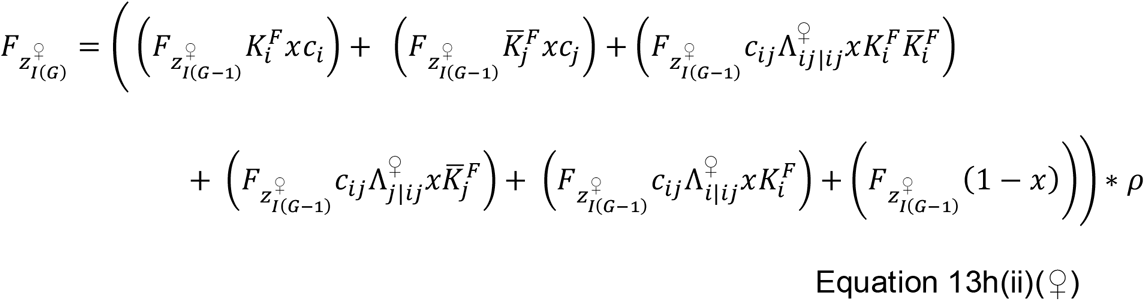

The mean PRS value of the exposed survivors 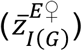 for each gonotrophic is calculated:

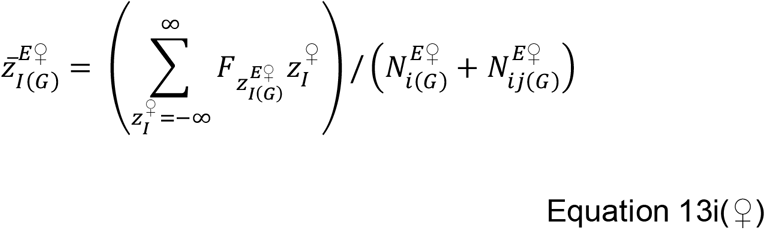

The value of 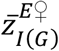 is then returned to Equation 13a(*♀*) to calculate 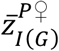, which is used to calculate the selection differentials (Equation 11a(i), Equation 11a(ii)) and response (Equation 11b).

Details of implementing the multiple gonotrophic cycle model allowing for dispersal between the intervention site and refugia in each gonotrophic cycle is given in Supplement 8.

## Methods Section 4: Calibrating and Running Simulations to Investigate IRM Strategies

### Methods Section 4.1. Parameter Estimation/Model Calibration

Details of parameter estimation and model calibration are found in the supplementary information. Supplement 5 details the estimation of the standard deviation (*σ*_*I*_) the estimation of how the standard deviation changes with regards to the mean PRS. Supplement 2 details the calibration of the models such that a “novel” insecticide would be expected to have an average of 10 years continuous deployment until 10% bioassay is reached, as was used in the “polyres” model (Hobbs et al., 2023) using a fixed standard deviation. Supplement 6 details the estimation of fitness cost selection differentials.

### Methods Section 4.2. Simulations: Comparing Polytruncate versus Polysmooth

We compare “polytruncate” and “polysmooth” against the IRM modelling literature and against each other. To allow for direct comparisons between strategies, simulations consisted of 2 insecticides (with equivalent properties) deployed as monotherapy sequences, monotherapy rotations and full-dose mixtures. The starting resistance was 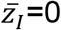. Simulations had a 10-generation deployment interval. Insecticide decay was not included, therefore allowing comparison as the previous “polyres” model did not include insecticide decay (Hobbs et al., 2023). Simulations used 10% and 8% as bioassay survival withdrawal and return thresholds respectively (Hobbs et al., 2023).

The parameter space of coverage, dispersal, fitness costs, female exposure, male exposure, heritability was sampled using a Latin hypercube (Carnell, 2020) to get 5000 parameter sets. These were replicated across cross-resistance values (−0.3 to 0.3 at 0.1 intervals) for a total of 35,000 parameter sets. The same parameter sets were used for each strategy and model (“polytruncate” and “polysmooth”) to allow for direct comparisons. Simulations were terminated when no insecticides were available for deployment (because resistance had evolved to them all), or the 500-generation (∼50 years) cap was reached.

The primary outcome was simulation duration (in generations) which were compared between sequences-rotations, sequences-mixtures, and rotations-mixtures. Strategies could therefore “win”, “lose” or “draw” in each comparison.

Agreement between “polytruncate” and “polysmooth” was assessed:

1. Full Agreement: both models indicate the same strategy performed best.
2. Agreement: both strategies draw in both models, this is because draws may not be equivalent if the simulations were allowed to run for longer durations (i.e., >500 generations).
3. Partial Disagreement: Both strategies drew in one model, but one strategy performed better than the other in the other model.
4. Full Disagreement: Models diverge as to which strategy performed best.

### Methods Section 4.3 Importance of Multiple Gonotrophic Cycles for Micro-Mosaics

The importance of including the multiple gonotrophic cycles for evaluating more complex IRM strategies was evaluated by comparing two insecticides (*i* and *j*) deployed as a rotation versus being deployed as a micro-mosaic (where the coverage of insecticide *i* (*c*_*i*_) is 0.5 and the coverage of insecticide *j* (*c*_*j*_) is 0.5). Each strategy used the same 5000 randomly sampled parameter values, allowing for direct comparison between the strategies. Parameters were sampled from uniform distributions using Latin hyperspace sampling (Carnell, 2020): Heritability 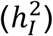 [0.05 to 0.3]; Female insecticide exposure (*x*) [0.4 to 0.9]; Male insecticide exposure (*m*)[0 to 1]; Female Fitness Cost 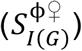 [0.04 to 0.58]; Male Fitness Cost 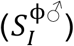 [0.04 to 0.58]; Intervention (*C*) [0.1 to 0.9]; Dispersal (*θ*)[0.1 to 0.9]. The male insecticide exposure is implemented as a proportion of the female insecticide exposure. The strategies were then additionally run using a single gonotrophic cycle, five gonotrophic cycles with no natural mortality, and finally with five gonotrophic cycles with a daily natural mortality (*ρ* = 0.8) and a gonotrophic cycle length of three days. Simulations were run for 500 generations (50 years), and the difference in bioassay survival to insecticide *i* was reported each year. No withdrawal threshold value was used so all simulations ran for the 50-year time horizon.

## 3. Results

### 3.1. Comparing the “polysmooth” and “polytruncate” Models

Comparing the “polysmooth” (Figure 8) and “polytruncate” (Figure 9) at the indicates these models give similar results to one another and is qualitatively similar to results obtained previously using the “polyres” model (Hobbs et al., 2023), i.e. full-dose mixtures are best, rotations perform better than sequences if there is negative cross resistance, and sequences perform better than rotations if there is positive cross resistance. These general conclusions are in-line with monogenic models (Hastings et al., 2022; Madgwick & Kanitz, 2022; South & Hastings, 2018).

**Figure 8.**
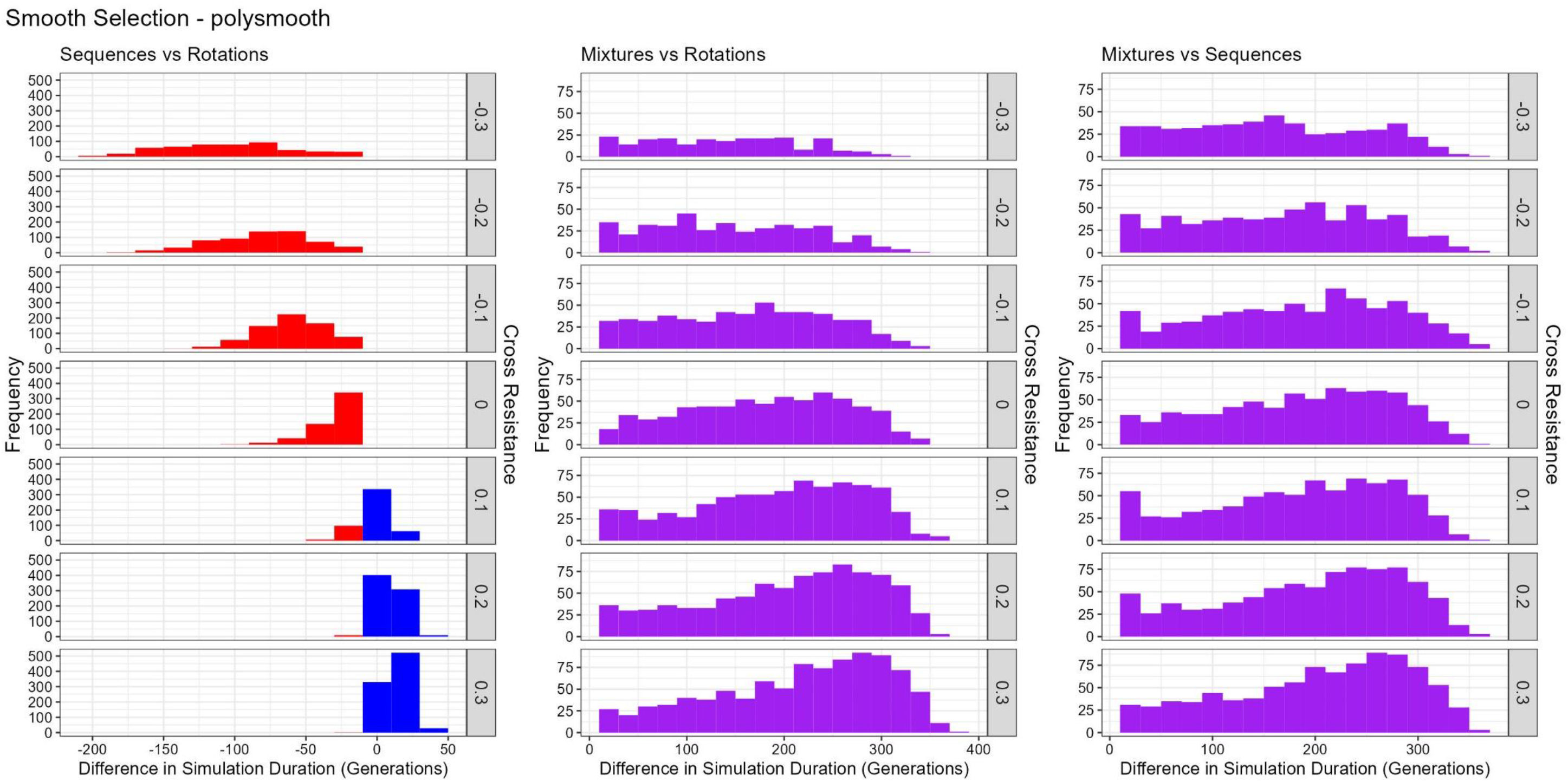
Comparison of Sequences, Rotations and Mixtures for Polysmooth. Left panel: Sequences versus Rotations; Centre Panel: Mixtures vs Rotations; Right Panel: Mixtures vs Sequences. Colours indicate which strategy had the longest duration in the direct comparison. Red = rotations lasted longest. Blue = Sequences lasted longest. Purple = Full-Dose Mixtures lasted longest. Draws were excluded. The rows indicate the amount of cross resistance between the two insecticides.

**Figure 9.**
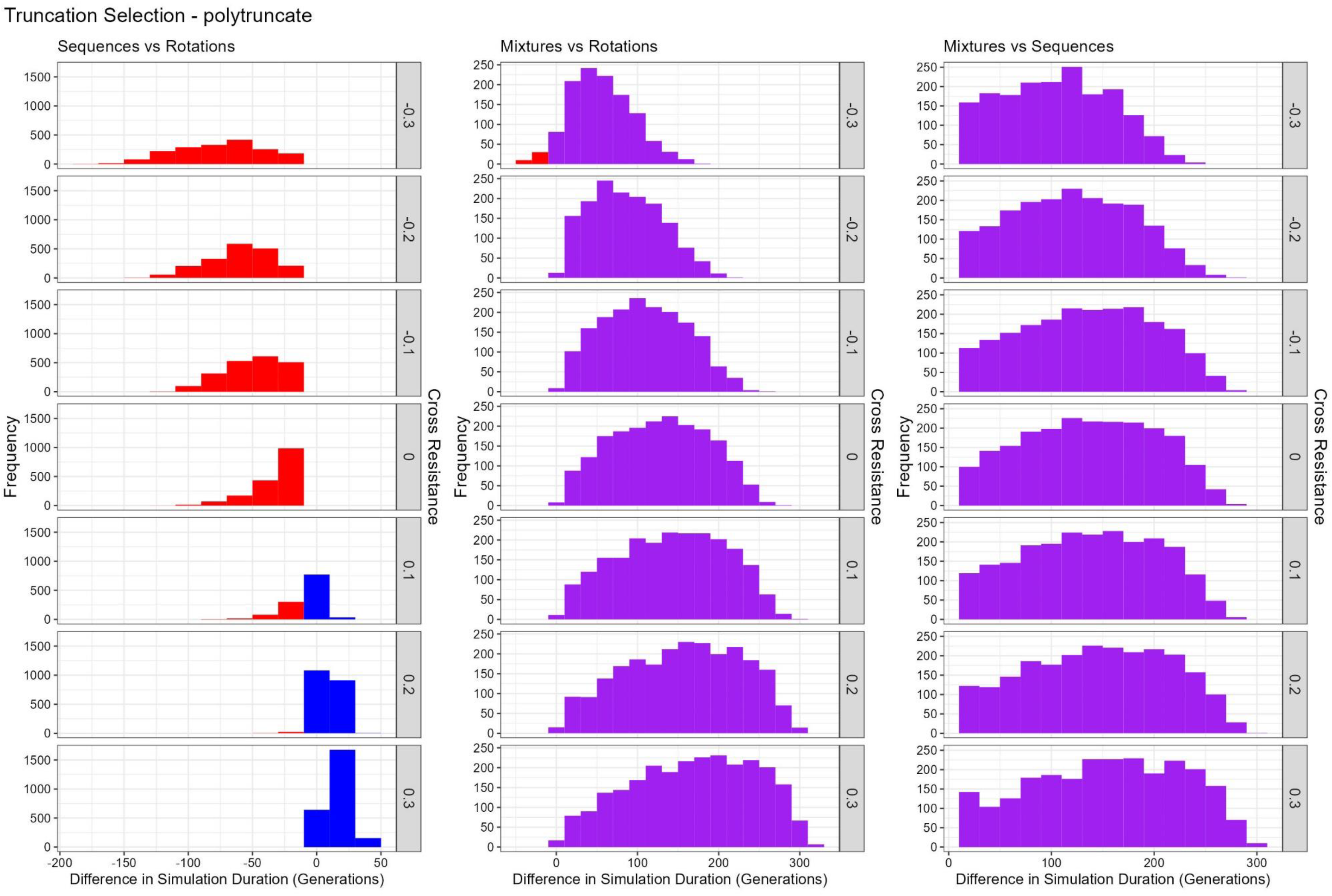
Comparison of Sequences, Rotations and Mixtures for Polytruncate. Left panel: Sequences versus Rotations; Centre Panel: Mixtures vs Rotations; Right Panel: Mixtures vs Sequences. Colours indicate which strategy had the longest duration in the direct comparison. Red = rotations lasted longest. Blue = Sequences lasted longest. Purple = Full-Dose Mixtures lasted longest. Draws were excluded. The rows indicate the amount of cross resistance between the two insecticides.

Figure 10 shows the percentage agreement between the models and shows “full disagreement” was rare, occurring only when comparing sequences and rotations. The choice between sequences and rotations has previously been shown to be highly unpredictable (Hastings et al., 2022), and therefore divergence between “polysmooth” and “polytruncate” is not unexpected for these two IRM strategies. Partial agreements were most common, due to the frequency of draws.

**Figure 10.**
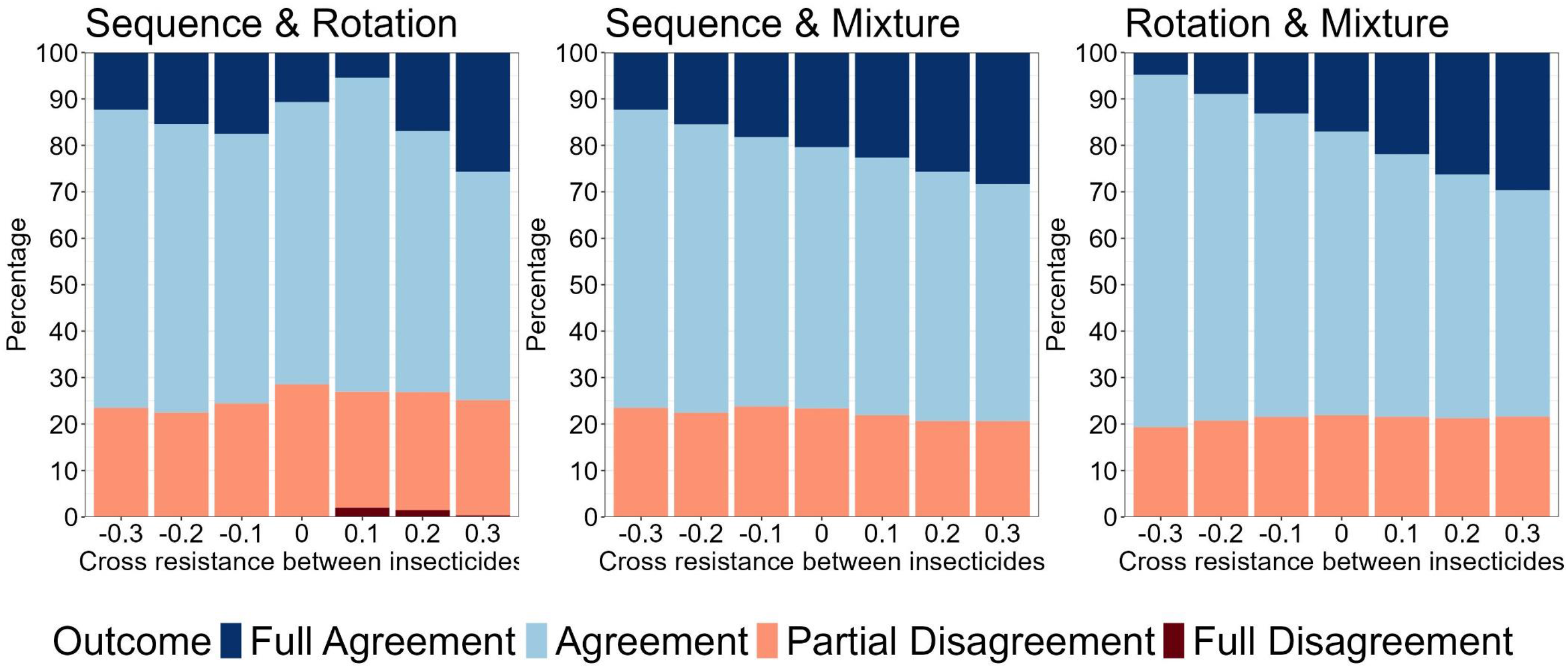
Comparing Outcomes between Polysmooth and Polytruncate. Looks at the difference in the strategy outcome between the “polysmooth” and “polytruncate” models for the same parameter inputs. Colours indicate how the strategies performed in both models. Dark blue = Both models indicate the same strategy performed best (“agreement”). Light blue = both strategies draw in both models (“agreement”). Dark red = Models diverge as to which strategy performed best (“full disagreement”). Light red = Both strategies drew in one model, but one strategy performed better than the other in the other model (“partial disagreement”).

### 3.2. Demonstrating the Importance of Multiple Gonotrophic Cycles and Natural Survival: Micro-Mosaics vs Rotations

While the quantitative effect may not be dramatic, the inclusion of multiple gonotrophic cycles and natural mortality can alter the qualitative results: micro-mosaics go from performing worse (single gonotrophic cycle, Figure 11 left panel) to performing equally (multiple gonotrophic cycles, Figure 11 centre panel), to micro-mosaics performing better (multiple gonotrophic cycles with natural mortality, Figure 11 right panel). This highlights why the inclusion of multiple gonotrophic cycles and natural mortality was required for the model. The reason for this is that the benefit of halving the selection over space becomes more pronounced as the first gonotrophic cycle becomes most important for insecticide selection combined with the benefit of the temporal mixture being more beneficial when including natural mortality also.

**Figure 11.**
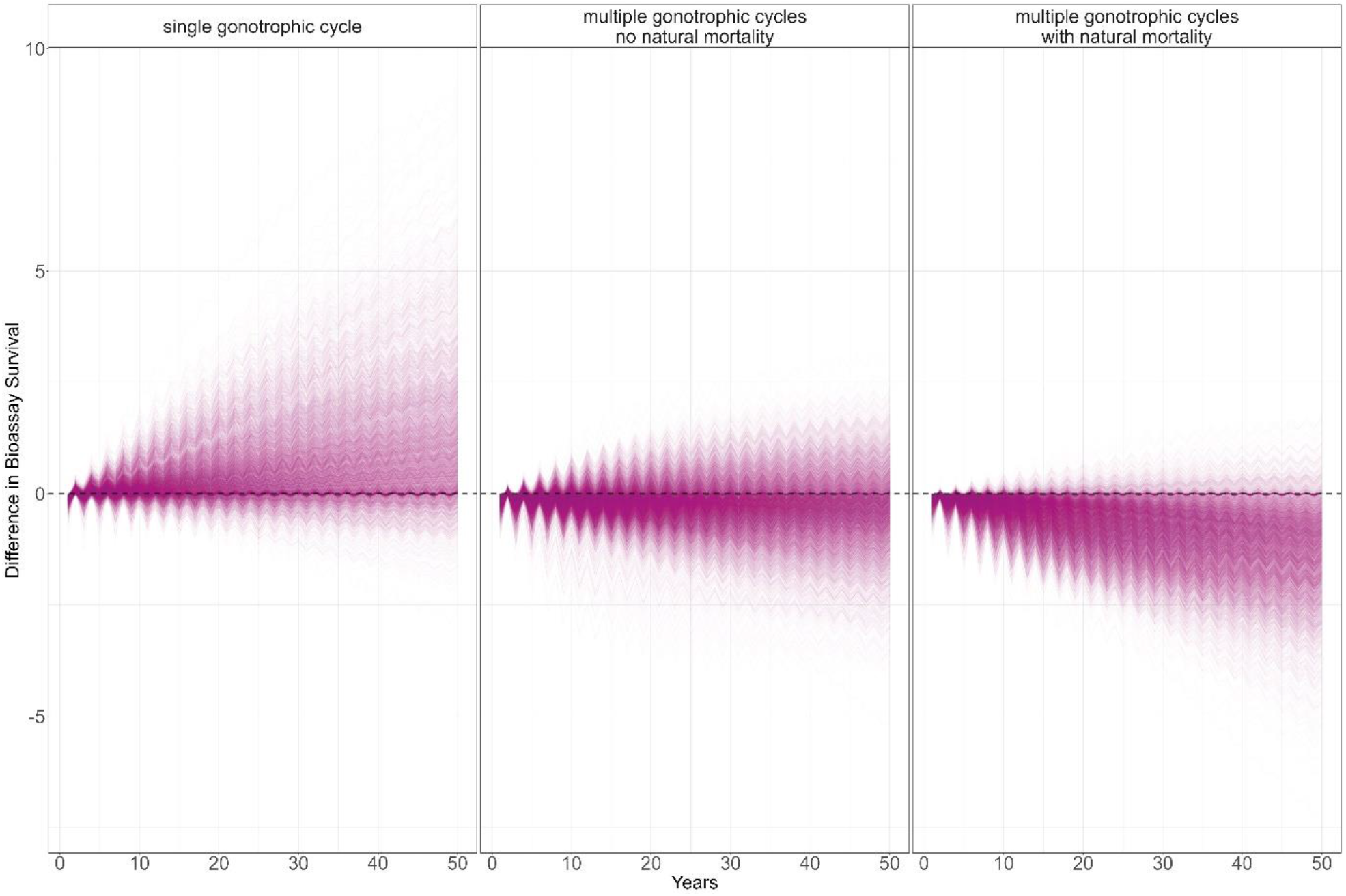
Plot of the impact of including multiple gonotrophic cycles and natural survival on the performance of deploying insecticides i and j in rotation versus micro-mosaics. Left Plot: The model is run only for a single gonotrophic cycle. **Centre Plot:** The model is run allow for multiple gonotrophic cycles (5 cycles) but natural survival is not included. **Right plot:** The model is run with multiple gonotrophic cycles and allowing for natural mortality (gonotrophic length = 3 days, natural daily survival = 0.8). Values above zero indicate the rotation strategy performed best, and values below zero indicates the micro-mosaic strategy performed best.

## 4. Discussion

We have presented a methodology for dynamically modelling the evolution of polygenic IR. This model provides a more sophisticated method for calculating the evolutionary trajectory of IR, allowing for the ability to evaluate the impacts of a larger number of parameters compared to a previous quantitative model assuming polygenic resistance (Hobbs et al., 2023).

### 4.1. Benefits of the Dynamic Model

The benefit of the developed dynamic polygenic model comes from an increase in the biological and operational realism, notably the simultaneous inclusion of three important factors: insecticide dosing, insecticide decay and cross resistance. These three factors are considered critical in the evaluation of IRM strategies despite being frequently absent from models (Rex Consortium, 2010; South et al., 2020).

An additional advantage of these dynamic models is the comparative ease with which an insecticide armoury of more than 2 insecticides can be investigated. Monogenic models are generally limited to two insecticides as tracking more than 2 loci involves tracking large numbers of genotypes and linkage dis-equilibriums, rapidly become computationally unfeasible. Allowing larger insecticide armouries to be considered allows for the evaluation of more complex IRM strategies, for example the combination strategy with rotations of IRS insecticides (see Supplement 1 for details and examples).

We also demonstrated the importance of extending the model to incorporate multiple gonotrophic cycles, and how this capability is required for micro-mosaics and combinations (Figure 11). The performance of micro-mosaics against rotations and mixtures (both full and reduced dose) is still to be evaluated across a wide biological and operational parameter space.

### 4.2. Consistency of Dynamic Models with Previous Model of IR

Despite these differences in complexity, the dynamic models generated results consistent with a previous quantitative genetics model (Hobbs et al., 2023) and monogenic models (e.g., Hastings et al., 2022; Madgwick & Kanitz, 2022; South & Hastings, 2018). Most notably full-dose mixtures being more effective than monotherapies as sequences or rotations, and little difference between the sequence and rotation strategies (Figures 9 and 10) (Hobbs et al., 2023). Future work will investigate issues surrounding lower dose mixtures especially with regards to cross resistance.

### 4.3. Future Applications for the Dynamic Models

The malERA consortium identified the need to evaluate mosaics and combinations of pyrethroid-LLINs with non-pyrethroid IRS (Rabinovich et al., 2017). The Rex Consortium noted half-dose mixtures required evaluation (Rex Consortium et al., 2013). These are areas which the presented dynamic models have the capability of evaluating, and we therefore highlight four priority areas to explore in future projects:

#### 4.3.1. Future Applications for the Dynamic Models: Insecticide Decay

Insecticide decay has recently been demonstrated as an important, yet often lacking, parameter in models evaluating IRM strategies when considering monogenic resistance (South et al., 2020). This issue requires further evaluation considering polygenic resistance.

#### 4.3.2. Future Applications for the Dynamic Models: Mixtures

One condition identified to maximise the benefit of mixtures is to have resistance to both insecticides to be at low levels (Rex Consortium, 2013). Current next-generation mixture LLINs include a pyrethroid as one of the insecticide partners. These next-generation mixture LLINs are showing better epidemiological efficacy than standard (pyrethroid-only) LLINs (Accrombessi et al., 2023; Mosha et al., 2022). Evaluating the long-term evolutionary consequences of mixtures where there are high levels of resistance to one of the partners (e.g., the pyrethroid) and potentially reduced insecticide dosages in next-generation mixture LLINs. Understanding the impact of reduced dosing and high levels of resistance is issue of immediate operational relevance.

#### 4.3.3. Future Applications for the Dynamic Models: Micro-mosaics

Micro-mosaics are likely to be a logistically complex strategy to implement. Micro-mosaics should be checked through computational simulations before extensive time, resources and finances are used in field trials. Simulations can also be used to understand any issues which may occur form micro-mosaics which may occur from this strategy occurring unintentionally due to differential distribution pathways (Worrall et al., 2020), and therefore potential mis-matched coverages of insecticides in the same setting.

#### 4.3.4. Future Applications for the Dynamic Models: Combinations

The evaluation of combinations for IRM remains lacking (Rabinovich et al., 2017), despite inclusion as a recommended IRM strategy in the Global Plan for Insecticide Resistance Management (WHO, 2012). This strategy was recently evaluated (S. Jones et al., 2023), however this methodology did not consider the benefit from being able to rotate out the insecticide used for the IRS independently of the pyrethroid-LLIN.

### 4.4. Conclusion

The presented dynamic models allow for simulation of complex IRM strategies and the implications of such strategies which would be impractical to evaluate in field settings. Simulations of more complex IRM strategies will help guide whether these complex strategies are worth pursuing or whether the proposed benefits of such strategies are large enough to justify the required complex procurement and deployment systems.

## Acknowledgements

The methodology developed in this manuscript has benefitted from extensive discussions with colleagues at the Liverpool School of Tropical Medicine. In particular, we would like to acknowledge the advice of David Weetman for help grounding the model in biological and operational relevance. We would also like to acknowledge the Vector Informatics and Genomics group at the Liverpool School of Tropical Medicine for their general critiques of the model methodology.

## References

Accrombessi, M., Cook, J., Dangbenon, E., Yovogan, B., Akpovi, H., Sovi, A., Adoha, C., Assongba, L., Sidick, A., Akinro, B., Ossè, R., Tokponnon, F., Aïkpon, R., Ogouyemi-Hounto, A., Padonou, G. G., Kleinschmidt, I., Messenger, L. A., Rowland, M., Ngufor, C., … Akogbeto, M. C. (2023). Efficacy of pyriproxyfen-pyrethroid long-lasting insecticidal nets (LLINs) and chlorfenapyr-pyrethroid LLINs compared with pyrethroid-only LLINs for malaria control in Benin: a cluster-randomised, superiority trial. The Lancet, 401(10375), 435–446. 10.1016/S0140-6736(22)02319-4

Alemayehu, E., Asale, A., Eba, K., Getahun, K., Tushune, K., Bryon, A., Morou, E., Vontas, J., Van Leeuwen, T., Duchateau, L., & Yewhalaw, D. (2017). Mapping insecticide resistance and characterization of resistance mechanisms in Anopheles arabiensis (Diptera: Culicidae) in Ethiopia. Parasites and Vectors, 10(1), 1–11. 10.1186/s13071-017-2342-y

Carnell, R. (2020). lhs: Latin Hypercube Samples (R package version 1.0.2). https://cran.r-project.org/package=lhs

Churcher, T. S., Lissenden, N., Griffin, J. T., Worrall, E., & Ranson, H. (2016). The impact of pyrethroid resistance on the efficacy and effectiveness of bednets for malaria control in Africa. ELife, 5(AUGUST), 1–26. 10.7554/eLife.16090

Curtis, C. F. (1985). Theoretical models of the use of insecticide mixtures for the management of resistance. Bulletin of Entomological Research, 75(2), 259–266. 10.1017/S0007485300014346

Freeman, J. C., Smith, L. B., Silva, J. J., Fan, Y., Sun, H., & Scott, J. G. (2021). Fitness studies of insecticide resistant strains: lessons learned and future directions. Pest Management Science, January. 10.1002/ps.6306

Hastings, I., Jones, S., & South, A. (2022). Evaluating rotations as a strategy to delay the evolution of insecticide resistance in vectors of human diseases. 1– 38. 10.1101/2022.10.07.511276

Hobbs, N., Weetman, D., & Hastings, I. (2023). Insecticide resistance management strategies for public health control of mosquitoes exhibiting polygenic resistance: a comparison of sequences, rotations, and mixtures. Evolutionary Applications, 16(4), 936–959. 10.1111/eva.13546

Jones, C. M., Sanou, A., Guelbeogo, W. M., Sagnon, N., Johnson, P. C. D., & Ranson, H. (2012). Aging partially restores the efficacy of malaria vector control in insecticide-resistant populations of Anopheles gambiae s.l. from Burkina Faso. Malaria Journal, 11, 1–11. 10.1186/1475-2875-11-24

Jones, S., South, A., Richardson, J., Rees, S., Sherrard-Smith, E., Small, G., Nimmo, D., & Hastings, I. (2023). How best to co-deploy insecticides to minimise selection for resistance. 1–29. 10.1101/2023.04.15.536881

Liu, N. (2015). Insecticide resistance in mosquitoes: Impact, mechanisms, and research directions. Annual Review of Entomology, 60, 537–559. 10.1146/annurev-ento-010814-020828

Madgwick, P. G., & Kanitz, R. (2022). Modelling new insecticide-treated bed nets for malaria-vector control: how to strategically manage resistance? Malaria Journal, 21(1), 1–20. 10.1186/s12936-022-04083-z

Martin, J., Lukole, E., Messenger, L. A., Aziz, T., Mallya, E., Bernard, E., Matowo, N. S., Mosha, J. F., Rowland, M., Mosha, F. W., Manjurano, A., & Protopopoff, N. (2024). Monitoring of Fabric Integrity and Attrition Rate of Dual-Active Ingredient Long-Lasting Insecticidal Nets in Tanzania: A Prospective Cohort Study Nested in a Cluster Randomized Controlled Trial. Insects, 15(2), 108. 10.3390/insects15020108

Matthews, J., Bethel, A., & Osei, G. (2020). An overview of malarial Anopheles mosquito survival estimates in relation to methodology. Parasites and Vectors, 13(1), 1–12. 10.1186/s13071-020-04092-4

Mechan, F., Katureebe, A., Tuhaise, V., Mugote, M., Oruni, A., Onyige, I., Bumali, K., Thornton, J., Maxwell, K., Kyohere, M., Kamya, M. R., Mutungi, P., Kigozi, S. P., Yeka, A., Opigo, J., Maiteki-Sebuguzi, C., Gonahasa, S., Hemingway, J., Dorsey, G., … Lynd, A. (2022). LLIN Evaluation in Uganda Project (LLINEUP): The fabric integrity, chemical content and bioefficacy of long-lasting insecticidal nets treated with and without piperonyl butoxide across two years of operational use in Uganda. Current Research in Parasitology & Vector-Borne Diseases, 2(April), 100092. 10.1016/j.crpvbd.2022.100092

Mosha, J. F., Kulkarni, M. A., Lukole, E., Matowo, N. S., Pitt, C., Messenger, L. A., Mallya, E., Jumanne, M., Aziz, T., Kaaya, R., Shirima, B. A., Isaya, G., Taljaard, M., Martin, J., Hashim, R., Thickstun, C., Manjurano, A., Kleinschmidt, I., Mosha, F. W., … Protopopoff, N. (2022). Effectiveness and cost-effectiveness against malaria of three types of dual-active-ingredient long-lasting insecticidal nets (LLINs) compared with pyrethroid-only LLINs in Tanzania: a four-arm, cluster-randomised trial. The Lancet, 399(10331), 1227–1241. 10.1016/S0140-6736(21)02499-5

Mutagahywa, J., Ijumba, J. N., Pratap, H. B., Molteni, F., Mugarula, F. E., Magesa, S. M., Ramsan, M. M., Kafuko, J. M., Nyanza, E. C., Mwaipape, O., Rutta, J. G., Mwalimu, C. D., Ndong, I., Reithinger, R., Thawer, N. G., & Ngondi, J. M. (2015). The impact of different sprayable surfaces on the effectiveness of indoor residual spraying using a micro encapsulated formulation of lambda-cyhalothrin against Anopheles gambiae s.s. Parasites and Vectors, 8(1), 1–7. 10.1186/s13071-015-0795-4

Osoro, J. K., Machani, M. G., Ochomo, E., Wanjala, C., Omukunda, E., Munga, S., Githeko, A. K., Yan, G., & Afrane, Y. A. (2021). Insecticide resistance exerts significant fitness costs in immature stages of Anopheles gambiae in western Kenya. Malaria Journal, 20(1), 1–7. 10.1186/s12936-021-03798-9

Platt, N., Kwiatkowska, R. M., Irving, H., Diabaté, A., Dabire, R., & Wondji, C. S. (2015). Target-site resistance mutations (kdr and RDL), but not metabolic resistance, negatively impact male mating competiveness in the malaria vector Anopheles gambiae. Heredity, 115(3), 243–252. 10.1038/hdy.2015.33

Rabinovich, R. N., Drakeley, C., Djimde, A. A., Hall, B. F., Hay, S. I., Hemingway, J., Kaslow, D. C., Noor, A., Okumu, F., Steketee, R., Tanner, M., Wells, T. N. C., Whittaker, M. A., Winzeler, E. A., Wirth, D. F., Whitfield, K., & Alonso, P. L. (2017). malERA: An updated research agenda for malaria elimination and eradication. PLoS Medicine, 14(11), 1–16. 10.1371/journal.pmed.1002456

Rex Consortium. (2007). Structure of the scientific community modelling the evolution of resistance. PLoS ONE, 2(12). 10.1371/journal.pone.0001275

Rex Consortium. (2010). The skill and style to model the evolution of resistance to pesticides and drugs. Evolutionary Applications, 3(4), 375–390. 10.1111/j.1752-4571.2010.00124.x

Rex Consortium. (2013). Heterogeneity of selection and the evolution of resistance. Trends in Ecology and Evolution, 28(2), 110–118. 10.1016/j.tree.2012.09.001

Rex Consortium, Bourguet, D., Delmotte, F., Franck, P., Guillemaud, T., Reboud, X., Vacher, C., Bordeaux, U., & Walker, A. S. (2013). Heterogeneity of selection and the evolution of resistance. Trends in Ecology and Evolution, 28(2), 110–118. 10.1016/j.tree.2012.09.001

South, A., & Hastings, I. M. (2018). Insecticide resistance evolution with mixtures and sequences: A model-based explanation. Malaria Journal, 17(1), 1–20. 10.1186/s12936-018-2203-y

South, A., Lees, R., Garrod, G., Carson, J., Malone, D., & Hastings, I. (2020). The role of windows of selection and windows of dominance in the evolution of insecticide resistance in human disease vectors. Evolutionary Applications, 13(4), 738–751. 10.1111/eva.12897

Thomsen, E. K., Strode, C., Hemmings, K., Hughes, A. J., Chanda, E., Musapa, M., Kamuliwo, M., Phiri, F. N., Muzia, L., Chanda, J., Kandyata, A., Chirwa, B., Poer, K., Hemingway, J., Wondji, C. S., Ranson, H., & Coleman, M. (2014). Underpinning sustainable vector control through informed insecticide resistance management. PLoS ONE, 9(6). 10.1371/journal.pone.0099822

Toé, K. H., Mechan, F., Tangena, J. A. A., Morris, M., Solino, J., Tchicaya, E. F. S., Traoré, A., Ismail, H., Maas, J., Lissenden, N., Pinder, M., Lindsay, S. W., Tiono, A. B., Ranson, H., & Sagnon, N. (2019). Assessing the impact of the addition of pyriproxyfen on the durability of permethrin-treated bed nets in Burkina Faso: A compound-randomized controlled trial. Malaria Journal, 18(1), 1–16. 10.1186/s12936-019-3018-1

Walsh, B., & Lynch, M. (2018). Evolution and Selection of Quantitative Traits. Oxford University Press.

Wat’Senga, F., Manzambi, E. Z., Lunkula, A., Mulumbu, R., Mampangulu, T., Lobo, N., Hendershot, A., Fornadel, C., Jacob, D., Niang, M., Ntoya, F., Muyembe, T., Likwela, J., Irish, S. R., & Oxborough, R. M. (2018). Nationwide insecticide resistance status and biting behaviour of malaria vector species in the Democratic Republic of Congo. Malaria Journal, 17(1), 1–13. 10.1186/s12936-018-2285-6

WHO. (2012). Global Plan for Insecticide Resistance Management.

WHO. (2018). Test procedures for insecticide resistance monitoring in malaria vector mosquitoes (Second edition) (Updated June 2018). In Who. http://www.who.int/malaria/publications/atoz/9789241511575/en/

World Health Organization. (2022). Manual for monitoring insecticide resistance in mosquito vectors and selecting appropriate interventions. World Health Organisation. https://www.who.int/publications/i/item/9789240051089

Worrall, E., Were, V., Matope, A., Gama, E., Olewe, J., Mwambi, D., Desai, M., Kariuki, S., Buff, A. M., & Niessen, L. W. (2020). Coverage outcomes (effects), costs, cost-effectiveness, and equity of two combinations of long-lasting insecticidal net (LLIN) distribution channels in Kenya: a two-arm study under operational conditions. BMC Public Health, 20(1), 1–16. 10.1186/s12889-020-09846-4

