## Supplement 1 for "Mathematical Methodology for Dynamic Models of Insecticide Selection Assuming a Polygenic Basis of Resistance"

### Supplement 1: Examples of Insecticide Resistance Management Strategies

Here we detail the IRM strategies and decision-making rules used in the models. We make a distinction between how insecticides are deployed (e.g., monotherapies, mixtures, micro-mosaics, and combinations) and how insecticides are switched (e.g., rotations, sequences). With more complex IRM deployments (e.g., mixtures, micro-mosaics, and combinations (LLIN + IRS)) there can be more complex deployment decisions. Table S1 details the definitions used to describe IRM strategies. Figures S1.1 to S1.4 provide visual examples. Note, the example simulations used parameter values which best illustrate and describe the functioning of the strategies and therefore should not be compared against one another. These examples are given to highlight the capability of the "polysmooth" and "polytruncate" models.

**Table S1: Describing Insecticide Resistance Management Strategies: Deployments, Switches and Thresholds.** Here we provide a full of what is meant by each IRM deployment and switching strategy

| **Insecticide Deployment Strategy Description** | **Insecticide Switching Strategy Description** |
|---|---|
| MONOTHERAPIES: The formulation contains only a single insecticide. Only one insecticide is deployed at any time. Example Simulations: Figure S1.1 | SEQUENCES: Insecticide $i$ is deployed continuously until it reaches a pre-defined withdrawal threshold. At this point insecticide $i$ is said to have "failed". Insecticide $i$ is then withdrawn from deployment (and being available for deployment) and is replaced by the next insecticide, insecticide $j$. Insecticide $j$ is then deployed continuously, until it too has failed. When insecticide $j$ reaches the pre-defined withdrawal threshold insecticide $j$ is then withdrawn. This process continues until all insecticides have been withdrawn. |
| | ROTATIONS: Insecticides are switched at each opportunity, $i \rightarrow j \rightarrow k \rightarrow i \rightarrow j \rightarrow k$. The rotation strategy fails when either no insecticides are available to deployed (they are all "failed" insecticides), or a rotation is not possible, such that insecticide $i$ is being redeployed immediately (is deployed in sequence). |
| | ADAPTIVE ROTATIONS: The condition in rotations whereby the strategy had failed when an insecticide would be immediately re-deployed. Instead, the strategy is allowed to be as a sequence when required, but defaults back to rotations whenever possible: $i \rightarrow j \rightarrow i \rightarrow j \rightarrow i \rightarrow i$. |
| MIXTURES: A mixture involves two different insecticides being deployed in the same formulation, such that any | ROTATE NOVEL PARTNER: In these scenarios, one mixture partner is common to all the mixture formulations, as is the case with current mixture LLINs, where a pyrethroid partner is common to all. Here, the novel insecticide is changed at each deployment opportunity, but the pyrethroid partner remains deployed throughout. Deployment decisions are made only on the novel insecticide partner, such that even if the pyrethroid partner exceeds the withdrawal threshold the mixture can still be deployed. For example, mixture |

| | |
|---|---|
| mosquito which contacting the mixture formulation encounters both insecticides simultaneously. Example Simulations: Figure S1.2 and S1.3 | A is a mixture of $i$ and $j$, and mixture B is a mixture of $i$ and $k$. The mixtures are then deployed: A→B→A→B. Example Simulation: Figure S1.2. |
| | SEQUENCE NOVEL PARTER: In these scenarios, one mixture partner is common to all the mixture formulations, as is the case with current mixture LLINs, where a pyrethroid partner is common to all. Here, the novel insecticide is deployed in sequence and deployment decisions are made on the novel insecticide partner, for example A→A→A→B→B. |
| | ROTATE MIXTURE FORMULATION: The mixture formulation is rotated at each opportunity. For example, mixture A of insecticides $i$ and $j$ is rotated with mixture B of insecticides $k$ and $m$ are rotated: A→B→A→B. This differs from the Rotate novel partner strategy in that if any insecticide in the mixture reaches the withdrawal threshold the insecticide is no longer available for deployment and therefore the mixture is also not available for deployment. |
| | SEQUENCE MIXTURE FORMULATION: The mixture formulation is deployed in sequence. For example, if two mixture formulations are available (mixture A [$i$ and $j$] and mixture B [$k$ and $m$] are deployed in sequence: A→A→A→B→B. This differs from the rotate novel partner strategy in that if any insecticide in the mixture reaches the withdrawal threshold the insecticide is no longer available for deployment and therefore the mixture is also not available for deployment. |
| MICRO-MOSAICS: Micro-mosaics involve deploying two insecticides in the same village, but such that each household receives only one insecticide. Example Simulation: Figure S1.4 | INDIVIDUAL SEQUENCE: For example, if insecticide $i$ and $j$ were both being deployed as a micro-mosaic, the deployment decisions are made individually on each insecticide. If insecticide $i$ reached the defined failure threshold, it is withdrawn and replaced with insecticide $k$; and this is not impacted by insecticide $j$. |
| | FULL ROTATION: Both insecticides deployed in the micro-mosaic are rotated out at each opportunity: $i$+$j$ →$k$+$l$ → $i$+$j$ →$k$+$l$. |
| | PARTIAL ROTATION: Only one of the insecticides in the micro-mosaic is rotated. For example, in areas where pyrethroid IRS is used (and is currently effective), this would be used every year (as cheaper) with the other insecticides rotated: $i$+$j$ →$i$+$k$ → $i$+$j$. |
| COMBINATIONS: Insecticides are deployed on both an LLIN and as an IRS, households can therefore receive just an LLIN, just an IRS or both the LLIN and IRS. Example Simulation: Figure S1.5 | ROTATE IRS: Standard pyrethroid LLINs are the standard vector control strategy, and withdrawing them from use is not possible; and are therefore deployed in sequence (with no upper limit for a withdrawal threshold). The IRS insecticides can be rotated at each opportunity. |
| | SEQUENCE IRS: Standard pyrethroid LLINs are the standard vector control strategy, and withdrawing them from use is not possible; and are therefore deployed in sequence (with no upper limit for a withdrawal threshold). The IRS insecticide is deployed in sequence until it reaches the pre-defined withdrawal threshold, at which point it is withdrawn and replaced with the next available IRS insecticide. |

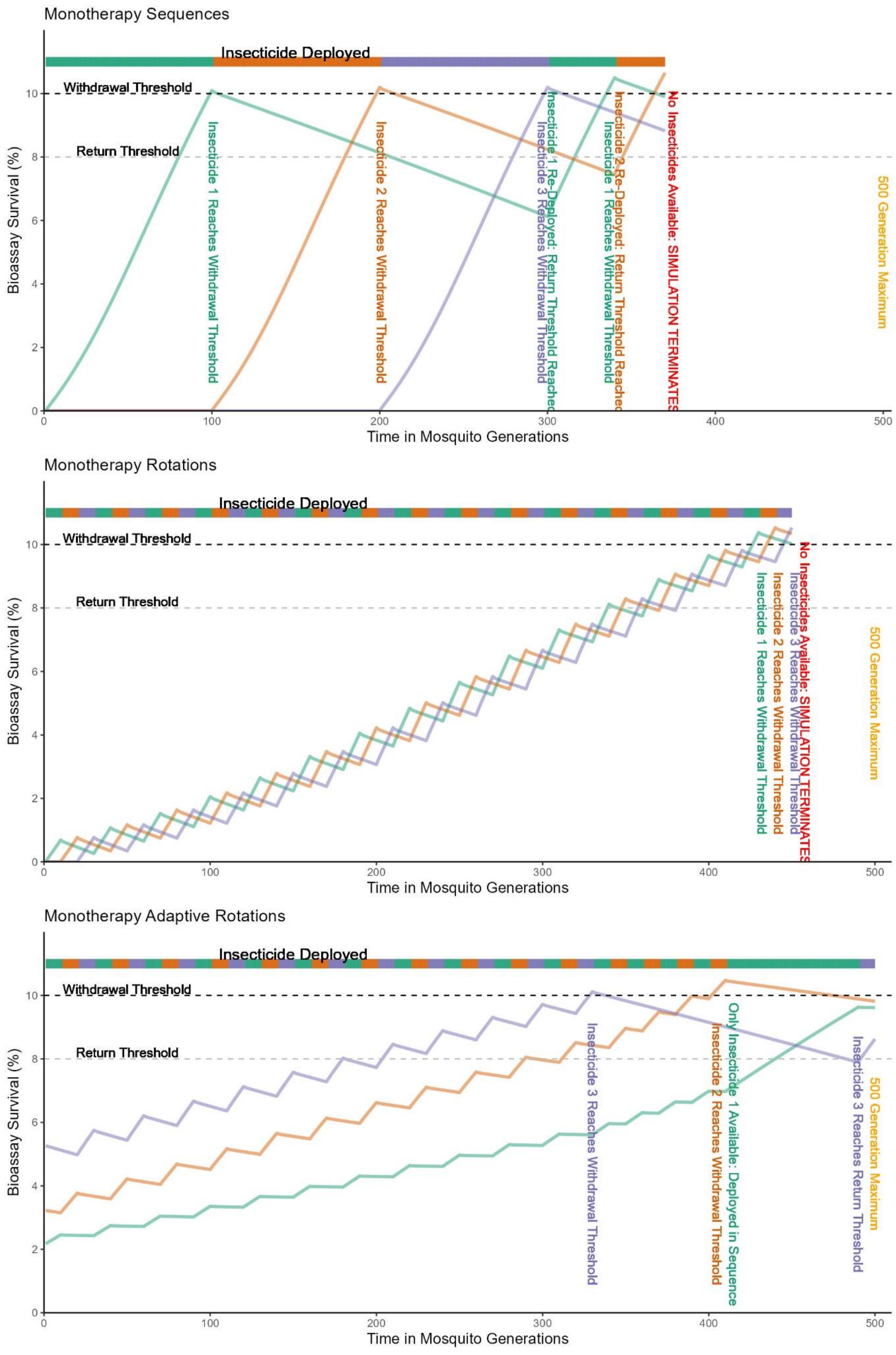
Monotherapy Sequences
Insecticide Deployed
Withdrawal Threshold
Return Threshold
Insecticide 1 Reaches Withdrawal Threshold
Insecticide 2 Reaches Withdrawal Threshold
Insecticide 1 Re-Deployed: Return Threshold Reached
Insecticide 3 Reaches Withdrawal Threshold
Insecticide 2 Re-Deployed: Return Threshold Reached
Insecticide 1 Reaches Withdrawal Threshold
No Insecticides Available: SIMULATION TERMINATES
500 Generation Maximum
Bioassay Survival (%)
Time in Mosquito Generations
Monotherapy Rotations
Insecticide Deployed
Withdrawal Threshold
Return Threshold
Insecticide 1 Reaches Withdrawal Threshold
Insecticide 2 Reaches Withdrawal Threshold
Insecticide 3 Reaches Withdrawal Threshold
No Insecticides Available: SIMULATION TERMINATES
500 Generation Maximum
Bioassay Survival (%)
Time in Mosquito Generations
Monotherapy Adaptive Rotations
Insecticide Deployed
Withdrawal Threshold
Return Threshold
Insecticide 3 Reaches Withdrawal Threshold
Insecticide 2 Reaches Withdrawal Threshold
Only Insecticide 1 Available: Deployed in Sequence
Insecticide 3 Reaches Return Threshold
500 Generation Maximum
Bioassay Survival (%)
Time in Mosquito Generations

**Figure S1.1 Illustrative examples of the monotherapy deployment strategy with switching strategies. Top: Sequences:** Insecticides are deployed until a withdrawal threshold is reached, where the insecticide is withdrawn, and the next insecticide is deployed. If a withdrawn insecticide reaches the return threshold the insecticide is re-available. Simulations terminate when no insecticides are available. **Middle: Rotations:** Insecticide are switched at defined intervals. Simulations terminate when either the same insecticide would be deployed in sequence, or no insecticides are available. **Bottom: Adaptive Rotations:** Insecticides are deployed as rotations (default) but deployed as sequences when required. Simulations end when there are no insecticides available to deploy.

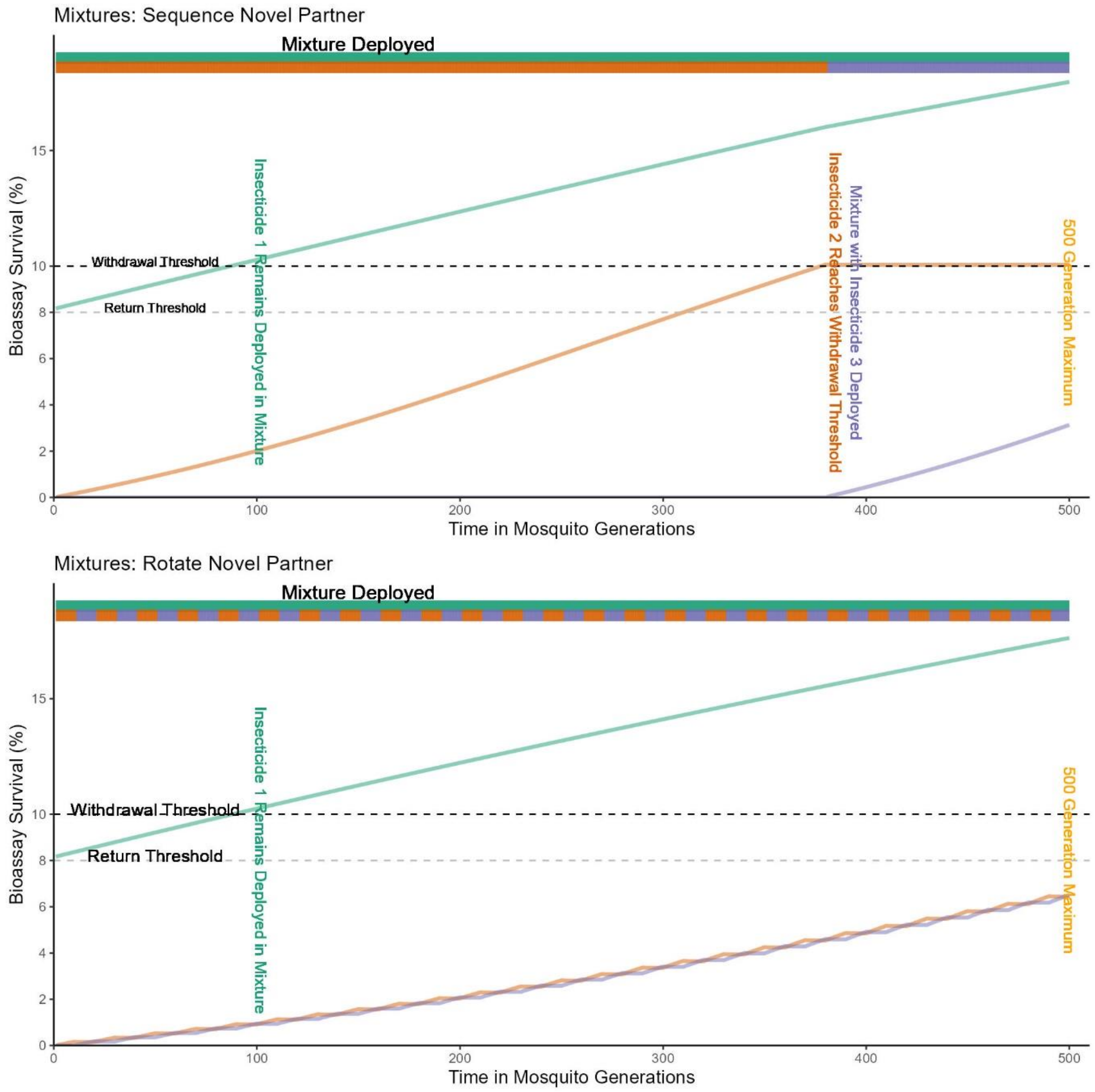


**Figure S1.2 Illustrative example of the mixture deployment strategy with switching decisions on the novel partner. Top: Sequence Novel** Assuming all mixtures share a common partner (i.e., a pyrethroid). The "pyrethroid" insecticide is deployed even after it has exceeded the withdrawal threshold. Deployment decisions are made on the novel partner. **Bottom: rotations novel:** Like with monotherapy rotations, if the mixture of insecticides $i$ & $j$ was to be redeployed immediately (in sequence) then the simulation terminates.

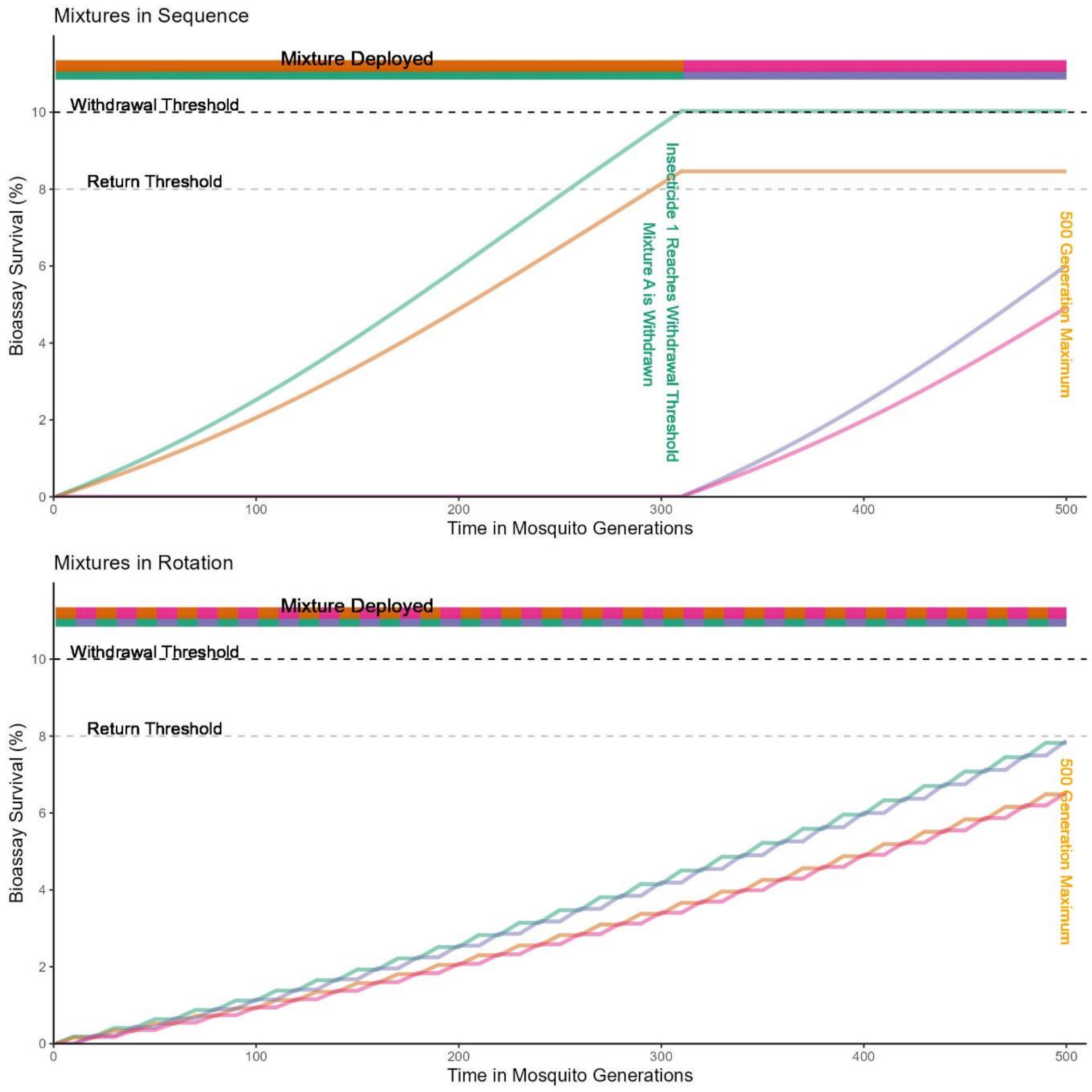


**Figure S1.3 Illustrative example of the mixture deployment strategy with switching decisions on the mixture. Top: Sequence Mixture.** Each mixture formulation is deployed in sequence. Therefore, if either of the insecticides in the mixture reaches the pre-defined withdrawal threshold, then this means the mixture itself is withdrawn. The next mixture formulation in the sequence is then deployed. If a failed insecticide is in multiple mixtures, all those mixture formulations are withdrawn. **Bottom: Rotate Mixture.** The mixture formulation is rotated at each opportunity. For example, a mixture A of insecticides $i$ and $j$ is rotated with mixture B of insecticides $k$ and $m$. Here, the mixture formulation is rotated at each available opportunity. Note, if one of the insecticides in the mixture reaches the withdrawal threshold the mixture is withdrawn from being available for deployment.

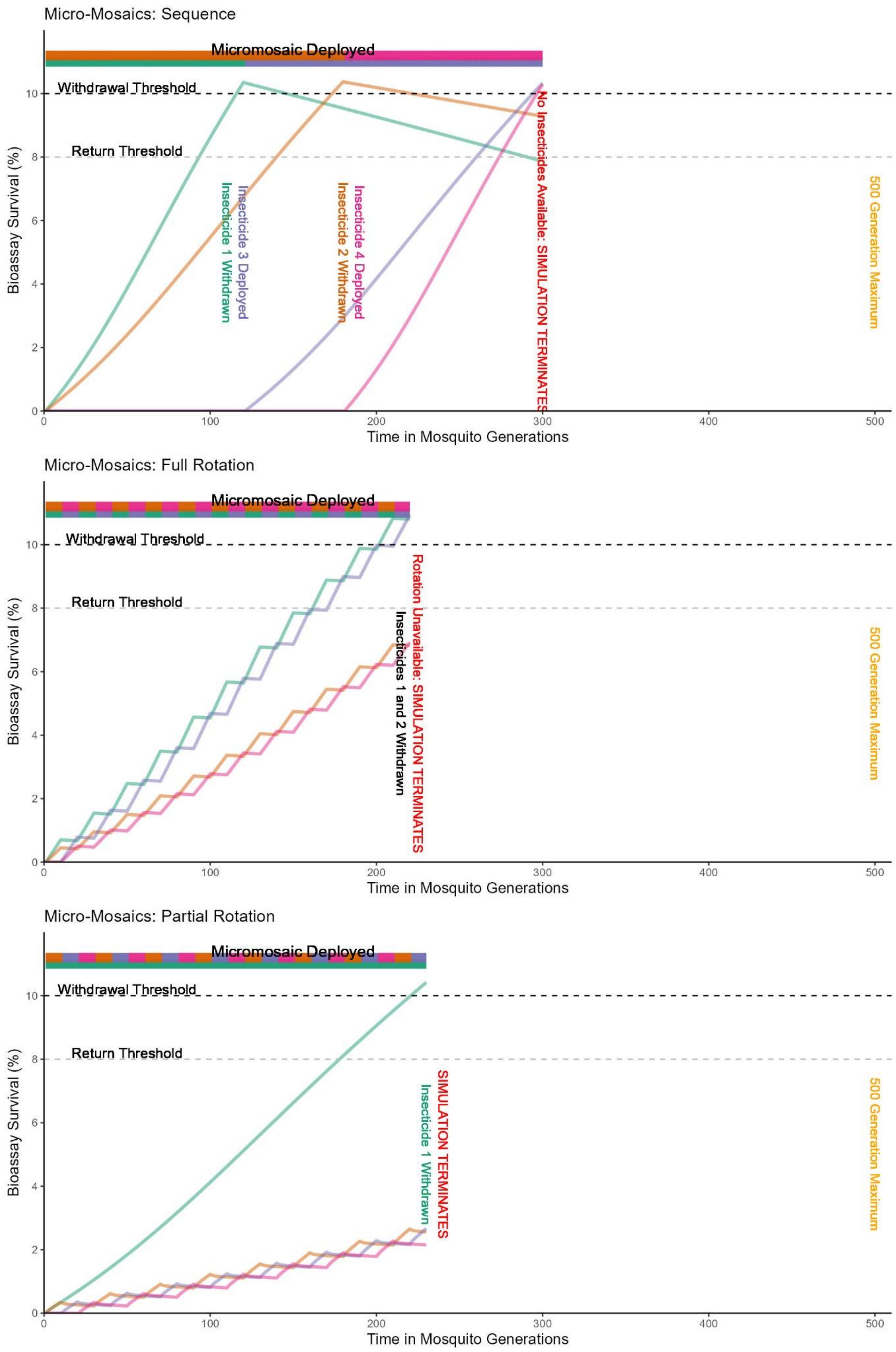

Micro-Mosaics: Sequence
Micromosaic Deployed
Withdrawal Threshold
Return Threshold
Insecticide 1 Withdrawn
Insecticide 3 Deployed
Insecticide 2 Withdrawn
Insecticide 4 Deployed
No Insecticides Available: SIMULATION TERMINATES
500 Generation Maximum
Bioassay Survival (%)
Time in Mosquito Generations
Micro-Mosaics: Full Rotation
Micromosaic Deployed
Withdrawal Threshold
Return Threshold
Insecticides 1 and 2 Withdrawn
Rotation Unavailable: SIMULATION TERMINATES
500 Generation Maximum
Bioassay Survival (%)
Time in Mosquito Generations
Micro-Mosaics: Partial Rotation
Micromosaic Deployed
Withdrawal Threshold
Return Threshold
Insecticide 1 Withdrawn
SIMULATION TERMINATES
500 Generation Maximum
Bioassay Survival (%)
Time in Mosquito Generations

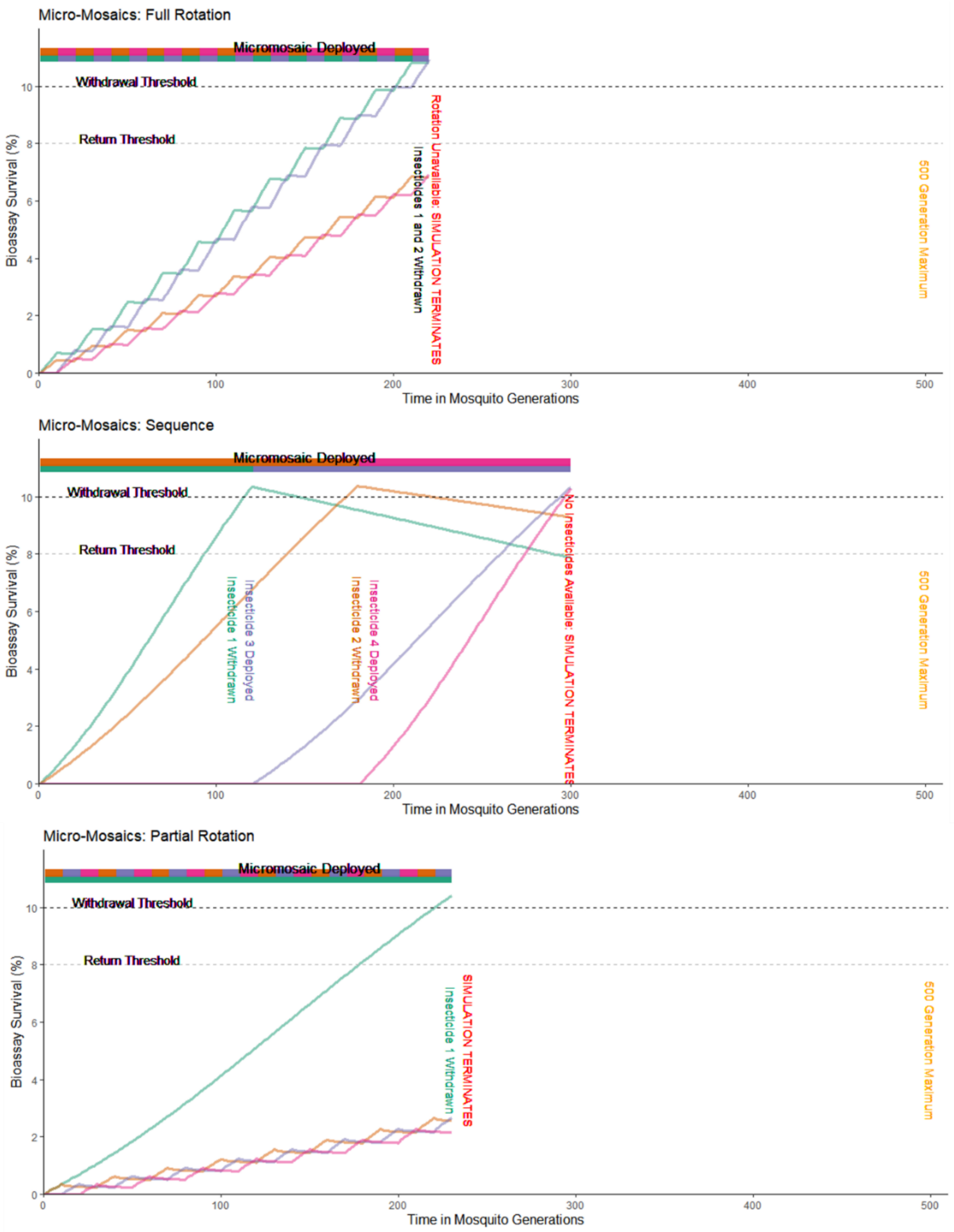


**Figure S1.4 Illustrative example of the micro-mosaic deployment strategy. Top: Individual Sequences:** Insecticides deployed in a micro-mosaic are deployed such that deployment decisions are made on the individual insecticides and can therefore be withdrawn at different times. **Middle: Full Rotation.** Here, both insecticides are rotated out at each deployment opportunity. **Bottom: Partial Rotation.** Here, one of the insecticides remained deployed throughout (e.g., due to being the cheaper insecticide) with the other (potentially more expensive) insecticides rotated.

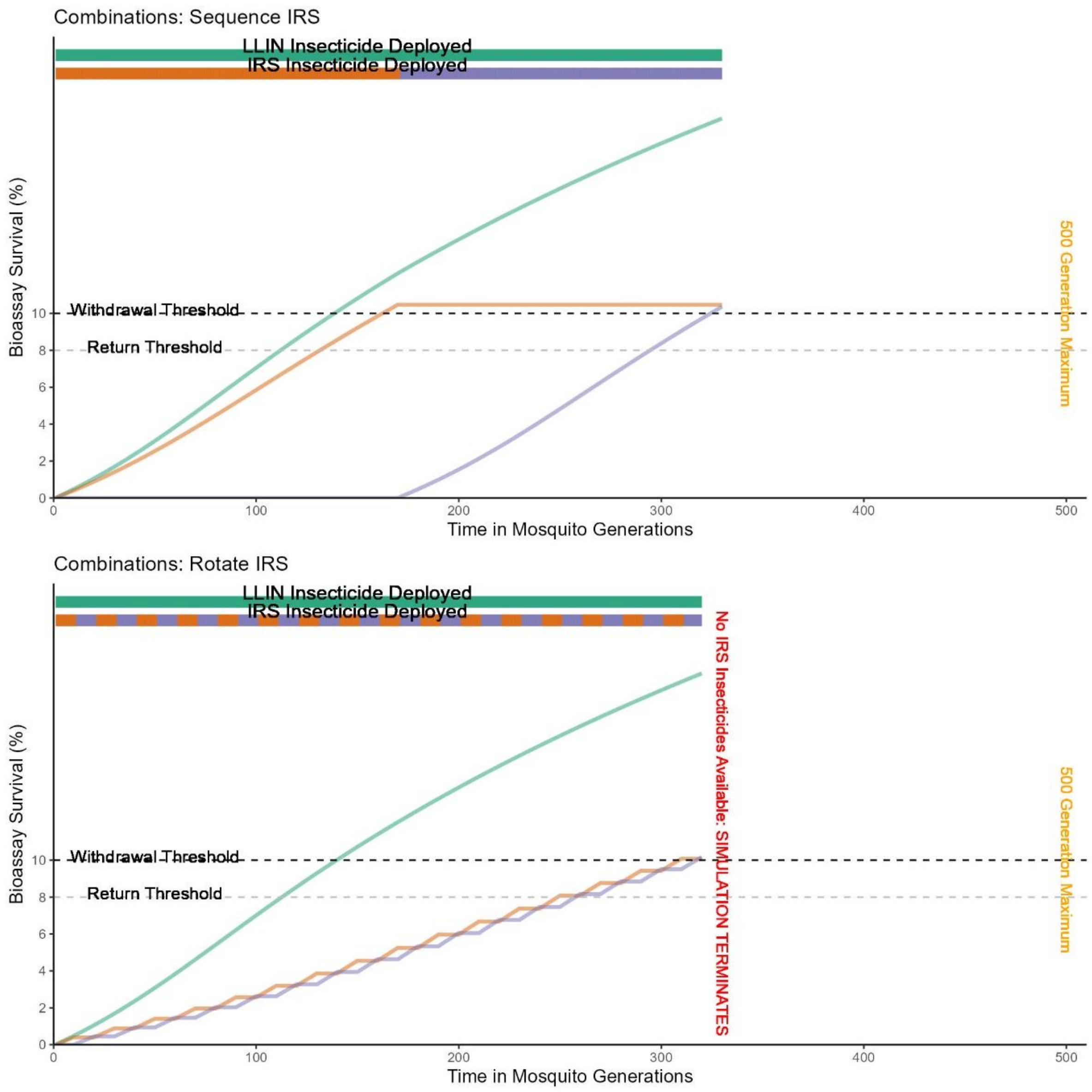


**Figure S1.5 Illustrative examples of the combination deployment strategy with switching decision based on the IRS insecticides. Top: Sequence IRS:** The IRS insecticide is deployed in sequence until it reaches the pre-defined withdrawal threshold, at which point it is withdrawn and replaced with the next available IRS insecticide. The LLIN and IRS can have different deployment intervals (LLINs 3 years, ~30 generations and IRS yearly, ~10 generations). Deployment decisions are only made on the IRS only. **Bottom: Rotate IRS:** The IRS insecticides are rotated at each opportunity, for example yearly.
