## Supplement 2 for "Mathematical Methodology for Dynamic Models of Insecticide Selection Assuming a Polygenic Basis of Resistance"

### Supplement 2: Calibration of the Exposure Scaling Factor (Beta) for “novel” insecticides

As with the “polyres” model, the model is calibrated such, on average, a novel insecticide would be expected to last approximately 10 years under continuous deployment in the absence of fitness costs and refugia (Hobbs et al., 2023). With the dynamic models (“polytruncate” and “polysmooth”) there are two values to calibrate. First, the standard deviation ($\sigma_I$) of the mean PRS ($\bar{z}_I$), and second the exposure scaling factor ($\beta$). The standard deviation for novel insecticides ($\bar{z}_I$=0 to $\bar{z}_I$=100) is input as values as estimated (see Supplement 5), where plausible range of $\sigma_I$ = 20 to $\sigma_I$ = 80. The exposure scaling factor ($\beta$) is a factor converting the selection our desired timescale and is used to account for uncertainty in the value of selection differentials and heritability.

##### Beta and Standard Deviation Calibration Simulations

Parameters were sampled using Latin hypercube sampling (Carnell, 2020). Parameter values were sampled within uniform distributions. A total of 5000 parameter sets for values of female exposure (0.4-0.9), male exposure (0-1) and heritability (0.05 – 0.30 were used. Coverage was set to 1, dispersal is therefore absent (if coverage is 1, there is no refugia) and fitness costs were set to zero. No insecticide decay was allowed, and the insecticide efficacy ($\omega_\tau^i$) remained at 1 for the duration of the simulation. These same 5000 parameter sets were used for each standard deviation and exposure scaling factor permutation, and for both the “polytruncate” and “polysmooth” model. Simulations were run with a single insecticide deployed in sequence. The withdrawal threshold was 10% bioassay survival. The outcome measured was time (in generations) to 10% bioassay survival, defined as the “operational lifespan” of the insecticide.

#### Beta and Standard Deviation Calibration Results

The permutation of exposure scaling factor and standard deviation which gave an average insecticide lifespan of ~10 years (within the 8-12 year range) was found to be standard deviation = 50 and exposure scaling factor = 10 for “polysmooth” (Figure S2.1), and for “polytruncate” was found to be standard deviation = 20 and exposure scaling factor = 1 (Figure S2.2). It should be noted the exposure scaling factor for the “polysmooth” model was identified to be 10, the same value as used for the calibration of the “polyres” model.

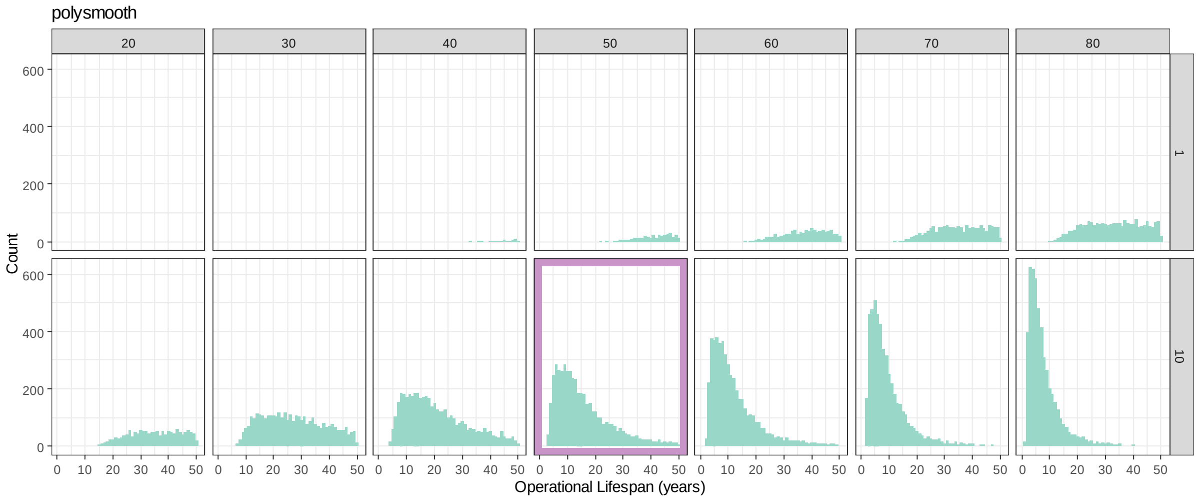


**Figure S2.1 Polysmooth Calibration with Exposure Scaling Factor and Standard Deviation**. Each row of plots indicates the exposure scaling factor used. Each column of the plots indicates the standard deviation used. A standard deviation of 50 and an exposure scaling factor of 10 was found to best calibrate the “polysmooth” model, highlighted in purple.

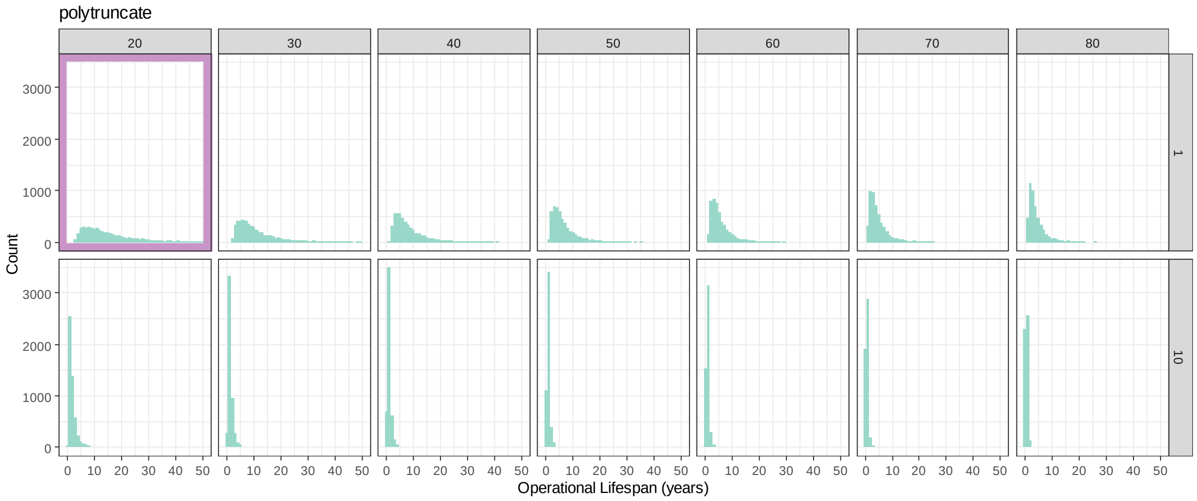


**Figure S2.2 Polytruncate Calibration with Exposure Scaling Factor and Standard Deviation.** The highlighted plot indicates the standard deviation and exposure scaling factor which best calibrated the model. Each row of plots indicates the exposure scaling factor used. Each column of the plots indicates the standard deviation used. A standard deviation of 20 and an exposure scaling factor of 1 was found to best calibrates the "polytruncate" model.

#### References:


Carnell, R. (2020). *lhs: Latin Hypercube Samples* (R package version 1.0.2). https://cran.r-project.org/package=lhs

Hobbs, N., Weetman, D., & Hastings, I. (2023). Insecticide resistance management strategies for public health control of mosquitoes exhibiting polygenic resistance: a comparison of sequences, rotations, and mixtures. *Evolutionary Applications*, *16*(4), 936–959. https://doi.org/DOI: 10.1111/eva.13546
