## Supplement 3 for "Mathematical Methodology for Dynamic Models of Insecticide Selection Assuming a Polygenic Basis of Resistance"

### Supplement 3: Symbols used in the “polysmooth” and “polytruncate” Models

In this supplement we provide detailed descriptions of all symbols used in the mathematical model. Figure S3.1 provides additional detail on what is meant by $F_{z_I}$.

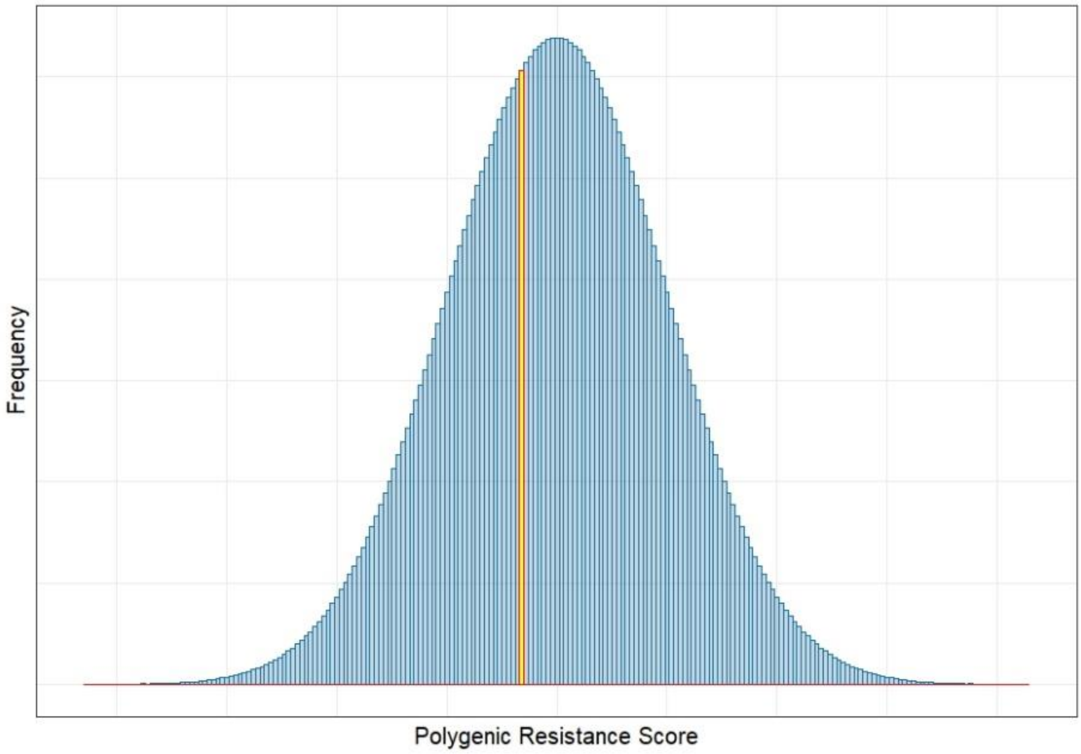


**Figure S3.1 Explanation of what is meant by $F_{z_I}$.** The model code tracks four vectors, two containing the binned values of $z_I^{♀}$ or $z_I^{♂}$ in the Normal distribution, and two vectors containing the corresponding frequencies of the binned values of $z_I^{♀}$ or $z_I^{♂}$ . Therefore, each value of $z_I^{♀}$ or $z_I^{♂}$ is grouped into small bins, of which the precision is dependent upon the length of the vector used in the model code, providing the vector length is long enough, sufficient numerical precision is achieved.

**Table S3.1: Symbols introduced in Methods Section 1.1 to 1.4**

| Symbol | Description | Value/Ranges |
|---|---|---|
| $K_i^B$ | The bioassay survival of a mosquito given a particular polygenic resistance score (PRS). | Internally calculated. |
| $K_{max}$ | The maximum bioassay survival. | 1 |
| $z_I$ | The PRS of a mosquito for trait $I$ which gives resistance to insecticide $i$. | Calculated Internally |
| $z_{50}$ | The polygenic resistance score which gives 50% bioassay survival. | 900 (see Hobbs et al, 2023) |
| $n$ | The slope of the Michaelis-Menten Equation | 1 |
| $K_i^F$ | The field survival of a mosquito given their bioassay survival (dependent on the PRS). | Internally calculated |
| $\varphi_1$ | The regression coefficient between the field (experimental hut) survival and bioassay survival. | 0.48 (Hobbs et al., 2023) |
| $\varphi_2$ | The regression intercept between the field (experimental hut) survival and bioassay survival. | 0.15 (Hobbs et al., 2023) |
| $\zeta_\tau$ | The current concentration of the insecticide. | Internally calculated |
| $\zeta_0$ | The deployed concentration of the insecticide. | Insecticide dependent |
| $k$ | The instantaneous decay rate of the insecticide. | Insecticide dependent |
| $\tau$ | The number of mosquito generations since the insecticide was deployed. | Internally calculated |

| | | |
|---|---|---|
| $\omega_{\tau}^{i}$ | The current efficacy of the insecticide after $\tau$ generations against fully susceptible mosquitoes ($z_I$ < 0) | Internally Calculated |
| $\omega_{0}^{i}$ | The initial deployed efficacy of the insecticide against fully susceptible mosquitoes ($z_I$ < 0). | 1 = full dose, 0.5 = half dose, User input. |
| ${\delta_{\mathrm{b}}}^{i}$ | The base decay rate of the insecticide, where the insecticide generally decays slowly. | Insecticide dependent, user input |
| $\tau_{b}^{i}$ | The number of generation post-deployment at which there is a change in the decay of the insecticide. | Insecticide dependent, user input |
| ${\delta_{\mathrm{r}}}^{i}$ | The rapid decay rate of the insecticide which occurs after the threshold generation has been exceeded. | Insecticide dependent, user input |
| $\varphi_3$ | The regression coefficient of a linear model between the mean PRS from bioassays and the standard deviation from those bioassays. | 0.4 modelled estimate (see Supplement 5) |
| $\varphi_4$ | The regression intercept of a linear model between the mean PRS from bioassays and the standard deviation from those bioassays. | 18 modelled estimate (see Supplement 5) |
| $\sigma_I$ | The standard deviation of trait $I$. | "polytruncate"=20, "polysmooth"=50, or internally calculated |
| ${\sigma_I}^{Int}$ | The standard deviation of trait $I$ in the intervention site. | "polytruncate"=20, "polysmooth"=50, or internally calculated |
| ${\sigma_I}^{Ref}$ | The standard deviation of trait $I$ in the refugia. | "polytruncate"=20, "polysmooth"=50, or internally calculated |

| **Table S3.2: Symbols introduced in Methods Section 2.1 to 2.7** | | |
|---|---|---|
| Symbol | Description | Value/Range |
| $R_I^{S\Phi}$ | The response to selection due to both insecticide selection and fitness costs. | Internally Calculated |
| $R_I^{\Phi}$ | The response to selection due to fitness costs only. | Internally Calculated |
| $h_I^2$ | The narrow sense heritability of the trait, and is therefore the proportion of the selection differential is inherited by the next generation. Which can be unique to each IR trait. | Uniform(0.05 – 0.3) |
| $S_I^{S\Phi}$ | The overall selection differential because of both insecticide selection and fitness costs. | Internally Calculated |
| $\beta$ | A factor which is used to calibrate the model to the desired timescale, and is used to account for parameters not considered in the model. | "polytruncate"=1, "polysmooth"=10 |
| $S_I^{S}$ | The selection differential due to insecticide selection only. | Internally Calculated |
| $S_i^{\Phi}$ | The selection differential due to fitness costs only. | Internally Calculated |
| $S_I^{S\Phi♀}$ | The female selection differential due to both insecticide selection and fitness costs. | Internally Calculated |
| $S_I^{S♀}$ | The female selection differential due to insecticide selection. | Internally Calculated |
| $S_I^{\Phi♀}$ | The female selection differential due to fitness costs. | Internally Calculated |
| $S_I^{S\Phi♂}$ | The male selection differential due to both insecticide selection and fitness costs. | Internally Calculated |
| $S_I^{S♂}$ | The male selection differential due to insecticide selection. | Internally Calculated |
| $S_I^{\Phi♂}$ | The male selection differential due to fitness costs. | Internally Calculated |
| $\bar{z}_I$ | The mean PRS of Trait $I$ which gives resistance to insecticide $i$. | Internally Calculated |
| $F_{z_I}$ | The relative frequency of a single value of $z_I$ in the population. Calculated from the Unit Normal Density Distribution of a value of $z_I$, given $\bar{z}_I$ and $\sigma_I$ | Internally Calculated |
| $\bar{z}_I^{♀}$ | The mean PRS of the female mosquitoes prior to any selection. | Internally Calculated |
| $\bar{z}_I^{P♀}$ | The mean PRS for females of Trait $I$ which gives resistance to insecticide $i$ in the population after insecticide selection. | Internally Calculated |
| $\bar{z}_I^{♂}$ | The mean Polygenic Resistance Score of the male mosquitoes prior to any selection. | Internally Calculated |
| $\bar{z}_I^{P♂}$ | The mean polygenic resistance for males of Trait $I$ which gives resistance to insecticide $i$ in the population after insecticide selection. | Internally Calculated |
| $N^{u♀}$ | The number of female mosquitoes not encountering the insecticide. | Internally Calculated |
| $N_i^{E♀}$ | The number of female mosquitoes encountering and surviving the insecticide. | Internally Calculated |
| $\bar{z}_I^{E♀}$ | The mean PRS of females for Trait $I$ which gives resistance to insecticide $i$ of individuals who survived the insecticide exposure. | Internally Calculated |
| $N^{P♀}$ | The total number of females surviving selection and is therefore the number of females surviving insecticide encounter and the number of females not encountering the insecticide. | Internally Calculated |
| $N^{u♂}$ | The number of males not encountering the insecticide. | Internally Calculated |
| $N_i^{E♂}$ | The number of male mosquitoes encountering and surviving the insecticide. | Internally Calculated |
| $\bar{z}_I^{E♂}$ | The mean PRS of males for Trait $I$ which gives resistance to insecticide $i$ of individuals who survived the insecticide exposure. | Internally Calculated |

| | | |
|---|---|---|
| $N^{P♂}$ | The total number of males surviving selection and is therefore the number of males surviving insecticide encounter and the number of males not encountering the insecticide. | Internally Calculated |
| $N^{T♀}$ | The initial total number of females in the population prior to any selection. | Internally Calculated |
| $x$ | The proportion of female mosquitoes encountering the insecticide. | Uniform(0.4 – 0.9) |
| $N^{T♂}$ | The initial total number of males in the population prior to any selection. | Internally Calculated |
| $m$ | The proportion of male mosquitoes encountering the insecticide as a proportion of the female encounter rate. | Uniform(0-1) |
| $S_I^{E♀}$ | The within generation change in the mean PRS of female mosquitoes exposed to the insecticide and survive and their mean PRS before the encounter. | Internally Calculated |
| $S_I^{E♂}$ | The within generation change in the mean PRS of male mosquitoes exposed to the insecticide and survive and their mean PRS before the encounter. | Internally Calculated |
| $F_{z_I^{E♀}}$ | The frequency of values of $z_I$ of female mosquitoes who have been exposed and survived the insecticide. | Internally Calculated |
| $F_{z_I^{E♂}}$ | The frequency of values of $z_I$ of male mosquitoes who have been exposed and survived the insecticide. | Internally Calculated |
| $N_{ij}^{E}$ | The number of individuals encountering the mixture and surviving the exposure. | Internally Calculated |
| $N_{ij}^{E♀}$ | The number of female mosquitoes encountering the mixture and surviving the exposure | Internally Calculated |
| $N_{ij}^{E♂}$ | The number of male mosquitoes encountering the mixture and surviving the exposure | Internally Calculated |
| $\bar{K}_i^F$ | The mean field survival of the mosquito population to insecticide $i$. | Internally Calculated |
| $\bar{K}_j^F$ | The mean field survival of the mosquito population to insecticide $j$. | Internally Calculated |
| $\bar{z}_I^{E♀}$ | The mean PRS of females for Trait $I$ of individuals who survived the insecticide exposure. | Internally Calculated |
| $\phi$ | The fitness cost of the trait, as a proportion of the standard deviation. | Uniform(0.05 to 0.2) |
| $\phi^{♀}$ | The fitness cost of the trait for females, as a proportion of the standard deviation. | Uniform(0.05 to 0.2). |
| $\phi^{♂}$ | The fitness cost of the trait for males, as a proportion of the standard deviation. | Uniform(0.05 to 0.2) |
| $\bar{z}_I^{Int}$ | The initial mean PRS at the start of the generation in the intervention site. | Internally calculated |
| $\bar{z}_I^{Int'}$ | The mean PRS of the eggs which the females will lay from the intervention site. | Internally calculated |
| $\alpha_{\Gamma I}$ | The degree of genetic correlation and the amount of cross resistance between trait $\Gamma$ and trait $I$. | -1 to 1. |
| $\bar{z}_I^{Ref}$ | The initial mean PRS at the start of the generation in the refugia. | Internally calculated |
| $\bar{z}_I^{Ref'}$ | The mean PRS of the eggs which the females will lay from the refugia. | Internally calculated |
| $r_{Int}$ | The number of mosquitoes migrating from the intervention site to the refugia. | Internally calculated |
| $r_{Ref}$ | The number of mosquitoes migrating from the refugia site to the intervention site. | Internally calculated |
| $\theta$ | The dispersal rate of the populations. | Uniform(0.1 – 0.9) |
| $C$ | The proportion of the total population of mosquitoes residing in the intervention site. | Uniform(0.1 - 0.9) |
| $\bar{z}_I^{Int''}$ | The mean PRS of eggs laid in the intervention site, and is therefore the mean of the next generation in the intervention. | Internally calculated |
| $\bar{z}_I^{Ref''}$ | The mean PRS of eggs laid in the refugia, and is therefore the mean of the next generation in the refugia. | Internally calculated |

| **Table S3.3 Symbols introduced in Methods Section 3.1** | | |
|---|---|---|
| Symbol | Description | Value/Range |
| $c_i$ | The proportion of the coverage where a house is treated with only insecticide $i$. | 0-1 |
| $c_j$ | The proportion of the coverage where a house is treated with only insecticide $j$. | 0-1 |
| $c_{ij}$ | The proportion of the coverage where a house is treated with both insecticide $i$ and insecticide $j$. | 0-1 |
| $\Lambda_{i\|ij}$ | The probability of encountering only insecticide $i$ given the mosquito entered a house with both insecticide $i$ and $j$. | 0-1 |
| $\Lambda_{j\|ij}$ | The probability of encountering only insecticide $j$ given the mosquito entered a house with both insecticide $i$ and $j$. | 0-1 |
| $\Lambda_{ij\|ij}$ | The probability of encountering both insecticide $i$ and $j$ given the mosquito entered a house with both insecticide $i$ and $j$. | 0-1 |
| $\Lambda^{♀}_{i\|ij}$ | The probability of encountering only insecticide $i$ given the mosquito entered a house with both insecticide $i$ and $j$ for female mosquitoes. | 0-1 |
| $\Lambda^{♀}_{j\|ij}$ | The probability of encountering only insecticide $j$ given the mosquito entered a house with both insecticide $i$ and $j$ for female mosquitoes. | 0-1 |
| $\Lambda^{♀}_{ij\|ij}$ | The probability of encountering both insecticide $i$ and $j$ given the mosquito entered a house with both insecticide $i$ and $j$ for female mosquitoes. | 0-1 |
| $\Lambda^{♂}_{i\|ij}$ | The probability of encountering only insecticide $i$ given the mosquito entered a house with both insecticide $i$ and $j$ for male mosquitoes. | 0-1 |
| $\Lambda^{♂}_{j\|ij}$ | The probability of encountering only insecticide $j$ given the mosquito entered a house with both insecticide $i$ and $j$ for male mosquitoes. | 0-1 |
| $\Lambda^{♂}_{ij\|ij}$ | The probability of encountering both insecticide $i$ and $j$ given the mosquito entered a house with both insecticide $i$ and $j$ for male mosquitoes. | 0-1 |
| $\rho$ | The proportion of female mosquitoes who laid eggs in the previous gonotrophic cycle starting the next gonotrophic cycle. | Internally calculated |
| $d$ | The daily "natural" survival probability in the absence of insecticides. | 0.8 [default] (Matthews et al., 2020) |
| $g$ | The length of the gonotrophic cycle in days. | 3 days [default] |

| **Table S3.4 Symbols introduced in Methods Sections 3.2 to 3.3** | | |
|---|---|---|
| Symbol | Description | Value/Range |
| $S^{S♀}_{I(G)}$ | The selection differential as a result of insecticide selection in females in gonotrophic cycle $G$. | Internally calculated |
| $\bar{z}^{P♀}_{I\,(G)}$ | The mean polygenic resistance for females of Trait $I$ which gives resistance to insecticide $i$ in the population after insecticide selection in gonotrophic cycle $G$. | Internally calculated |
| $\bar{z}^{♀}_{I\,(G=0)}$ | The mean Polygenic Resistance Score of the female mosquitoes prior to any selection and therefore before the start of any gonotrophic cycles. | Internally calculated |
| $S^{S\Phi♀}_{I(G)}$ | The selection differential as a result of insecticide selection and fitness costs in females in gonotrophic cycle $G$. | Internally calculated |
| $R^{S\Phi}_{I(G)}$ | The response (as a result of insecticide selection and fitness costs) for trait $I$ in gonotrophic cycle $G$. | Internally calculated |
| $N_o$ | The total number of oviposition events by female mosquitoes in a single mosquito generation. | Internally calculated |
| $R^T_I$ | The total overall response for the mosquito generation. And is therefore the average between generation change. | Internally calculated |
| $\bar{z}^{P♀}_{I(G)}$ | The mean polygenic resistance for females of Trait $I$ which gives resistance to insecticide $i$ in the population after insecticide selection in gonotrophic cycle $G$. | Internally calculated |
| $N^{S♀}_{(G)}$ | The total number of females surviving selection and is therefore the number of females surviving insecticide encounter and the number of females not encountering the insecticide in gonotrophic cycle $G$. | Internally calculated |
| $N^{u♀}_{(G)}$ | The number of female mosquitoes not encountering the insecticide in gonotrophic cycle $G$. | Internally calculated |
| $\bar{z}^{E♀}_{I(G)}$ | The mean PRS of females for Trait $I$ which gives resistance to insecticide $i$ of individuals who survived the insecticide exposure in gonotrophic cycle $G$. | Internally calculated |
| $N^{P♀}_{(G)}$ | The number of female mosquitoes encountering and surviving the insecticide in gonotrophic cycle $G$ | Internally calculated |
| $F_{z^{♀}_{I(G)}}$ | The frequency of values of $z_I$ for all female mosquitoes in gonotrophic cycle $G$. | Internally calculated |
| $F_{z^{E♀}_{I(G)}}$ | The relative frequency of values of $z_I$ of female mosquitoes who have encountered and survived the insecticide in gonotrophic cycle $G$. | Internally calculated. |

| **Table S3.5 Symbols introduced in Methods Section 3.4** | | |
|---|---|---|
| Symbol | Description | Value/Range |
| $\bar{z}_{I\ (G=0)}^{Ref♀}$ | The mean PRS of female mosquitoes emerging in the refugia prior to any selection. | Internally calculated |
| $F_{z_{I\,Ref\,(G)}^{Ref♀}}$ | The relative frequency of a value of $z_I$ for female mosquitoes who emerged in the refugia and lay eggs in the refugia in the gonotrophic cycle $G$. | Internally calculated |
| $F_{z_{I\,Ref\,(G)}^{Int♀}}$ | The relative frequency of a value of $z_I$ for female mosquitoes who emerged in the intervention site and lay eggs in the refugia in gonotrophic cycle $G$. | Internally calculated |
| $\bar{z}_{I\,Ref\,(G)}^{S\,Ref♀}$ | The mean PRS of female mosquitoes which emerged in the refugia and laying eggs in the refugia in the gonotrophic cycle $G$. | Internally calculated |
| $\bar{z}_{I\,Ref\,(G)}^{S\,Int♀}$ | The mean PRS of female mosquitoes which emerged in the intervention site and laying eggs in the refugia in the gonotrophic cycle $G$. | Internally calculated |
| $N_{Ref\,(G)}^{Ref♀}$ | The number of female mosquitoes which emerged (and mated) in the refugia laying eggs in the refugia in gonotrophic cycle $G$. | Internally calculated |
| $N_{Ref\,(G)}^{Int♀}$ | The number of female mosquitoes which emerged (and mated) in the intervention site laying eggs in the refugia in gonotrophic cycle $G$. | Internally calculated |
| $S_{I\,Ref\,(G)}^{S\,Ref♀}$ | The insecticide selection differential of female mosquitoes who emerged in the refugia and lay eggs in the refugia in gonotrophic cycle $G$. | Internally calculated |
| $S_{I\,Ref\,(G)}^{S\,Int♀}$ | The insecticide selection differential of female mosquitoes who emerged in the intervention site and lay eggs in the refugia in gonotrophic cycle $G$. | Internally calculated |
| $R_{I\,Ref\,(G)}^{Ref}$ | The response for trait $I$ of eggs laid by female mosquitoes which emerged (and mated) in the refugia and lay eggs in the refugia in gonotrophic cycle $G$. | Internally calculated |
| $R_{I\,Ref\,(G)}^{Int}$ | The response for trait $I$ of eggs laid by female mosquitoes which emerged (and mated) in the intervention site and lay eggs in the refugia in gonotrophic cycle $G$. | Internally calculated |
| $F_{z_{I\,Int\,(G)}^{Ref♀}}$ | The frequency of a value of $z_I$ for female mosquitoes who emerged in the refugia and lay eggs in the intervention site in gonotrophic cycle $G$. | Internally calculated |
| $F_{z_{I\,Int\,(G)}^{Int♀}}$ | The frequency of a value of $z_I$ for female mosquitoes who emerged in the intervention site and lay eggs in the intervention site in gonotrophic cycle $G$. | Internally calculated |
| $\bar{z}_{I\,Int\,(G)}^{P\,Ref♀}$ | The mean PRS of female mosquitoes which emerged in the refugia and laying eggs in the intervention site in the gonotrophic cycle $G$. | Internally calculated |
| $\bar{z}_{I\,Int\,(G)}^{P\,Int♀}$ | The mean PRS of female mosquitoes which emerged in the intervention site and laying eggs in the intervention site in the gonotrophic cycle $G$. | Internally calculated |
| $S_{I\,Int\,(G)}^{S\,Ref♀}$ | The insecticide selection differential of female mosquitoes who emerged in the refugia and lay eggs in the intervention site in gonotrophic cycle $G$. | Internally calculated |
| $S_{I\,Int\,(G)}^{S\,Int♀}$ | The insecticide selection differential of female mosquitoes who emerged in the intervention site and lay eggs in the intervention site in gonotrophic cycle $G$. | Internally calculated |
| $N_{Int\,(G)}^{Ref♀}$ | The number of female mosquitoes which emerged (and mated) in the refugia laying eggs in the intervention site in gonotrophic cycle $G$. | Internally calculated |
| $N_{Int\,(G)}^{Int♀}$ | The number of female mosquitoes which emerged (and mated) in the intervention site laying eggs in the intervention site in gonotrophic cycle $G$. | Internally calculated |
| $R_{I\,Int\,(G)}^{Ref}$ | The response for trait $I$ of eggs laid by female mosquitoes which emerged (and mated) in the refugia and lay eggs in the intervention site in gonotrophic cycle $G$. | Internally calculated |
| $R_{I\,Int\,(G)}^{Int}$ | The response for trait $I$ of eggs laid by female mosquitoes which emerged (and mated) in the intervention site and lay eggs in the intervention site in gonotrophic cycle $G$. | Internally calculated |
| $N_{o\,Ref}^{Total♀}$ | The total number of oviposition events in the refugia across all gonotrophic cycles. | Internally calculated |
| $N_{o\,Ref}^{Ref♀}$ | The total number of oviposition events in the refugia across all gonotrophic cycles for females which had originally emerged (and mated) in refugia. | Internally calculated |
| $N_{o\,Ref}^{Int♀}$ | The total number of oviposition events in the refugia across all gonotrophic cycles for females which had originally emerged (and mated) in the intervention site. | Internally calculated |
| $N_{o\,Int}^{Total♀}$ | The total number of oviposition events in the intervention site across all gonotrophic cycles. | Internally calculated |
| $N_{o\,Int}^{Int♀}$ | The total number of oviposition events in the intervention site across all gonotrophic cycles for females which had originally emerged (and mated) in the intervention site. | Internally calculated |
| $N_{o\,Int}^{Ref♀}$ | The total number of oviposition events in the intervention site across all gonotrophic cycles for females which had originally emerged (and mated) in the refugia. | Internally calculated |
| $R_{I\,Int}^{T\,Int}$ | The average overall response for trait $I$ of eggs laid in the intervention site across all gonotrophic cycles by females which had originally emerged (and mated) in the intervention site. | Internally calculated |
| $R_{I\,Int}^{T\,Ref}$ | The average overall response for trait $I$ of eggs laid in the intervention site across all gonotrophic cycles by females which had originally emerged (and mated) in the refugia. | Internally calculated |
| $R_{I\,Ref}^{T\,Ref}$ | The average overall response for trait $I$ of eggs laid in the refugia across all gonotrophic cycles by females which had originally emerged (and mated) in the refugia. | Internally calculated |

| | | |
|---|---|---|
| $R_{I\ Ref}^{T\ Int}$ | The average overall response for trait $I$ of eggs laid in the refugia across all gonotrophic cycles by females which had originally emerged (and mated) in the intervention site. | Internally calculated |

#### References


Matthews, J., Bethel, A., & Osei, G. (2020). An overview of malarial Anopheles mosquito survival estimates in relation to methodology. *Parasites and Vectors*, *13*(1), 1–12. https://doi.org/10.1186/s13071-020-04092-4

Hobbs, N., Weetman, D., & Hastings, I. (2023). Insecticide resistance management strategies for public health control of mosquitoes exhibiting polygenic resistance: a comparison of sequences, rotations, and mixtures. *Evolutionary Applications*, *16*(4), 936–959. https://doi.org/DOI: 10.1111/eva.13546
