## Supplement 4 for "Mathematical Methodology for Dynamic Models of Insecticide Selection Assuming a Polygenic Basis of Resistance"

### Supplement 4: Describing and Defining the Quantification of Resistance

#### Quantifying Resistance - Polygenic Resistance Scores

A primary challenge for quantitative traits is having a measurable scale for the trait. Classically, in the quantitative genetics and selective breeding literature the trait of interest is clearly defined and directly measurable (for example, weight, wing length, milk yield). Conversely, measuring the “amount” of IR in an individual mosquito or a population of mosquitoes is not clearly defined as it can be measured is several contexts. In IR surveillance, the level of IR in a population of mosquitoes is regularly measured using standardised bioassays such as WHO tube bioassay or CDC bottle bioassay. The WHO tube bioassay is more commonly used due to its ease in conducting and interpretation. In the WHO tube bioassay female mosquitoes (aged 3-5 days) are exposed to a fixed concentration of insecticide for a fixed time-period (WHO, 2022). This constitutes two definitions and IR scales for surveillance i.e.one for the WHO cylinder and one for the CDC bioassays. For selection and evolution of IR the better definition would be the ability to survive insecticide contact under realistic field conditions but there are many definitions of what may be “realistic” and hence an almost infinite number of different scales for measuring IR.

We therefore constructed an arbitrary scale of IR (Hobbs et al., 2023) using the Hill-variant of the Michaelis-Menten equation (where $n$=1) to convert an “amount of IR” ($z_I$, the value of the Polygenic Resistance Score (PRS)) to insecticide $i$ to the measured survival to insecticide $i$ in a bioassay ($K_i^B$).

$$K_i^B = \frac{K_{max} * {z_I}^n}{z_{50} + {z_I}^n}$$

Equation 2a

$z_{50}$ is the PRS giving 50% bioassay survival. $K_{max}$ is the maximum proportion of mosquitoes surviving in a bioassay, which is, by definition, 1. The choice of what value to use $z_{50}$ is user defined, with the choice of $z_{50}$ = 900 being based on having $z_I$=100 has 10% bioassay survival, a commonly suggested criterion for confirmed resistance (WHO, 2018). All parameterisation and calculation were conducted with $z_{50}$ = 900. A graphical representation of the relationship between PRS and bioassay survival is given in

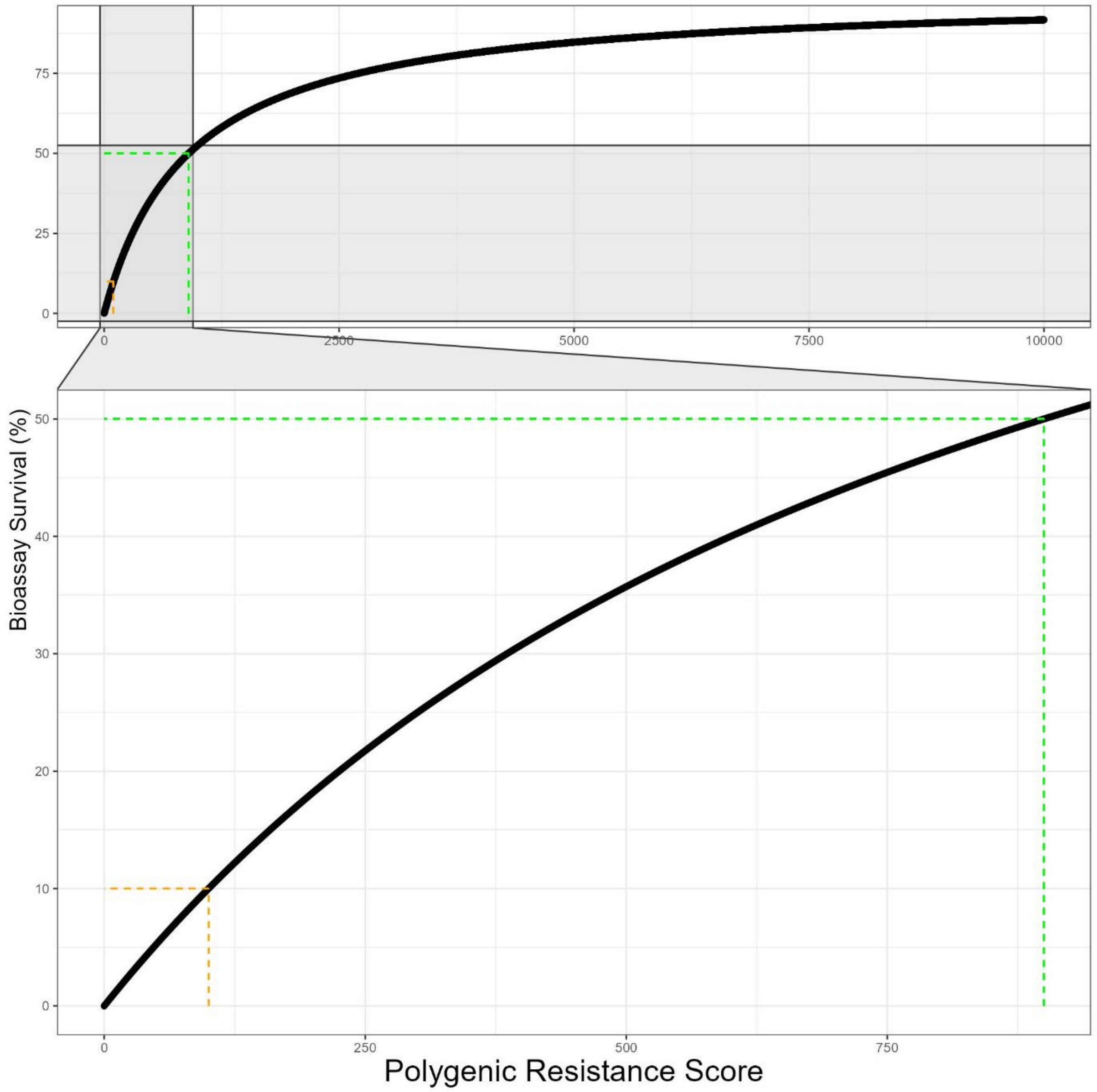


Figure .1 (with a $z_{50}$ = 900).

**Converting Bioassay Survival to Field Survival**

The model tracks the mean PRS in a mosquito population over discrete generations. The PRS can be converted to bioassay survival as this information would be used to inform operational decisions and is how IR is measured in the field. However, bioassay survival is not field survival. Bioassays for measuring IR are highly standardised using 2-5 day old non-bloodfed females, with mortality assessed against a fixed concentration of insecticide for a fixed time-period in a highly standardised process

(WHO, 2018). Whereas the field exposure to insecticides is a far from standardised process, with both contact durations and insecticide concentrations likely to vary substantially between each insecticide/mosquito encounter. Mosquito mortality, for females only, in bioassays has been found to be correlated with survival in experimental huts (Churcher et al., 2016). Therefore, we can model the relationship between bioassay survival and experimental hut survival, allowing us to convert the bioassay survival (given a PRS) into the corresponding experimental hut survival, which we use as a suitable approximation for field survival.

$$K_i^F = \varphi_1 K_i^B + \varphi_2$$

Equation 2b

Where $K_i^F$ is the survival to insecticide $i$ under field conditions. $K_i^B$ is the bioassay survival to insecticide $i$, given a PRS value calculated from equation 1a. $\varphi_1$ and $\varphi_2$ are regression coefficients obtained from linear modelling, $\varphi_1$ = 0.48 and $\varphi_2$ = 0.15 (see Hobbs et al., 2022). The relationship between bioassay survival and field survival, and the relationship between PRS and field survival is presented in Figure S4.1.

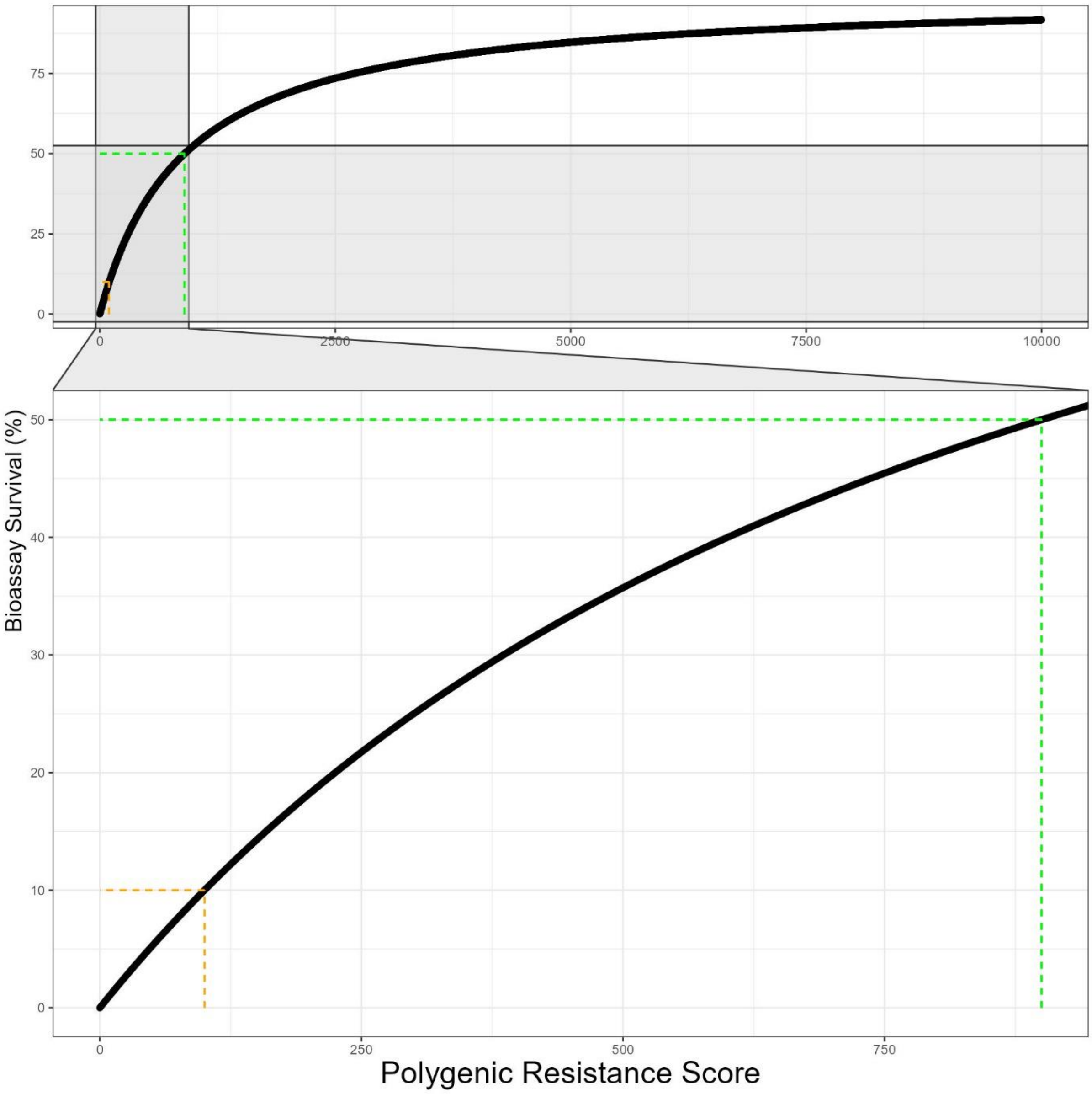


**Figure S4.1: The Polygenic Resistance Score (PRS) relationship with Bioassay Survival.** The top graph shows the PRS on a scale of 0 to 10000. The lower graph shows the same data on a scale from 0 and 1000 which are the values especially important in the evaluation of resistance to novel insecticides; the orange dotted line indicates an indicative 10% bioassay survival (PRS = 100) threshold which is our default for withdrawing an insecticide. The green dotted line is our $z_{50}$ value, which has been set at 900.

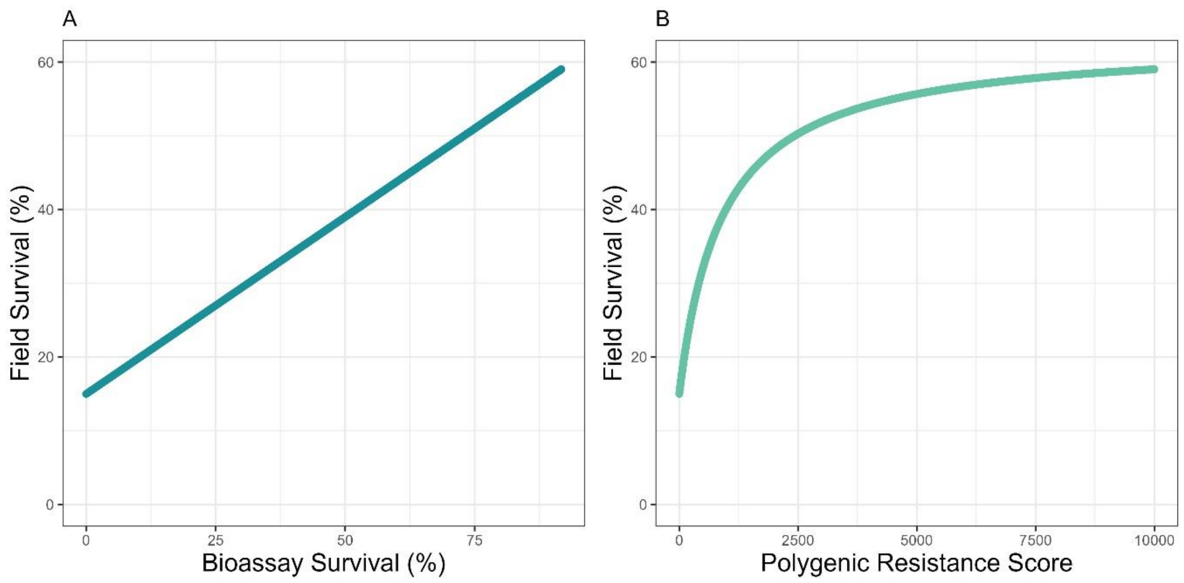


**Figure S4.1: The relationship between bioassay survival and field survival.** The left-hand graph shows the relationship between bioassay survival and field survival as estimated from the linear model using $\varphi_1$=0.48 and $\varphi_2$=0.15. The right-hand graph shows the updated relationship between the Polygenic Resistance Score (PRS) and field survival once converted via the bioassay survival.

#### References


Churcher, T. S., Lissenden, N., Griffin, J. T., Worrall, E., & Ranson, H. (2016). The impact of pyrethroid resistance on the efficacy and effectiveness of bednets for malaria control in Africa. *ELife*, *5*(AUGUST), 1–26. https://doi.org/10.7554/eLife.16090

Hobbs, N., Weetman, D., & Hastings, I. (2023). Insecticide resistance management strategies for public health control of mosquitoes exhibiting polygenic resistance: a comparison of sequences, rotations, and mixtures. *Evolutionary Applications*, *16*(4), 936–959. https://doi.org/DOI: 10.1111/eva.13546

WHO. (2018). Test procedures for insecticide resistance monitoring in malaria vector mosquitoes (Second edition) (Updated June 2018). In *Who*. http://www.who.int/malaria/publications/atoz/9789241511575/en/

World Health Organization. (2022). *Manual for monitoring insecticide resistance in mosquito vectors and selecting appropriate interventions*. World Health Organisation. https://www.who.int/publications/i/item/9789240051089
