## Supplement 5 for "Mathematical Methodology for Dynamic Models of Insecticide Selection Assuming a Polygenic Basis of Resistance"

### Supplement 5: Fixed and Variable Standard Deviations

This supplement explains the rationale behind allowing the standard deviation to be fixed or to scale with the population mean value of the Polygenic Resistance Score (PRS) (Methods Section 1.4). And provides details of the estimation of the value of the population standard deviation ($\sigma_I$).

**Scenario 1: standard deviation ($\sigma_I$) is fixed:**

Mathematical models of IRM have focused mainly on the deployment of insecticides to which there is initially little resistance with the aim to identify strategies which can maintain low levels of resistance. For the "polysmooth" and "polytruncate" models, this would generally limit the value of $\bar{z}_I$ to 100, corresponding to 10% bioassay survival, our default defition of a "failing insecticide". It would seem intuitive to keep $\sigma_I$ fixed when $\bar{z}_I$ remains low. This additionally allows simple exploration of the implications of having a highly variable starting population versus a less variable population (i.e., comparing high versus low $\sigma_I$). Additionally calibrating models (e.g. the exposure scaling factor ($\beta$), see Supplement 2) in Equation 3c to our expected timescales (i.e., a novel insecticide on average has a lifespan of 10 years (see Hobbs et al., 2023 for details) is simpler when using fixed standard deviation.

**Scenario 2: standard deviation ($\sigma_I$) varies:**

Not all changes in Polygenic Resistance Score (PRS) correspond to the same unit change in bioassay survival, as the relationship between PRS and bioassay survival is curved (Supplement 4, Figure S4.1), suggesting $\sigma_I$ may need to increase with an increasing $\bar{z}_I$. For example if $\sigma_I$ = 30, this is a comparatively large value at low PRS (e.g., $\bar{z}_I$ is in the range 0-100) but a very small value at high PRS values (e.g., $\bar{z}_I$ is in the range 3600-8100). Therefore $\sigma_I$ needs to change with respect to $\bar{z}_I$ (see Figure S2 Supplement 1) to account for this. A linear model of $\sigma_I \sim \bar{z}_I$ was used to estimate this, using WHO tube bioassay results from the field and the associated standard deviation of those bioassay results (see Supplement 1 for details).

$$\sigma_I = \varphi_3 \bar{z}_I + \varphi_4$$

Equation 1e

Where $\varphi_3$ and $\varphi_4$ are the regression coefficient and regression intercept respectively of a linear model. If the intervention site and refugia have different $\bar{z}_I$, then site specific standard deviations can be calculated as:

$$\sigma_I^{Int} = \varphi_3 \bar{z}_I^{Int} + \varphi_4$$

Equation 1e(Int)

$$\sigma_I^{Ref} = \varphi_3 \bar{z}_I^{Ref} + \varphi_4$$

Equation 1e(Ref)

**Estimating the Population Standard Deviation ($\sigma_I$) of the Mean Polygenic Resistance Score ($\bar{z}_I$) from Field Data**

The “polytruncate” and “polysmooth” models require a manual input of the standard deviation of the mean PRS of the population ($\sigma_I$). The first step is to therefore calculate the standard deviation (of the PRS) of WHO bioassay results. This required identifying papers which reported both the mean bioassay mortality and the associated 95% confidence interval of this estimate. It should be noted the WHO dataset of bioassay results (the Malaria Threat Map) does not include the 95% confidence interval, and only includes the point estimate of the bioassay mortality.

For inclusion publications were required to meet the requirements of: 1). reporting the mean bioassay mortality; 2). reporting the 95% confidence interval (CI) of the estimate; 3). Reporting the number of bioassays or number of mosquitoes. For publications where the number of mosquitoes was given instead of the number of bioassays, we estimated the expected number of bioassays as number of mosquitoes divided by 25; and rounding to the nearest who number. The number of mosquitoes was divided by 25 as this is the recommended number of mosquitoes per cylinder bioassay (WHO, 2018). Data meeting these requirements was obtained and extracted from WHO cylinder bioassay results from Uganda (Thomsen et al., 2014), Democratic Republic of the Congo (Wat'Senga et al., 2018) and Ethiopia (Alemayehu et al., 2017). Data was from *Anopheles gambiae s.l.* and *Anopheles arabiensis* and included a wide variety of insecticides.

The reported mean bioassay mortality and 95% CI to the corresponding bioassay survival (survival = 1 - mortality). These survival values were then converted to the PRS (using Equation 1a). With the values now as PRS values the standard deviation of the PRS can then be calculated:

$$\sigma_i = \sqrt{N} * (Upper\ 95\%\ CI\ - Lower\ 95\%CI)/3.92$$

Equation S1

Where $N$ is the number of WHO cylinder bioassays conducted (or estimated to have been conducted).

One of the major purposes of the model is to evaluate how to deploy novel (or near novel) insecticides to extend their operational lifespan. Therefore, we restricted the bioassay survival to include only occasions where the bioassay survival was less than 10%. Plausible values for the standard deviation (of the PRS) for novel (or novel-like) insecticides (i.e., have a mean PRS ($\bar{z}_I$) of between 0 and 100) look to be in the range of 20 to 80 (Figure S5.1), which are used to calibrate the simulations with along with the exposure scaling factor ($\beta$) (see Supplement 2).

We can see the standard deviation varies with the magnitude of resistance in the population (Figure S5.2). This is an important point, especially when considering mixtures, where the two insecticides in the mixture may have substantially different levels of resistance. We therefore conduct a linear model of standard deviation ~ mean PRS (Equation 1e). Values were limited to PRS values less than 3600 (80% bioassay survival). The results of the linear model are in Table S5. For ease of use, we will use the intercept value of 18 and the regression coefficient value for the mean PRS as 0.4.

| **Table S5 Linear Model Mean PRS and Standard Deviation** | | | | |
|---|---|---|---|---|
| | Estimate | Lower 95% CI | Upper 95% CI | p value |
| Intercept | 18.19349 | -1.9884385 | 38.3754220 | 0.0768 |
| Mean PRS | 0.40220 | 0.3822935 | 0.4221146 | <2e-16 |
| Residual standard error: 98.43 on 127 degrees of freedom<br>Multiple R-squared: 0.9264, Adjusted R-squared: 0.9258<br>F-statistic: 1598 on 1 and 127 DF, p-value: < 2.2e-16 | | | | |

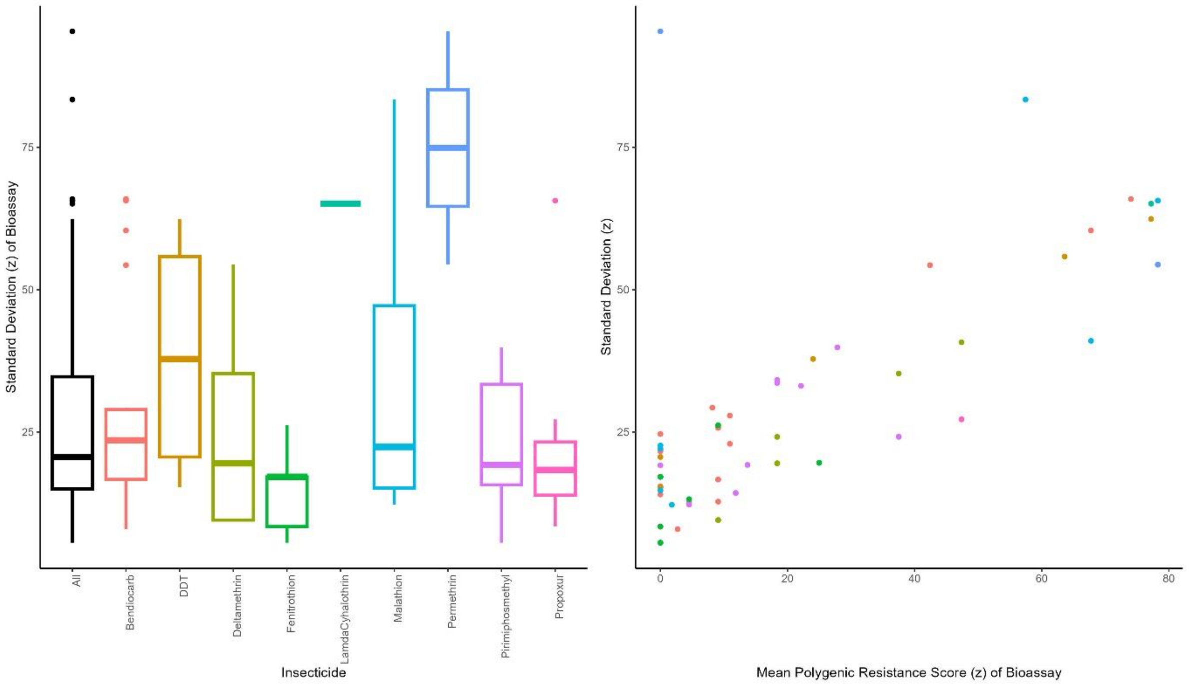


**Figure S5.1: Standard deviation of Polygenic Resistance Score at 10% Bioassay Survival.** Left panel: Boxplot of the standard deviation of the calculated mean polygenic resistance score for all (black) and individual insecticides (colours). Right panel: scatterplot of the relationship between the mean PRS and the corresponding calculated standard deviation of the mean PRS value.

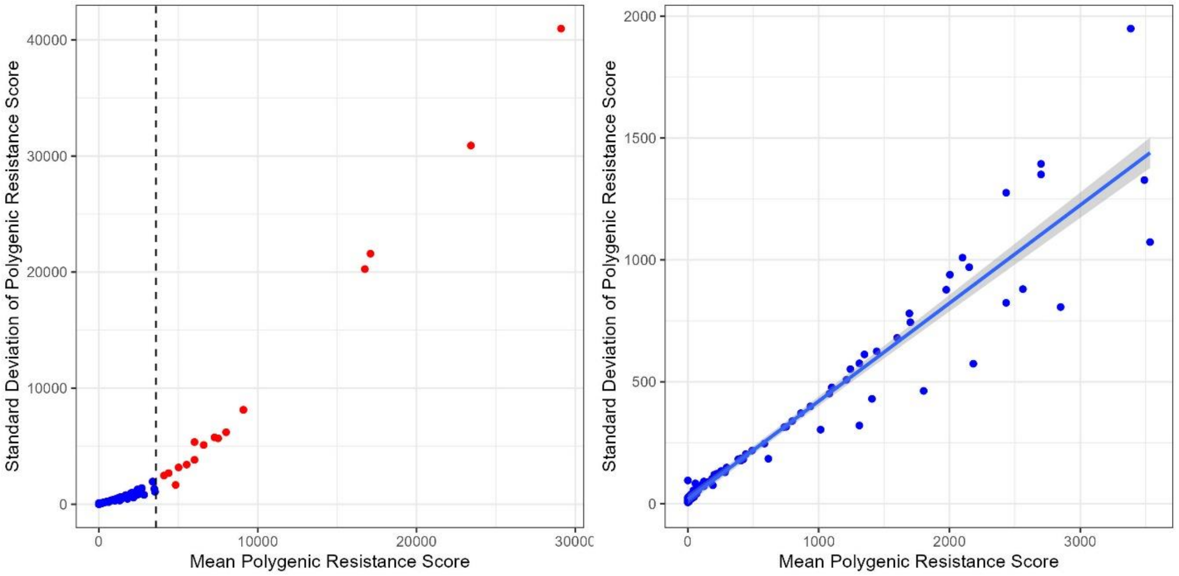


**Figure S5.2 Relationship between Mean Polygenic Resistance Score and Standard Deviation.** Left Panel: Relationship for all PRS values. Right Panel: PRS values limited to 0-3600 (0-80% bioassay survival), with fitted linear model (and 95% CI).

**References**

Alemayehu, E., Asale, A., Eba, K., Getahun, K., Tushune, K., Bryon, A., Morou, E., Vontas, J., Van Leeuwen, T., Duchateau, L., & Yewhalaw, D. (2017). Mapping insecticide resistance and characterization of resistance mechanisms in Anopheles arabiensis (Diptera: Culicidae) in Ethiopia. *Parasites and Vectors*, *10*(1), 1–11. https://doi.org/10.1186/s13071-017-2342-y

Thomsen, E. K., Strode, C., Hemmings, K., Hughes, A. J., Chanda, E., Musapa, M., Kamuliwo, M., Phiri, F. N., Muzia, L., Chanda, J., Kandyata, A., Chirwa, B., Poer, K., Hemingway, J., Wondji, C. S., Ranson, H., & Coleman, M. (2014). Underpinning sustainable vector control through informed insecticide resistance management. *PLoS ONE*, *9*(6). https://doi.org/10.1371/journal.pone.0099822

Wat'Senga, F., Manzambi, E. Z., Lunkula, A., Mulumbu, R., Mampangulu, T., Lobo, N., Hendershot, A., Fornadel, C., Jacob, D., Niang, M., Ntoya, F., Muyembe, T., Likwela, J., Irish, S. R., & Oxborough, R. M. (2018). Nationwide insecticide resistance status and biting behaviour of malaria vector species in the Democratic Republic of Congo. *Malaria Journal*, *17*(1), 1–13. https://doi.org/10.1186/s12936-018-2285-6
