## Supplement 6 for "Mathematical Methodology for Dynamic Models of Insecticide Selection Assuming a Polygenic Basis of Resistance"

### Supplement 6: Fitness cost Selection Differential Values:

In this supplement the two options for implementing fitness costs and provide parameter estimates are detailed.

**Option 1: Fitness selection differentials remain fixed:**

If $\sigma_I$ remains fixed, then it seems intuitive for $S_I^{\phi♀}$ and $S_I^{\phi♂}$ to remain fixed throughout the simulations. This is because the difference in PRS between the most resistant individuals and least resistant remains constant, and therefore the relative fitness difference between the most and least resistant individuals remains constant.

**Option 2: Fitness selection differentials vary with $\sigma_I$ and $\bar{z}_I$:**

If $\sigma_I$ varies with $\bar{z}_I$ (see Supplement 5) the difference in PRS between the most and least resistant individuals in the population to varies. We can therefore allow the fitness costs to be a fixed proportion of the standard deviation, allowing the fitness costs selection differential to increase with an increasing $\bar{z}_I$ and therefore increasing $\sigma_I$.

$$S_I^{\phi} = \phi\sigma_I$$

Equation 6a

As it may be expected for fitness costs differentially affect males and females, we can separately calculate the sex-specific fitness cost selection differentials.

$$S_I^{\phi♀} = \phi^{♀}\sigma_I$$

Equation 6b(♀)

$$S_I^{\phi♂} = \phi^{♂}\sigma_I$$

Equation 6b(♂)

By manually inputting the fitness selection differentials (option 1 or option 2), it may be possible for some simulations to never "take-off" if the fitness cost selection differential is higher than selection differential brought about by insecticide selection ($S_I^{\Phi♀}$ and $S_I^{\Phi♂}$ are greater than $S_I^{S♀}$ and $S_I^{S♂}$). These would be situations where insecticide selection is low (e.g. low coverage and low exposure), and in such situations the selection pressure would be low anyway such that even in the absence of any fitness cost resistance would be expected to be very slow to build up anyway. And of course, in situations where IR cannot take off, there is no need for any IRM as IR does not become a problem.

**Option 1 Parameter Estimation: Fixed Fitness Cost Selection Differential.**

When the standard deviation is constant in the simulation, the fitness cost selection differential is input manually as a fixed value. In the previous model version (polyres), fitness costs were calculated as a proportion of the response. This was performed as the response was constant and to ensure fitness costs were lower than the response (guaranteeing resistance takes off). However, with this dynamic model, the response varies depending on the level of resistance in the population and is therefore not fixed throughout the simulation.

For the "polyres" model the fitness costs response (Equation 5 in Hobbs et al., 2023) was calculated maximally at:

$$fitness\ cost\ response = -0.2\left(10\frac{0.3 * 0.9 * (1+1)}{2}\right) = 0.174$$

and minimally at:

$$fitness\ cost\ response = -0.01\left(10\frac{0.05 * 0.4 * (1+0)}{2}\right) = 0.002$$

Assuming $S_i^{\phi♀} = S_i^{\phi♂}$ and $h_I^2$ is maximally 0.3 and minimally 0.05. These values can be input into Equation 3c. The minimum and maximum fitness cost selection differentials are $S_i^{\phi} = -0.58$ and $S_i^{\phi} = -0.04$.

**Option 2 Parameter Estimation: Fitness selection differentials vary with $\boldsymbol{\sigma_I}$ and $\boldsymbol{\bar{z}_I}$**

When the standard deviation varies, so too will the fitness cost selection differential. $S_I^{\phi}$ is maximally and minimally at $-0.58$ and $-0.04$, for a fixed standard deviation. That these would be the selection differentials when $\bar{z}_I$=0. For when $\bar{z}_I$=0, then $\sigma_I$=18 (Equation 1e). Therefore, using Equation 6a, ϕ is maximally 0.0322 and minimally 0.0022 when allowing the fitness costs to vary with the magnitude of the mean resistance of the population.
