## Supplement 7 for "Mathematical Methodology for Dynamic Models of Insecticide Selection Assuming a Polygenic Basis of Resistance"

### Supplement 7: Updating the Male Equations to allow for different insecticide coverages and encounter probabilities.

We must account for male mosquitoes also being able to contact complex insecticide deployments such as combinations or micro-mosaics, which will impact the level of selection on male mosquitoes and therefore impact the male insecticide selection differential. This involves updating Equation 4b(♂), to allow for male mosquitoes to encounter just insecticide $i$, just insecticide $j$ or both insecticide $i$ and $j$. The mean PRS of the male mosquitoes forming the breeding parental population ($\bar{z}_I^{P♂}$) is calculated from all insecticide encounters, where the superscript "$P$" indicates the parental population. The superscript "$E$" indicates the mosquitoes survived their encounter with the insecticide(s) and the superscript "$u$" indicates those males did not encounter the insecticide.

$$\bar{z}_I^{P♂} = \frac{\left(\left(N_i^{E♂}\bar{z}_I^{E♂}\right) + \left(N_j^{E♂}\bar{z}_I^{♂}\right) + \left(N_{ij}^{E♂}\bar{z}_I^{E♂}\right) + \left(N^{u♂}\,\bar{z}_I^{♂}\right)\right)}{N^{P♂}}$$

Equation 12a(♂)

The total number of males is then:

$$N^{P♂} = N_i^{E♂} + N_j^{E♂} + N_{ij}^{E♂} + N^{u♂}$$

Equation 12b(♂)

The number of males surviving the encounter only with insecticide $i$ ($N_i^{E♂}$). This is first those entering houses with insecticide $i$ only and surviving. Second, those entering houses with both insecticide $i$ and $j$ but contacting only insecticide $i$ and surviving.

$$N_i^{E♂} = \left(c_i x m \bar{K}_i^F N^{T♂}\right) + \left(c_{ij}\Lambda_{i|ij}^{♂} x m \bar{K}_i^F N^{T♂}\right)$$

Equation 12c(♂)

Similarly for encountering only insecticide $j$:

$$N_j^{E♂} = \left(c_j xm\bar{K}_j^F N^{T♂}\right) + \left(c_{ij}\Lambda_{j|ij}^{♂} xm\bar{K}_j^F N^{T♂}\right)$$

Equation 12d(♂)

The number of males surviving the encounter with both insecticides is the number of males entering a house with both insecticides, encountering both insecticides, and surviving both insecticides.

$$N_{ij}^{E♂} = c_{ij}\Lambda_{ij|ij}^{♂} xm\bar{K}_i^F \bar{K}_j^F N^{T♂}$$

Equation 12e(♂)

The number of males not encountering insecticides:

$$N^{u♂} = N^{T♂}(1 - xm)$$

Equation 12f(♂)

The value of $F_{z_I^{E♂}}$ is calculated for those encountering insecticide $i$:

$$F_{z_I^{E♂}} = \left(F_{z_I^{♂}} K_i^F xmc_i\right) + \left(F_{z_I^{♂}} c_{ij}\Lambda_{i|ij}^{♂} xmK_i^F\right) + \left(F_{z_I^{♂}} c_{ij}\Lambda_{ij|ij}^{♂} xmK_i^F \bar{K}_j^F\right)$$

Equation 12g(♂)

The values of $F_{z_I^{E♂}}$ can then be used to calculate the mean PRS of the exposed surviving males:

$$\bar{z}_I^{E♂} = \left(\sum_{z_I^{♂}=-\infty}^{\infty} F_{z_I^{E♂}}\, z_I^{♂}\right) / \left(N_i^{E♂} + N_{ij}^{E♂}\right)$$

Equation 12h(♂)

$\bar{z}_I^{E♂}$ is returned to Equation 12a(♂) to calculate $\bar{z}_I^{P♂}$ which is required to calculate the male insecticide selection differential (Equation 4a(♂)) which can be combined with fitness costs to finally be incorporated into the sex-specific Breeder's equation allowing for multiple gonotrophic cycles (Equation 11b) to allow for the calculation of the response.
