## Supplement 8 for "Mathematical Methodology for Dynamic Models of Insecticide Selection Assuming a Polygenic Basis of Resistance"

### Supplement 8: Allowing for dispersal in the multiple gonotrophic cycle model.

A conceptual description of the multiple gonotrophic cycle model with dispersal is in Figure S8.1. A notation a guide is in Figure S8.2. As female mosquitoes mate once and non-overlapping generations are assumed, male dispersal is not tracked.

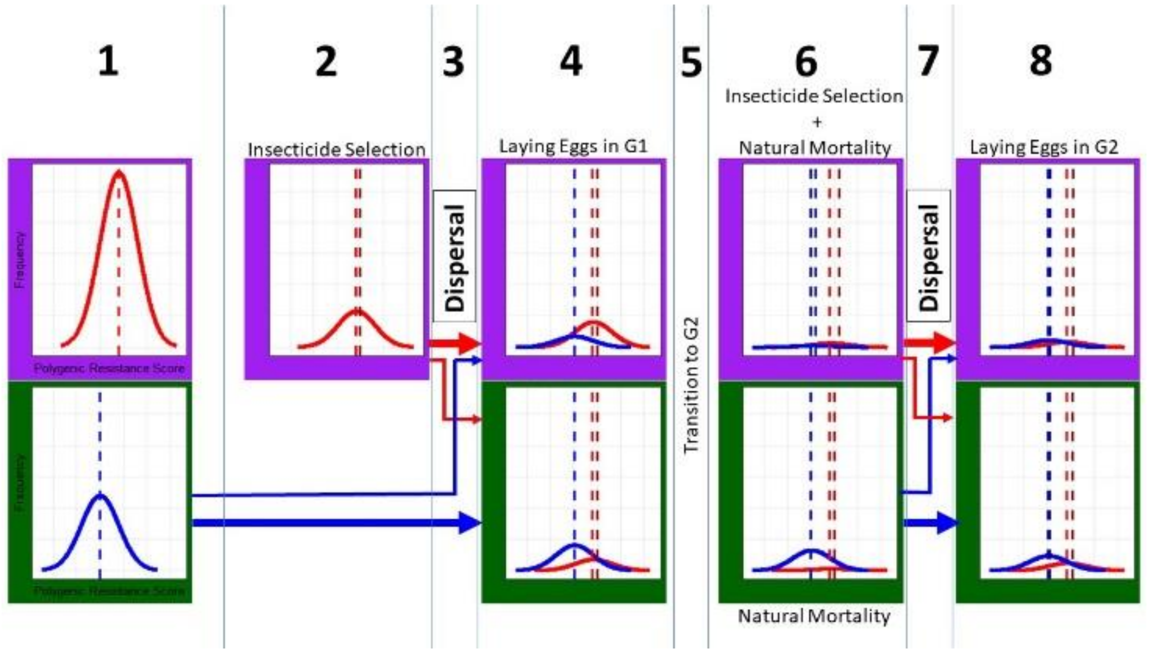


**Figure S8.1 Diagrammatic Representation of Multiple Dispersal Events with Multiple Gonotrophic Cycles.** Left to Right: **1):** The mosquito populations emerge in the intervention site (purple background) and the refugia (green background). **2):** Selection occurs. Insecticide selection and fitness costs occur in the intervention site. Fitness costs occur in the refugia. After insecticide selection the mosquitoes mate. Males and females which emerged in the intervention site mate with one another. Males and females which emerged in the refugia mate with each other. **3):** Female mosquitoes are then allowed to disperse between the two sites. **4):** Laying eggs in the intervention site (purple background) are therefore the remaining females from the intervention site (red) and the females joining from the refugia (blue). And laying eggs in the refugia (green background) is therefore the females remaining (blue) and those joining from the intervention site (red). **5):** The second gonotrophic cycle is then allowed to start. **6):** In the intervention site (purple background), insecticide selection and natural mortality is applied to the females there (both the “intervention” females (red) and the “refugia” females (blue)). In the refugia (green background) natural mortality is applied to the females there (both the “refugia” females (blue) and the “intervention” females (red)). **7):** The female mosquitoes are then allowed to disperse once against between the two sites. **8):** The female mosquitoes then lay eggs in the sites in to which they are in. After which the third, fourth, fifth gonotrophic cycles start following the same process until the maximum number of gonotrophic cycles is reached. The vertical lines are: light coloured = initial mean PRS at emergence, and the darker line = current mean PRS in the current location. The difference is therefore the female insecticide selection differential.

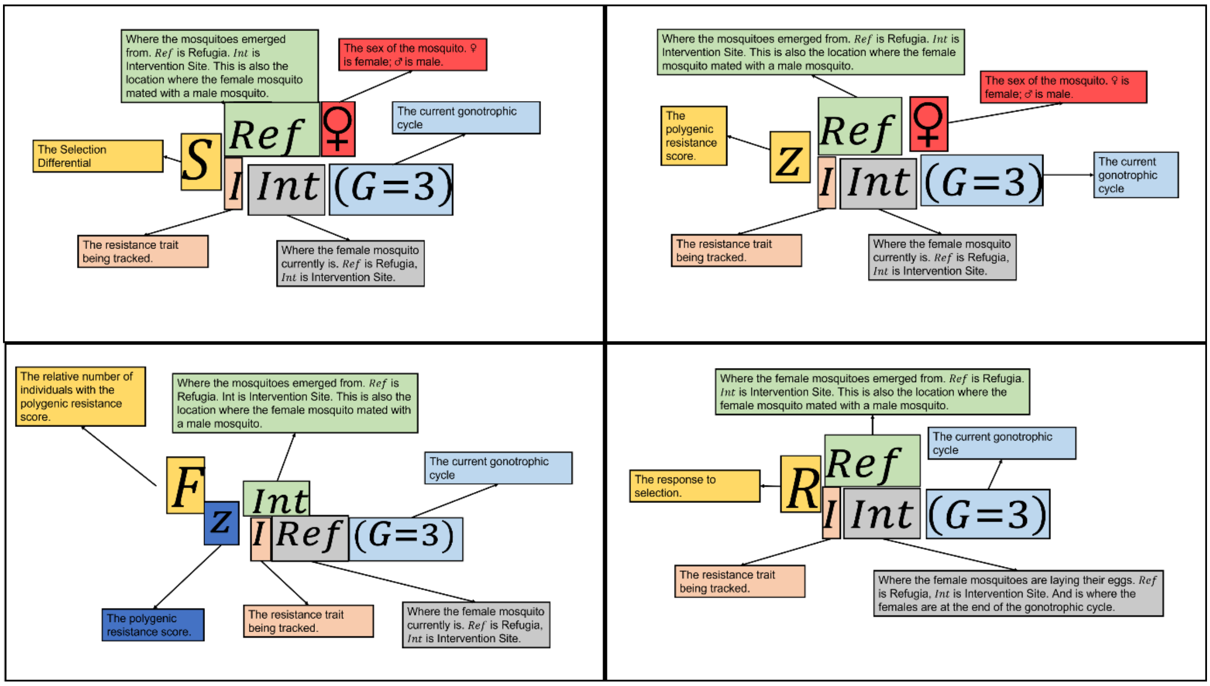


**Figure S8.2 Reading the Mathematical Symbols Used for Multiple Gonotrophic Cycles with Dispersal.** Superscripts refer to the original location of the mosquito, where the mosquito emerged. For females this also indicates where they mated. Subscripts refer to the current location of the mosquito, where is the mosquito having a selection pressure applied or is laying eggs.

#### Equations for Gonotrophic cycle 1: Laying Eggs in the Refugia

The frequency of females with a PRS of $z_I^{♀}$ laying eggs in the refugia in the first gonotrophic cycle is those who hatch and mate in the refugia and stay in the refugia ($F_{z_{I\,Ref\,(G=1)}^{Ref\,♀}}$) and those who hatching and mating in the intervention site (having insecticide selection) migrating to the refugia ($F_{z_{I\,Ref\,(G=1)}^{Int\,♀}}$):

$$F_{z_{I\,Ref\,(G=1)}^{Ref\,♀}} = \left(F_{z_{I\,Ref(G=0)}^{Ref\,♀}} * (1-C)\right) * (1-\theta)$$

Equation 14a(i)(Ref)

$$F_{z_{I\,Ref\,(G=1)}^{Int\,\text{♀}}} = \left(\left(F_{z_{I\,Int\,(G=0)}^{Int\,\text{♀}}} Cc_i x K_i^F\right) + \left(F_{z_{I\,Int\,(G=0)}^{Int\,\text{♀}}} Cc_j x\, \bar{K}_j^F\right) + \left(F_{z_{I\,Int\,(G=0)}^{Int\,\text{♀}}} Cc_{ij} \Lambda_{j|ij}^{\text{♀}} x\, \bar{K}_j^F\right)\right.$$

$$+ \left(F_{z_{I\,Int\,(G=0)}^{Int\,\text{♀}}} Cc_{ij} \Lambda_{i|ij}^{\text{♀}} x K_i^F\right) + \left(F_{z_{I\,Int\,(G=0)}^{Int\,\text{♀}}} Cc_{ij} \Lambda_{ij|ij}^{\text{♀}} x K_i^F \bar{K}_j^F\right)$$

$$\left. + \left(F_{z_{I\,Int\,(G=\,0)}^{Int\,\text{♀}}} C * (1 - x)\right)\right) * \ \theta$$

Equation 14b(i) (Ref)

The term $C$ scales the intervention and refugia population sizes by the coverage and is included only in the first gonotrophic cycle. $F_{z_{I\ (G=0)}^{Ref\,\text{♀}}}$ and $F_{z_{I\,Int\,(G=\,0)}^{Int\,\text{♀}}}$ are calculated from Equation 4g(♀). $K_i^F$ depends on the efficacy of insecticide $i$. $\bar{K}_j^F$ will depends on the mean resistance ($\bar{z}_{J\,Int\,(G=0)}^{Int\,\text{♀}}$) and efficacy of insecticide $j$ (Equation 2b(i)).

Laying eggs in the refugia in each gonotrophic cycle are from a total number of females who originally hatched and mated in the refugia ($N_{Ref\,(G)}^{Ref\,\text{♀}}$) and in the intervention site ($N_{Ref\,(G)}^{Int\,\text{♀}}$):

$$N_{Ref\,(G)}^{Ref\,\text{♀}} = \sum_{z_{I\,Ref}^{Ref\,\text{♀}} = -\infty}^{\infty} F_{z_{I\,Ref\,(G)}^{Ref\,\text{♀}}}$$

Equation 14c(i) (Ref)

$$N_{Ref\,(G)}^{Int\,\text{♀}} = \sum_{z_{I\,Ref}^{Int\,\text{♀}} = -\infty}^{\infty} F_{z_{I\,Ref\,(G)}^{Int\,\text{♀}}}$$

Equation 14d(i) (Ref)

The mean PRS for each gonotrophic cycle which originally emerged in the refugia:

$$\bar{z}^{P\,Ref\,\text{♀}}_{I\,Ref\,(G)} = \left(\sum_{z^{Ref\text{♀}}_{I\,Ref\,(G)}=-\infty}^{\infty} F_{z^{Ref\text{♀}}_{I\,Ref\,(G)}}\, z^{Ref\text{♀}}_{I\,(G=0)}\right) / N^{Ref\,\text{♀}}_{Ref\,(G)}$$

Equation 14e(i) (Ref)

And originally emerged in the intervention site:

$$\bar{z}^{P\,Int\,\text{♀}}_{I\,Ref\,(G)} = \left(\sum_{z^{Int\,\text{♀}}_{I\,Ref\,(G)}=-\infty}^{\infty} F_{z^{Int\text{♀}}_{I\,Ref\,(G)}}\, z^{Int\text{♀}}_{I\,(G=0)}\right) / N^{Int\text{♀}}_{Ref\,(G)}$$

Equation 14f(i) (Ref)

Female insecticide selection differentials in the refugia can be written for any gonotrophic cycle as:

$$S^{SRef\,\text{♀}}_{I\,Ref\,(G)} = \bar{z}^{PRef\text{♀}}_{I\,Ref\,(G)} - \bar{z}^{Ref\,\text{♀}}_{I\,(G=0)}$$

Equation 14g(i) (Ref)

$$S^{S\,Int\,\text{♀}}_{I\,Ref\,(G)} = \bar{z}^{P\,Int\,\text{♀}}_{I\,Ref\,(G)} - \bar{z}^{Int\text{♀}}_{I\,(G=0)}$$

Equation 14h(i) (Ref)

This is combined with the fitness cost selection differential (Equation 11a(ii)) to give the overall selection differential. The responses are then calculated, for the females which hatched and mated in the refugia ($R^{Ref}_{I\,Ref\,(G)}$) and females which hatched and mated in the intervention site ($R^{Int}_{I\,Ref\,(G)}$):

$$R^{Ref}_{I\,Ref\,(G)} = h^2\left(\frac{S^{Ref\Phi\text{♀}}_{I\,Ref\,(G)} + S^{Ref\Phi\text{♂}}_{I}}{2}\right)\beta$$

Equation 14i(i) (Ref)

$$R^{Int}_{I\,Ref\,(G)} = h^2\left(\frac{S^{Int\,S\Phi\text{♀}}_{I\,Ref\,(G)} + S^{Int\,S\Phi\,\text{♂}}_{I}}{2}\right)\beta$$

Equation 14j(i) (Ref)

**Gonotrophic cycle 1: Laying Eggs in the Intervention**

A parallel process occurs for the intervention site. The frequency of females laying eggs in the intervention site in the first gonotrophic cycle is those hatching and mating in the refugia migrating to the intervention site ($F_{z^{Ref\text{♀}}_{I\,Int(G=1)}}$) and those hatching and mating in the intervention site (having insecticide selection) staying in the intervention site ($F_{z^{Int\text{♀}}_{I\,Int(G=1)}}$):

$$F_{z^{Ref\text{♀}}_{I\,Int(G=1)}} = F_{z^{Ref\text{♀}}_{I\,Ref(G=0)}}\theta * (1 - C)$$

Equation 14a(i) (Int)

$$\begin{aligned} F_{z^{Int\text{♀}}_{I\,Int(G=1)}} = \Bigg(&\left(F_{z^{Int\,\text{♀}}_{I\,Int(G=0)}} Cc_i x\, K^F_i\right) + \left(F_{z^{Int\,\text{♀}}_{I\,Int(G=0)}} C\, c_j x \bar{K}^F_j\right) + \left(F_{z^{Int\,\text{♀}}_{I\,Int(G=0)}} C\, c_{ij}\Lambda^{\text{♀}}_{i|ij} x K^F_i\right) \\ &+ \left(F_{z^{Int\,\text{♀}}_{I\,Int(G=0)}} C\, c_{ij}\Lambda^{\text{♀}}_{j|ij} x \bar{K}^F_j\right) + \left(F_{z^{Int\,\text{♀}}_{I\,Int(G=0)}} C\, c_{ij}\Lambda^{\text{♀}}_{ij|ij} x K^F_i \bar{K}^F_j\right) \\ &+ \left(F_{z^{Int\,\text{♀}}_{I\,Int(G=0)}} C * (1 - x)\right)\Bigg) * (1 - \theta) \end{aligned}$$

Equation 14b(i) (Int)

Laying eggs in the intervention site in each gonotrophic cycle are from a total number of females who originally hatched and mated in the refugia ($N^{Ref\,\text{♀}}_{Int\,(G)}$) and in the intervention site ($N^{Int\,\text{♀}}_{Int\,(G)}$):

$$N^{Ref\,\text{♀}}_{Int\,(G)} = \sum_{z^{Ref\text{♀}}_{I\,Int\,(G)} = -\infty}^{\infty} F_{z^{Ref\text{♀}}_{I\,Int\,(G)}}$$

Equation 14c(i) (Int)

$$N_{Int\,(G)}^{Int\,♀} = \sum_{z_{I\,Int\,(G)}^{Int\,♀} = -\infty}^{\infty} F_{z_{I\,Int\,(G)}^{Int\,♀}}$$

Equation 14d(i) (Int)

The updated mean PRS for each gonotrophic cycle which originally emerged in the refugia is:

$$\bar{z}_{I\,Int\,(G)}^{P\,Ref\,♀} = \left( \sum_{z_{I\,Int\,(G)}^{Ref\,♀} = -\infty}^{\infty} F_{z_{I\,Int\,(G)}^{Ref♀}} \, z_{I\,(G=0)}^{Ref♀} \right) / N_{Int\,(G)}^{Ref\,♀}$$

Equation 14e(i) (Int)

And for those originally emerged in the intervention site:

$$\bar{z}_{I\,Int\,(G)}^{P\,Int\,♀} = \left( \sum_{z_{I\,Int\,(G)}^{Int\,♀} = -\infty}^{\infty} F_{z_{I\,Int\,(G)}^{Int\,♀}} \, z_{I\,(G=0)}^{Int♀} \right) / N_{Int\,(G)}^{Int\,♀}$$

Equation 14f(i) (Int)

Female insecticide selection differentials in the intervention site for any gonotrophic cycle are calculated as:

$$S_{I\,Int\,(G)}^{S\,Ref\,♀} = \bar{z}_{I\,Int\,(G)}^{S\,Ref\,♀} - \bar{z}_{I\,(G=0)}^{Ref\,♀}$$

Equation 14g(i) (Int)

$$S_{I\,Int\,(G)}^{S\,Int\,♀} = \bar{z}_{I\,Int\,(G)}^{S\,Int\,♀} - \bar{z}_{I\,(G=0)}^{Int♀}$$

Equation 14h(i) (Int)

Fitness costs are implemented using Equation 11a(ii) giving the overall selection differential for a gonotrophic cycle. Responses are calculated, for the females which hatched and mated in the refugia ($R^{Ref}_{I\,Int\,(G)}$) and females which hatched and mated in the intervention site ($R^{Int}_{I\,Int\,(G)}$):

$$R^{Ref}_{I\,Int\,(G)} = h^2\left(\frac{S^{Ref\,\Phi\text{♀}}_{I\,Int\,(G)} + S^{Ref\,\Phi\,\text{♂}}_{I}}{2}\right)\beta$$

Equation 14i(i) (Int)

$$R^{Int}_{I\,Int\,(G)} = h^2\left(\frac{S^{Int\,S\Phi\text{♀}}_{I\,Int\,(G)} + S^{Int\,S\Phi\,\text{♂}}_{I}}{2}\right)\beta$$

Equation 14j(i) (Int)

**Gonotrophic Cycle 2 and Beyond: Laying Eggs in the Refugia**

For subsequent gonotrophic cycles, the frequency of females laying eggs in the refugia is: Those emerged and mated in the refugia, who either are remain in the refugia or join from the intervention site (having undergone insecticide selection in this gonotrophic cycle):

$$F_{z^{Ref\text{♀}}_{I\,Ref\,(G)}} = \left(F_{z^{Ref\text{♀}}_{I\,Ref\,(G-1)}}(1-\theta)\rho\right)$$
$$+\left(\left(\left(F_{z^{Ref\text{♀}}_{I\,Int\,(G-1)}}(1-x)\right)+\left(F_{z^{Ref\text{♀}}_{I\,Int\,(G-1)}}c_i x K^F_i\right)+\left(F_{z^{Ref\text{♀}}_{I\,Int\,(G-1)}}c_j x \overline{K}^F_j\right)\right.\right.$$
$$+\left(F_{z^{Ref\text{♀}}_{I\,Int\,(G-1)}}c_{ij}\Lambda^{\text{♀}}_{i|ij}xK^F_i\right)+\left(F_{z^{Ref\text{♀}}_{I\,Int\,(G-1)}}c_{ij}\Lambda^{\text{♀}}_{j|ij}x\overline{K}^F_j\right)$$
$$\left.\left.+\left(F_{z^{Ref\text{♀}}_{I\,Int\,(G-1)}}c_{ij}\Lambda^{\text{♀}}_{ij|ij}xK^F_i\overline{K}^F_j\right)\right)*\theta\rho\right)$$

Equation 14a(ii)(Ref)

Those emerged and mated in the intervention site, who previously joined the refugia (and stay again) or those newly joining from the intervention site (having undergone insecticide selection):

$$F_{z_{I\,Ref\,(G)}^{Int\,\text{♀}}} = \left(F_{z_{I\,Ref\,(G-1)}^{Int\text{♀}}}(1-x)(1-\theta)\rho\right)$$

$$+\left(\left(F_{z_{I\,Int\,(G-1)}^{Int\,\text{♀}}}c_i x K_i^F\right)+\left(F_{z_{I\,Int\,(G-1)}^{Int\,\text{♀}}}c_j x \overline{K}_j^F\right)+\left(F_{z_{I\,Int\,(G-1)}^{Int\,\text{♀}}}c_{ij}\Lambda_{i|ij}^{\text{♀}} x K_i^F\right)\right.$$

$$+\left(F_{z_{I\,Int\,(G-1)}^{Int\,\text{♀}}}c_{ij}\Lambda_{j|ij}^{\text{♀}} x \overline{K}_j^F\right)+\left(F_{z_{I\,Int\,(G-1)}^{Int\,\text{♀}}}c_{ij}\Lambda_{ij|ij}^{\text{♀}} x K_i^F\,\overline{K}_j^F\right)$$

$$\left.+\left(F_{z_{I\,Int\,(G-1)}^{Int\,\text{♀}}}(1-x)\right)*\theta\rho\right)$$

Equation 14b(ii) (Ref)

$F_{z_{I\,Ref\,(G)}^{Ref\text{♀}}}$ is transferred to Equations 14c(i)(Ref) and 14e(i)(Ref) and $F_{z_{I\,Ref\,(G)}^{Int\,\text{♀}}}$ is transferred to Equations 14d(i)(Ref) and 14f(i)(Ref).

**Gonotrophic Cycle 2 and Beyond: Laying Eggs in the Intervention Site**

A parallel process occurs in the intervention site. The frequency of females laying eggs in the intervention site is: Those emerged and mated in the refugia (and joined the intervention site previously), who remain in the intervention site (undergoing insecticide selection) or are joining from the refugia:

$$F_{z_{I\,Int\,(G)}^{Ref\text{♀}}} = \left(F_{z_{I\,Ref\,(G-1)}^{Ref\text{♀}}}\theta\rho\right)$$

$$+\left(\left(\left(F_{z_{I\,Int(G-1)}^{Ref\text{♀}}}x c_i K_i^F\right)+\left(F_{z_{I\,Int(G-1)}^{Ref\text{♀}}}x c_j \overline{K}_j^F\right)+\left(F_{z_{I\,Int(G-1)}^{Ref\text{♀}}}x c_{ij}\Lambda_{i|ij}^{\text{♀}} K_i^F\right)\right.\right.$$

$$+\left(F_{z_{I\,Int(G-1)}^{Ref\text{♀}}}x c_{ij}\Lambda_{j|ij}^{\text{♀}} \overline{K}_j^F\right)+\left(F_{z_{I\,Int(G-1)}^{Ref\text{♀}}}x c_{ij}\Lambda_{ij|ij}^{\text{♀}} K_i^F\,\overline{K}_j^F\right)$$

$$\left.\left.+\left(F_{z_{I\,Int(G-1)}^{Ref\text{♀}}}(1-x)\right)\right)*(1-\theta)*\rho\right)$$

Equation 14a(ii) (Int)

Females who originally emerged and mated in the intervention site, who previously joined the refugia and now return to the intervention site and those who remained in the intervention site (and undergo insecticide selection):

$$F_{z_{I\,Int\,(G)}^{Int\,\text{♀}}} = \left(F_{z_{I\,Ref\,(G-1)}^{Int\,\text{♀}}}\theta\rho\right)$$

$$+\left(\left(\left(\left(F_{z_{I\,Int(G-1)}^{Int\text{♀}}} xc_i K_i^F\right) + \left(F_{z_{I\,Int(G-1)}^{Int\text{♀}}} xc_j \overline{K}_j^F\right) + \left(F_{z_{I\,Int(G-1)}^{Int\text{♀}}} xc_{ij}\Lambda_{i|ij}^{\text{♀}} K_i^F\right)\right.\right.\right.$$

$$+\left(F_{z_{I\,Int(G-1)}^{Int\text{♀}}} xc_{ij}\Lambda_{j|ij}^{\text{♀}}\overline{K}_j^F\right) + \left(F_{z_{I\,Int(G-1)}^{Int\text{♀}}} xc_{ij}\Lambda_{ij|ij}^{\text{♀}} K_i^F\overline{K}_j^F\right)$$

$$\left.\left.\left.+\left(F_{z_{I\,Int(G-1)}^{Int\text{♀}}}(1-x)\right)\right)\right) * (1-\theta) * \rho\right)$$

Equation 14b(ii) (Int)

$F_{z_{I\,Int\,(G)}^{Ref\text{♀}}}$ is transferred to Equation 14c(i) (Int) and 14e(i) (Int), and $F_{z_{I\,Int\,(G)}^{Int\,\text{♀}}}$ is transferred to equations 14d(i) (Int) and 14f(i) (Int).

**Calculating the Overall Selection Response for Multiple Gonotrophic Cycles with Dispersal**

Responses need to be weighted by the number of eggs laid in each gonotrophic cycle. The total number of oviposition events (and therefore proportional to the total number of eggs laid) in the Refugia is therefore:

$$N_{o\,Ref}^{Total\text{♀}} = N_{o\,Ref}^{Ref\text{♀}} + N_{o\,Ref}^{Int\text{♀}}$$

Equation 14k(Ref)

$N_{o\,Ref}^{Ref\text{♀}}$ is the total contribution from females born (and mated) in the refugia:

$$N_{o\,Ref}^{Ref\text{♀}} = \sum_{G=1}^{G_{max}} N_{Ref\,(G)}^{Ref\text{♀}}$$

Equation 14l(Ref)

$N_{o\,Ref}^{Int♀}$ is the total contribution from females born (and mated) in the intervention site:

$$N_{o\,Ref}^{Int♀} = \sum_{G=1}^{G_{max}} N_{Ref\,(G)}^{Int\,♀}$$

Equation 14m(Ref)

And correspondingly the total number of oviposition events in the intervention site:

$$N_{o\,Int}^{Total♀} = N_{o\,Int}^{Int♀} + N_{o\,Int}^{Ref\,♀}$$

Equation 14k(Int)

$N_{o\,Int}^{Int♀}$ is the total contribution from females born (and mated) in the intervention site:

$$N_{o\,Int}^{Int♀} = N_{Int(G=1)}^{Int♀} + N_{Int(G=2)}^{Int♀} + N_{Int(G=3)}^{Int♀} \dots$$

Equation 14(Int)

$N_{o\,Int}^{Ref\,♀}$ is the total contribution from females born (and mated) in the refugia:

$$N_{o\,Int}^{Ref\,♀} = N_{Int(G=1)}^{Ref\,♀} + N_{Int(G=2)}^{Ref\,♀} + N_{Int(G=3)}^{Ref\,♀} \dots$$

Equation 14m(Int)

The responses for each gonotrophic cycle are weighted to calculate the overall responses. In the intervention site this is for eggs laid by females originally emerged (and mated) in the intervention site:

$$R_{I\,Int}^{T\,Int} = \sum_{G=1}^{G_{max}} \left(R_{I\,Int(G)}^{Int} + (\alpha_{JI} R_{Int\,(G)}^{Int})\right) \frac{N_{Int(G)}^{Int♀}}{N_{o\,Int}^{Int♀}}$$

Equation 14n(Int)

And for females originally emerged (and mated) in the refugia:

$$R^{T\,Ref}_{I\,Int} = \sum_{G=1}^{G_{max}} \left(R^{Ref}_{I\,Int(G)} + (\alpha_{JI} R^{Ref}_{Int\,(G)})\right) \frac{N^{Ref\text{♀}}_{Int(G)}}{N^{Ref\text{♀}}_{o\,Int}}$$

Equation 14o(Int)

The mean of the eggs (the next generation) in the intervention site is:

$$\bar{z}^{Int''}_{I} = \frac{\left(N^{Int\text{♀}}_{o\,Int}(\bar{z}^{Int}_{I} + R^{T\,Int}_{I\,Int})\right) + \left(N^{Ref\text{♀}}_{o\,Int}\left(\bar{z}^{Ref}_{I} + R^{T\,Ref}_{I\,Int}\right)\right)}{N^{Total\text{♀}}_{o\,Int}}$$

Equation 14p(Int)

In the refugia this is as follows: Weighting the responses in the refugia is for eggs laid by females originally emerged (and mated) in the refugia (Equation 15n(Ref))

$$R^{T\,Ref}_{I\,Ref} = \sum_{G=1}^{G_{max}} \left(R^{Ref}_{I\,Ref(G)} + (\alpha_{JI} R^{Ref}_{Ref\,(G)})\right) \frac{N^{Ref\text{♀}}_{Ref(G)}}{N^{Ref\text{♀}}_{o\,Ref}}$$

Equation 14n(Ref)

And for females originally emerged (and mated) in the intervention site:

$$R^{T\,Int}_{I\,Ref} = \sum_{G=1}^{G_{max}} \left(R^{Int}_{I\,Ref(G)} + (\alpha_{JI} R^{Int}_{Ref\,(G)})\right) \frac{N^{Int\text{♀}}_{Ref(G)}}{N^{Int\text{♀}}_{o\,Ref}}$$

Equation 14o(Ref)

The mean of the eggs (the next generation) in the refugia is:

$$\bar{z}_I^{Ref''} = \frac{\left(N_{o\,Ref}^{Ref\text{♀}}\left(\bar{z}_I^{Ref} + R_{I\,Ref}^{T\,Ref}\right)\right) + \left(N_{o\,Ref}^{Int\text{♀}}\left(\bar{z}_I^{Int} + R_{I\,Ref}^{T\,Int}\right)\right)}{N_{o\,Ref}^{Total\text{♀}}}$$

Equation 14p(Ref)
